# Sequencing of 53,831 diverse genomes from the NHLBI TOPMed Program

**DOI:** 10.1101/563866

**Authors:** Daniel Taliun, Daniel N. Harris, Michael D. Kessler, Jedidiah Carlson, Zachary A. Szpiech, Raul Torres, Sarah A. Gagliano Taliun, André Corvelo, Stephanie M. Gogarten, Hyun Min Kang, Achilleas N. Pitsillides, Jonathon LeFaive, Seung-been Lee, Xiaowen Tian, Brian L. Browning, Sayantan Das, Anne-Katrin Emde, Wayne E. Clarke, Douglas P. Loesch, Amol C. Shetty, Thomas W. Blackwell, Quenna Wong, François Aguet, Christine Albert, Alvaro Alonso, Kristin G. Ardlie, Stella Aslibekyan, Paul L. Auer, John Barnard, R. Graham Barr, Lewis C. Becker, Rebecca L. Beer, Emelia J. Benjamin, Lawrence F. Bielak, John Blangero, Michael Boehnke, Donald W. Bowden, Jennifer A. Brody, Esteban G. Burchard, Brian E. Cade, James F. Casella, Brandon Chalazan, Yii-Der Ida Chen, Michael H. Cho, Seung Hoan Choi, Mina K. Chung, Clary B. Clish, Adolfo Correa, Joanne E. Curran, Brian Custer, Dawood Darbar, Michelle Daya, Mariza de Andrade, Dawn L. DeMeo, Susan K. Dutcher, Patrick T. Ellinor, Leslie S. Emery, Diane Fatkin, Lukas Forer, Myriam Fornage, Nora Franceschini, Christian Fuchsberger, Stephanie M. Fullerton, Soren Germer, Mark T. Gladwin, Daniel J. Gottlieb, Xiuqing Guo, Michael E. Hall, Jiang He, Nancy L. Heard-Costa, Susan R. Heckbert, Marguerite R. Irvin, Jill M. Johnsen, Andrew D. Johnson, Sharon L.R. Kardia, Tanika Kelly, Shannon Kelly, Eimear E. Kenny, Douglas P. Kiel, Robert Klemmer, Barbara A. Konkle, Charles Kooperberg, Anna Köttgen, Leslie A. Lange, Jessica Lasky-Su, Daniel Levy, Xihong Lin, Keng-Han Lin, Chunyu Liu, Ruth J.F. Loos, Lori Garman, Robert Gerszten, Steven A. Lubitz, Kathryn L. Lunetta, Angel C.Y. Mak, Ani Manichaikul, Alisa K. Manning, Rasika A. Mathias, David D. McManus, Stephen T. McGarvey, James B. Meigs, Deborah A. Meyers, Julie L. Mikulla, Mollie A. Minear, Braxton Mitchell, Sanghamitra Mohanty, May E. Montasser, Courtney Montgomery, Alanna C. Morrison, Joanne M. Murabito, Andrea Natale, Pradeep Natarajan, Sarah C. Nelson, Kari E. North, Jeffrey R. O’Connell, Nicholette D. Palmer, Nathan Pankratz, Gina M. Peloso, Patricia A. Peyser, Wendy S. Post, Bruce M. Psaty, D.C. Rao, Susan Redline, Alexander P. Reiner, Dan Roden, Jerome I. Rotter, Ingo Ruczinski, Chloé Sarnowski, Sebastian Schoenherr, Jeong-Sun Seo, Sudha Seshadri, Vivien A. Sheehan, M. Benjamin Shoemaker, Albert V. Smith, Nicholas L. Smith, Jennifer A. Smith, Nona Sotoodehnia, Adrienne M. Stilp, Weihong Tang, Kent D. Taylor, Marilyn Telen, Timothy A. Thornton, Russell P. Tracy, David J. Van Den Berg, Ramachandran S. Vasan, Karine A. Viaud-Martinez, Scott Vrieze, Daniel E Weeks, Bruce S. Weir, Scott T. Weiss, Lu-Chen Weng, Cristen J. Willer, Yingze Zhang, Xutong Zhao, Donna K. Arnett, Allison E. Ashley-Koch, Kathleen C. Barnes, Eric Boerwinkle, Stacey Gabriel, Richard Gibbs, Kenneth M. Rice, Stephen S. Rich, Edwin Silverman, Pankaj Qasba, Weiniu Gan, Trans-Omics for Precision Medicine (TOPMed) Program, TOPMed Population Genetics Working Group, George J. Papanicolaou, Deborah A. Nickerson, Sharon R. Browning, Michael C. Zody, Sebastian Zöllner, James G. Wilson, L Adrienne Cupples, Cathy C. Laurie, Cashell E. Jaquish, Ryan D. Hernandez, Timothy D. O’Connor, Gonçalo R. Abecasis

## Abstract

The Trans-Omics for Precision Medicine (TOPMed) program seeks to elucidate the genetic architecture and disease biology of heart, lung, blood, and sleep disorders, with the ultimate goal of improving diagnosis, treatment, and prevention. The initial phases of the program focus on whole genome sequencing of individuals with rich phenotypic data and diverse backgrounds. Here, we describe TOPMed goals and design as well as resources and early insights from the sequence data. The resources include a variant browser, a genotype imputation panel, and sharing of genomic and phenotypic data via dbGaP. In 53,581 TOPMed samples, >400 million single-nucleotide and insertion/deletion variants were detected by alignment with the reference genome. Additional novel variants are detectable through assembly of unmapped reads and customized analysis in highly variable loci. Among the >400 million variants detected, 97% have frequency <1% and 46% are singletons. These rare variants provide insights into mutational processes and recent human evolutionary history. The nearly complete catalog of genetic variation in TOPMed studies provides unique opportunities for exploring the contributions of rare and non-coding sequence variants to phenotypic variation. Furthermore, combining TOPMed haplotypes with modern imputation methods improves the power and extends the reach of nearly all genome-wide association studies to include variants down to ~0.01% in frequency.

---

Advancing DNA sequencing technologies and decreasing costs are enabling researchers to generate collections of human genetic variation at an unprecedented scale^1,2^. For these advances to improve understanding of human health and disease, they must be deployed in well phenotyped human samples and used to build resources such as variation catalogs^2–4^, control collections^5,6^ and imputation reference panels^7–9^ that will enhance all ongoing human genetic studies. Here, we describe high-coverage whole genome sequencing (WGS) of 53,831 TOPMed samples for which data are now available to qualified researchers through dbGaP.

Long-term goals of the TOPMed program include (1) characterizing the genetic architecture of phenotypic variation in heart, lung, blood, and sleep (HLBS) disorders and related phenotypes; (2) identifying causal genetic variants and assessing how they may interact with environmental factors; (3) characterizing the spectrum of disease types; (4) understanding ethnic differences in these disorders; and (5) establishing a foundation for personalized interventions for disease prediction, prevention, diagnosis, and treatment. We are pursuing these goals through generation of whole genome sequence (WGS) and “omics” data for research participants of diverse backgrounds and with deep phenotypic characterization through ongoing studies, and by developing analytical tools to effectively mine the resulting data.

The program currently consists of >80 participating studies^10^, ~1,000 investigators, and >30 Working Groups^11^. Study designs include prospective cohorts, families, population isolates, and case-only collections. The studies have measured thousands of phenotypic and related environmental risk factors. Some studies focus on heart (38%), lung (33%), blood (8%), or sleep phenotypes (1%), while others cover many phenotypic areas (20%) (Supplementary Figure 1). High-coverage whole genome sequencing (WGS, mean 38X coverage) is in progress for ~145,000 participants (with ~135,000 completed), with additional “omics” assays in progress for tens of thousands of participants with sequenced genomes. Approximately 82% of participants are U.S. residents with diverse ancestry and ethnicity (40% European, 32% African, 16% Hispanic/Latino, 10% Asian, and 2% other; Supplementary Figure 2). Supplementary Information (section 1.1) provides additional information about TOPMed design features, study selection, and participant characteristics.

Here we describe WGS from the first 53,831 TOPMed samples selected from data sets that are now available via dbGaP controlled-access (Supplementary Tables 1 and 2); additional data will be made available as quality control (QC), variant calling and dbGaP curation are completed. Our work identifies and characterizes the rare variants that comprise the majority of human genomic variation^7,12–14^ and extends previous efforts that relied on genotyping arrays^15–17^, low-coverage WGS^7,8^, exome sequencing^2,12,18^, or analyses of smaller sample collections^19–22^. Since rare variants represent more recent and potentially more deleterious changes that can impact protein function, gene expression, or other biologically important elements, their discovery and study are crucial for understanding the genetics and biology of human health and disease^13,23,24^. The TOPMed WGS dataset is the largest collection of human genetic variants that is now broadly available to researchers (through the <u>BRAVO browser</u>^25^) and has substantially more ancestral diversity than other whole genome datasets^2,26^ (Supplementary Figure 3).

## TOPMed WGS

TOPMed WGS data processing is performed periodically to produce genotype data “Freezes” that include all samples available at a given time. The Freeze 5 genotype call set was based on 65,000 samples, of which ~55,000 have now been curated and released on dbGaP. The 53,831 samples described here are drawn from this set.

To evaluate the reproducibility of our genotype calls, non-reference allele discordance was estimated for 378 samples sequenced in duplicate. For each pair of duplicates, discordances were counted as any difference in called genotypes, regardless of read depth. For single nucleotide variants (SNVs), the median discordance rate across duplicate pairs was 0.040% for passing and 33% for variants failing site-level quality filtering. For indels, the median discordance rate was 0.61% for passing and 23% for failing variants. Considering only singleton variants in a set of unrelated samples, the median discordance was 0.23% for passing, 34% for failing SNVs, 0% for passing indels, and 47% for failing indels. These reproducibility estimates indicate the high quality of the genotype calls, although they may be optimistic because members of each duplicate sample pair processed and called jointly.

We also evaluated potential benefits from high coverage WGS relative to exome sequencing (depth >30X)^18^ and low coverage WGS (depth >6X)^22^ in 430 Framingham Heart Study samples. TOPMed WGS identified 23.8 million variants in these samples, compared with 20.5 million variants in low coverage sequencing analysis (a 16% increase). Essentially all the newly observed variants had minor allele frequency <0.5%. When we restricted the analysis to coding regions, TOPMed identified ~17% more coding variants than both low-coverage WGS and exome sequencing (Supplementary Table 3). In comparison to exome chip genotypes for these samples, TOPMed averaged 0.999 concordance (across all frequencies), compared with a concordance of 0.990 to 0.998 for low coverage WGS. These comparisons reflect TOPMed’s increased sequencing depth, as well as its joint calling over tens of thousands of samples.

### 410 million genetic variants in 53,831 samples

A total of 7.0×10^15^ bases of DNA sequence data were generated, consisting of an average of 129.6×10^9^ bases of sequence distributed across 864.2 million paired reads (each 100-151-bp long, average 149.7) per individual. For a typical individual, 99.65% of the bases in the reference genome were covered, to a mean read depth of 38.2X.

Sequence analysis identified 410,323,831 genetic variants (381,343,078 SNVs and 28,980,753 indels), corresponding to an average of one variant per 7 bp throughout the reference genome (Table 1). Overall, 78.7% of these variants had not been described in dbSNP build 149 before the TOPMed Project (TOPMed variants account for the majority of variants in more recent dbSNP releases). Among all variant alleles, 46.0% were observed once across all samples (i.e. singletons). Since natural selection removes most deleterious variants from the population before they become common, the fraction of singletons closely tracked functional constraints. For example, among all 4,973,175 protein coding variants, the proportion of singletons was highest for the 113,805 frameshift variants (59.7%), high among the 105,042 putative splice and truncation variants (54.1%), intermediate among the 3,172,551 non-synonymous variants (48.1%), and lowest among the 1,525,971 synonymous variants (43.0%). Beyond protein coding sequences, we found elevated proportions of singletons in promoters (47.7%), 5’ untranslated regions (47.2%), regions of open chromatin (46.2%), and 3’ untranslated regions (46.1%) (Supplementary Table 4). Conversely, we found lower proportions of singletons in putative transcriptional repressor *CTCF* (45.4%) and transcription factor binding sites (45.7%), suggesting these regions are relatively tolerant of variation.

**Table 1.** Coverage, sequencing depth and number of variants.

|  | Total | Singletons (%) | Per Individual |  |  |  |
| --- | --- | --- | --- | --- | --- | --- |
|  |  |  | Average | 5<sup>th</sup><br>Percentile | Median | 95<sup>th</sup><br>Percentile |
| **Bases (Gb)** | 6,973,670 | - | 130 | 107 | 128 | 157 |
| **Depth (x)** | - | - | 38 | 31 | 38 | 46 |
| **Genome Covered (%)** | - | - | 98.5 | 96.2 | 99.2 | 99.9 |
| **Depth >10x** | - | - | 97.9 | 95.4 | 98.7 | 99.6 |
| **Total Variants** | 410,323,831 | 188,947,391 (46) | 3,776,362 | 3,515,416 | 3,567,439 | 4,364,075 |
| **SNVs** | 381,343,078 | 175,419,690 (46) | 3,579,423 | 3,334,782 | 3,383,710 | 4,129,868 |
| **Indels** | 28,980,753 | 13,527,701 (47) | 196,940 | 180,567 | 183,759 | 234,245 |
| **Novel\* Variants** | 323,113,479 | 178,243,307 (55) | 30,207 | 20,363 | 26,347 | 44,379 |
| **SNVs** | 298,028,808 | 165,082,153 (55) | 25,861 | 17,568 | 22,909 | 36,897 |
| **Indels** | 25,084,671 | 13,161,154 (52) | 4,345 | 2,752 | 3,378 | 7,392 |
| **Coding Variation** |  |  |  |  |  |  |
| **Synonymous** | 1,525,971 | 656,746 (43) | 11,743 | 10,840 | 11,073 | 13,693 |
| **Non-synonymous** | 3,172,551 | 1,527,247 (48) | 11,468 | 10,633 | 10,875 | 13,237 |
| **Stop/Essential Splice** | 105,042 | 56,801 (54) | 478 | 426 | 456 | 568 |
| **Indels** |  |  |  |  |  |  |
| **Frameshift** | 113,805 | 67,903 (60) | 133 | 112 | 127 | 167 |
| **Inframe** | 55,806 | 27,118 (49) | 103 | 85 | 99 | 129 |

\* Variant was not present in dbSNP b149, the most recent dbSNP version without TOPMed submissions

We identified an average of 3.78 million variants in each studied genome. Among these, an average of 30,207 were novel (0.8%) and 3,510 were singletons (0.1%). Thus while there are vast numbers of rare variants in humans, only a few of these are present in each genome and large numbers of genomes must be studied to draw inferences about their function. Also striking is the observation that while, among all variants, we observed 3.17 million non-synonymous and 1.53 million synonymous variants (a 2.1:1 ratio), individual genomes contained similar numbers of non-synonymous and synonymous variants (11,743 non-synonymous, 11,768 synonymous, on average, Table 1). The difference can be explained if more than half of non-synonymous variants are removed from the population before they become common and, indeed we observed a relative paucity of non-synonymous variants at higher frequencies (56,428 non-synonymous to 52,889 synonymous at >0.5% frequency; 21,150 non-synonymous to 22,018 synonymous at >5% frequency).

### Protein loss of function variants

A particularly interesting class of variants are the 228,966 putative loss of function (pLoF) variants, which include premature stop codons, frameshifts, and mutations in splice acceptor and donor sites (Table 2). This class includes the highest proportion of singletons among all variant classes we examined. An average individual carried 2.5 unique pLoF variants; across all individuals, we observed pLoF variants in 18,493 (95.0%) of GENCODE v29^27^ genes. Significantly, we identified more pLoF variants per individual than in previous surveys based on exome sequencing -- an increase that was mainly driven by the identification of additional frameshift variants (Supplementary Table 5) and by more uniform and complete coverage of protein coding regions (for example, whereas ~88.9% of protein coding bases are covered at depth >10X in the Exome Aggregation Consortium^2^ (ExAC) data, ~98.8% of such bases are covered at depth >10X in TOPMed; Supplementary Figures 4 and 5). Our catalog of variants is available to researchers at the BRAVO browser^28^ and through dbSNP. Both are important resources for studies of Mendelian disorders, which must find proverbial “needles-in-a-haystack” by distinguishing novel, likely deleterious variants from other variants segregating in the population.

**Table 2.** Putative loss of function (pLoF) variants.

|  | Total | Singletons (%) | Per Individual |  |  |  |
| --- | --- | --- | --- | --- | --- | --- |
|  |  |  | Average | 5<sup>th</sup><br>Percentile | Median | 95<sup>th</sup><br>Percentile |
| **pLoF** | 228,966 | 58.5 | 209 | 182 | 202 | 251 |
| **Stop gained** | 79,766 | 55.6 | 72 | 60 | 72 | 87 |
| **Frameshift** | 100,393 | 60.3 | 92 | 77 | 90 | 115 |
| **Splice** | 48,807 | 59.6 | 44 | 34 | 43 | 57 |
| **pLoF (AF<0.5%)** | 217,795 | 58.8 | 20.7 | 10 | 19 | 35 |
| **Stop gained (AF<0.5%)** | 75,904 | 55.8 | 7.9 | 3 | 7 | 15 |
| **Frameshift (AF<0.5%)** | 95,064 | 60.6 | 8.3 | 3 | 8 | 15 |
| **Splice (AF<0.5%)** | 46,827 | 59.9 | 4.5 | 1 | 4 | 9 |

We searched for gene sets with fewer rare (AF < 0.5%) pLoF variants than expected based on gene size. The gene sets with strong functional constraint included genes encoding DNA and RNA binding proteins, spliceosomal complexes, translation initiation machinery, and RNA splicing and processing proteins (Supplementary Table 6). Importantly, genes associated with human disease in COSMIC^29^ (31% depletion, p<1×10^−6^), GWAS Catalog^30^ (8% and 9% depletion in downstream and upstream genes correspondingly, p<1×10^−6^), OMIM^31^ (4% fewer than expected, p=5×10^−5^) and ClinVar^32^ (4% depletion, p=8×10^−4^) also all contained fewer rare pLoF variants than expected.

### The distribution of genetic variation

We examined the distribution of variant sites across the genome by counting variants in 1Mb of contiguous sequence ond in 1Mb segments containing ordered concatenations of sequence with similar conservation level (indicated by CADD score) or coding versus non-coding status (Figure 1). Using 40,772 unrelated samples, we find that the vast majority of human genomic variation is rare^12,13^ and located in putatively neutral, non-coding regions of the genome (Figure 1A). While coding regions have lower average levels of both common and rare variation, we find ultra-conserved non-coding regions with even lower levels of genetic variation, consistent with previous findings^33^ (Figure 1A).

**Figure 1.**
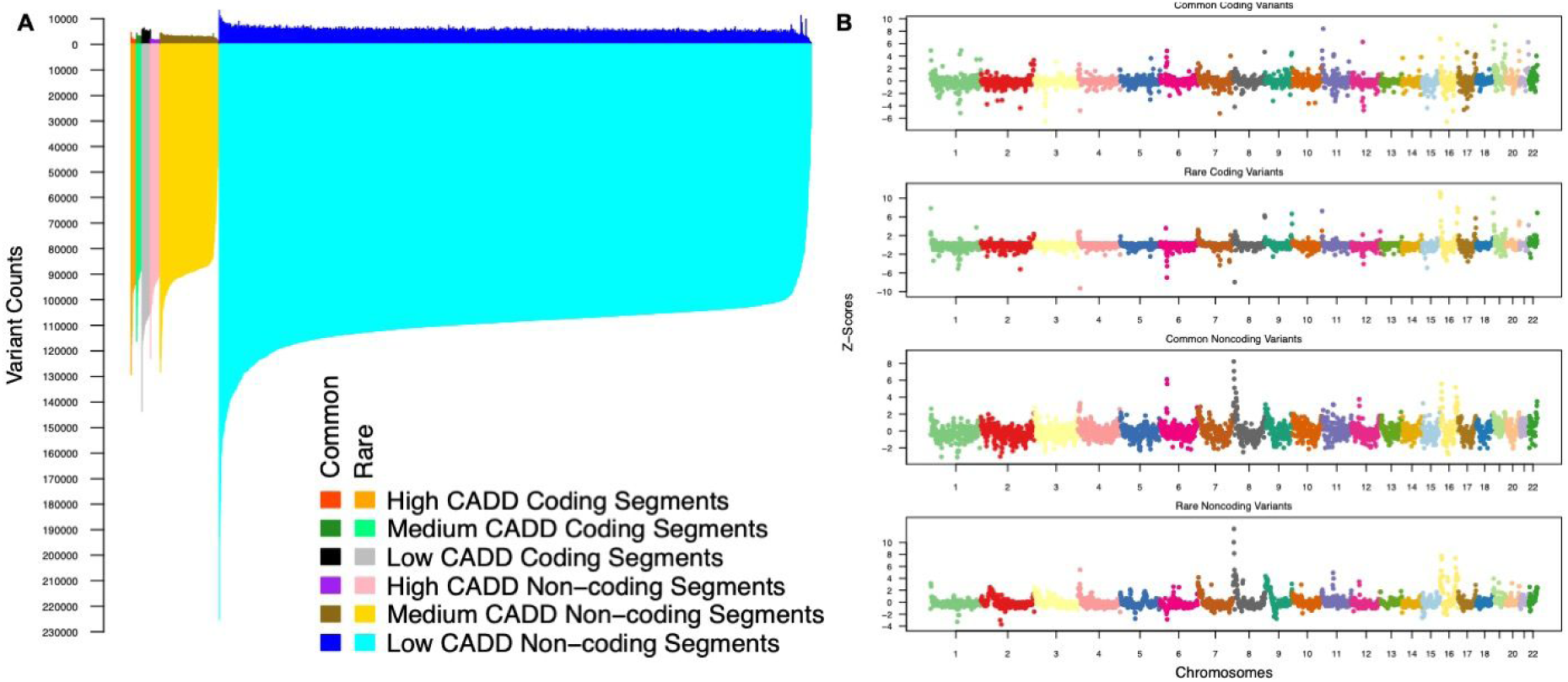
Distribution of genetic variants across the genome. **A.** Common (allele frequency ≥ 0.5%) and rare (allele frequency < 0.5%) variant counts are shown above and below the x-axis, respectively, and each 1MB concatenated segment is sorted based on their number of rare variants. Non-coding regions of the genome with CADD scores below 10 (lower predicted function) have the largest levels of common and rare variation (dark and light blue, respectively), followed by low CADD coding regions (black and grey). Overall, the vast majority of human genomic variation is comprised of rare variation. **B.** Levels of variation are shown across the genome for common coding variants (panel 1), rare coding variants (panel 2), common noncoding variants (panel 3), and rare noncoding variants (panel 4). Variation levels are represented on the Z-score (X-mean/SD) of the adjusted variant counts per 1MB contiguous segment for each variant category.

After filtering to focus on regions of the genome that are accessible through short read sequencing, most contiguous 1 Mb segments show similar levels of common (5,141 土 1,298 variants with MAF≥0.5%) and rare variation (120,414土19,862 variants with MAF<0.5%) (Figure 1B). However, outlier segments with notably high or low levels of variation do exist, and likely represent biologically interesting genomic regions that can serve as candidates of interest in follow-up analyses. One region of interest on chromosome 8p (GRC 38 positions 1,000,001-7,000,000 bp) has the highest overall levels of variation (Figure 1B). This region is estimated to have the highest mutation rate across the human genome (with the possible exception of the Y chromosome)^34^. Interestingly, high levels of variation in this region are driven primarily by non-coding variation.

While levels of common and rare variation within segments are significantly correlated (R^2^ = 0.462, p-value ≤ 2×10^−16^, Supplementary Figure 6), there are outliers. For example, segments overlapping the Major Histocompatibility Complex (MHC) genes have the highest levels of common variation across the genome while also having some of the lowest levels of rare variation, consistent with the evolutionary consequences of balancing selection^35–37^. Segments with a high proportion of coding bases are negatively correlated with overall variation, and feature a strong depletion in the number of common variants and a more modest depletion in rare variants. This is consistent with the expectation that rare variants observed in very large samples (such as ours) have not yet been effectively pruned by natural selection (Supplementary Figure 7).

### Insights into mutation processes

The distribution of singleton variants in large studies such as ours has been only modestly shaped by purifying selection and largely reflects underlying mutation processes^38^. Thus, studying singletons and other rare variants in our sample offers opportunities to dissect the mutation processes that generate extant human variation. We explored the spatial clustering of genetic variants, focusing on 32,058,136 singleton SNVs ascertained in a subset of 2,000 unrelated individuals (equal numbers of African and European ancestry). We particularly tried to dissect multi-nucleotide mutation events where two or more closely-spaced mutations arise simultaneously as a result of error-prone replication and repair mechanisms^39^.

If neighboring singletons were the result of independent mutation events, fewer than 0.07% of the singletons in an individual in this sample should be <100bp apart, while in fact, 2% of our singletons meet this criterion. We suspect closely spaced singletons reflect the consequences of translesion synthesis, an error-prone replication process in which specialized polymerases bypass DNA lesions at stalled replication forks^40^, often causing multiple mutations within very short spans^39^.

To explore the relative importance of different mutation processes and how they influence clustering of genetic variants, we modeled the spatial distribution of singletons in each individual as a mixture of exponential processes (Supplementary Figure 8). Each component represents a distinct mutational process, resulting in unique spacing and mutation patterns. We found clear evidence for four classes of clustered singletons, consistent across individuals and ancestries (Figure 2A). Class 1 represents singletons occurring an average of ~2 - 8 bp apart accounted for ~1.5% of singletons in each sample These appear to arise through translesion synthesis and are substantially enriched for A>T and C>A transversions (Figure 2B), consistent with known signatures of translesion synthesis errors^41,42^. Class 2 singletons occurring ~500 - 5,000 bp apart accounted for ~12-24% of singletons. Some of these variants likely result from hypermutability of single-stranded DNA intermediates during repair of double-strand breaks^43,44^ and are enriched for C>G transversions. They show prominent subtelomeric concentrations on chromosomes 8p, 9p, 16p, and 16q^43,44^ (Figures 2B and 2C, Supplementary Figure 9), consistent with evidence that subtelomeric regions are enriched for double-strand breaks in eukaryotic genomes^45^. Class 3 singletons occurring ~30,000 - 50,000 bp apart accounted for ~43-49% of all singletons. Class 4 singletons occurring ~125,000 - 170,000 bp apart accounted for ~31-37% of all singletons. Singletons in classes 3 and 4 show the same mutational spectrum as the genomic background, suggesting that these represent independent mutation events whose spacing is mainly driven by variation in local ancestry. Since multi-nucleotide mutations are uniquely implicated in the genetic architecture of complex diseases^46^ and the evolutionary history of the genome^47^, our findings illustrate how the TOPMed sample will facilitate more refined and robust approaches to interpreting patterns of genetic diversity.

**Figure 2.**
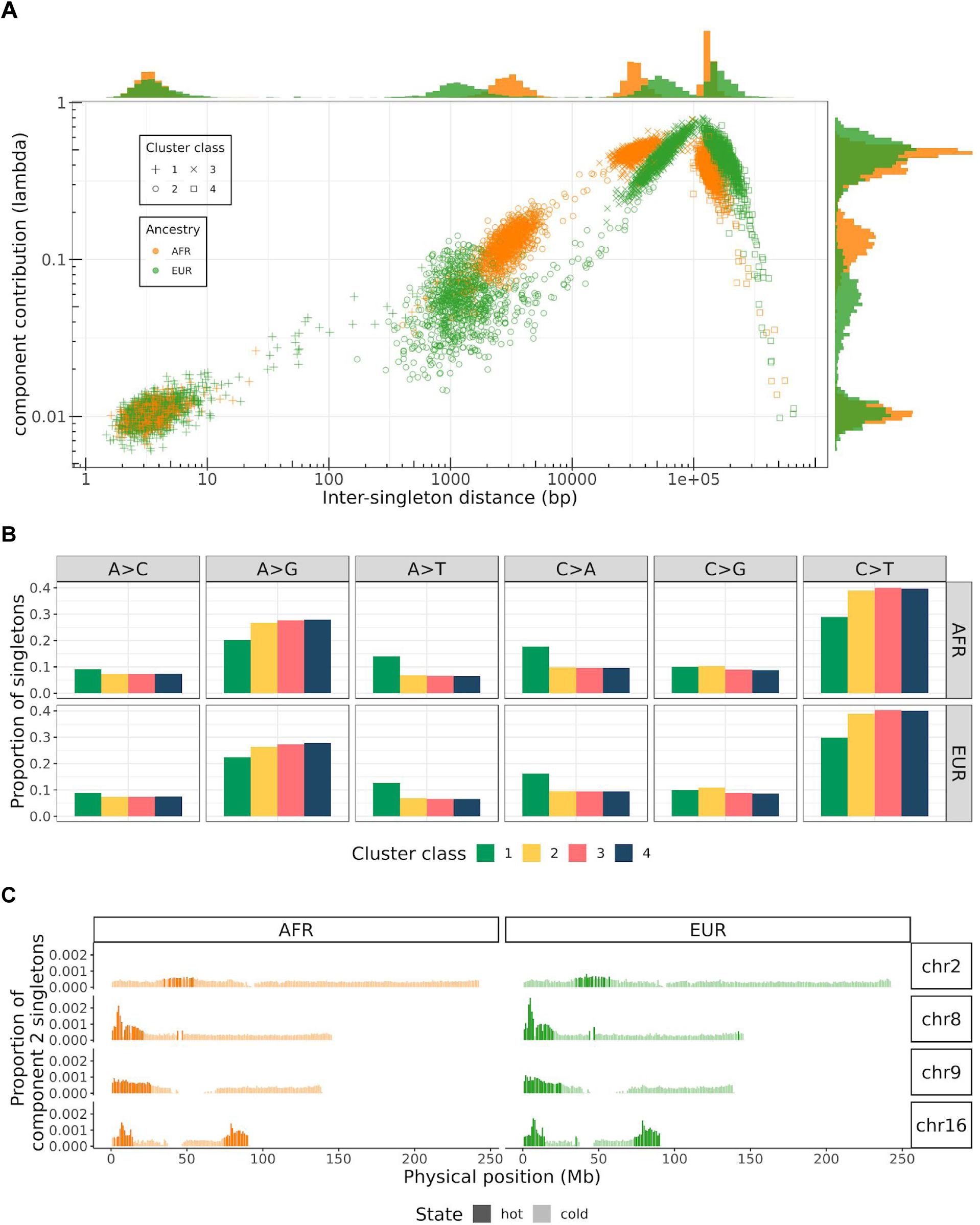
**A.** Parameter estimates for exponential mixture models of singleton density. Each point represents one of the four components in one of the 2,000 individuals in the sample, colored by the majority ancestry of that individual. The rate parameters of each component are shown across the x-axis, and the lambda parameters (i.e., the proportion that component contributes to the mixture) on the y-axis (on a log-log scale). Marginal histograms show the distribution of the lambda and rate parameters for each component. **B.** Mutational spectra of singletons assigned to each of the four cluster classes, separated by population. **C.** Density of cluster class 2 singletons in 1 Mbp windows on chromosomes 2, 8, 9, and 16. Windows with class 2 singleton counts above the 95th percentile (calculated genome-wide) are classified as hotspots and are represented with a darker shade.

### Beyond SNVs and Indels

To evaluate the potential of our data to generate even more comprehensive variation datasets, we developed and applied a method based on *de novo* assembly of unmapped and mismapped read pairs, enabling us to assemble sequences present in a sample but absent or improperly represented in the reference.

We identified ~200 - 500kb of non-reference hominid sequence per sequenced individual (N50 length of 954 bp, range: 200bp to 9,330bp). Merging across ~17,000 samples included in this analysis, we collapsed this to 1,627 scaffolds containing 1,932 contigs and spanning 2,179,874bp. From this set, we fully resolved the insertion sequence and precise breakpoint locations on the human genome for 737 contigs (totalling 1,160,253bp, of which 548,749 were inserted bases; largest contig: 12,488bp) and were able to locate one breakpoint for an additional 397 contigs (453,888bp; 323,675 putatively inserted/hanging bases) (Supplementary File 1).

In line with prior observations suggesting that the vast majority of human non-reference sequence is present in the assembled genomes of non-human primates^48,49^, we find that our assemblies likely represent retained ancestral sequences that have been deleted in some human lineages, including on the reference haplotype. Consistent with this, the frequencies of the newly assembled alleles (Supplementary Figure 10) are higher than those observed for SNVs and indels, with 78.3% of the events present in >5% of the samples and only 6% having a frequency <0.5%. Comparing our findings to two previous studies on different smaller datasets^48,49^, 243 sequences (164,099bp retained sequence) are wholly novel. Additionally, we have resolved length and both breakpoints for 137 events (170,133bp) for which only one breakpoint was previously known (Figure 3D).

Overall, retained ancestral sequences appear evenly distributed across the genome (Figure 3A), occurring at near expected frequencies with respect to genes. Of the 737 fully resolved events, 331 are intergenic and 406 overlap with GENCODE v29^27^ genes (15 exonic, 370 intronic, 21 in promoters, Figure 3B). Interestingly, we identified a sequence, present in all individuals, that fully contains exons of the *UBE2QL1* gene (including the translation start site). The sequence is absent from current human annotation but present in *Mus musculus* and *Rattus norvegicus*, suggesting this common gene is not currently represented in the reference genome (Supplementary Figure 11).

**Figure 3.**
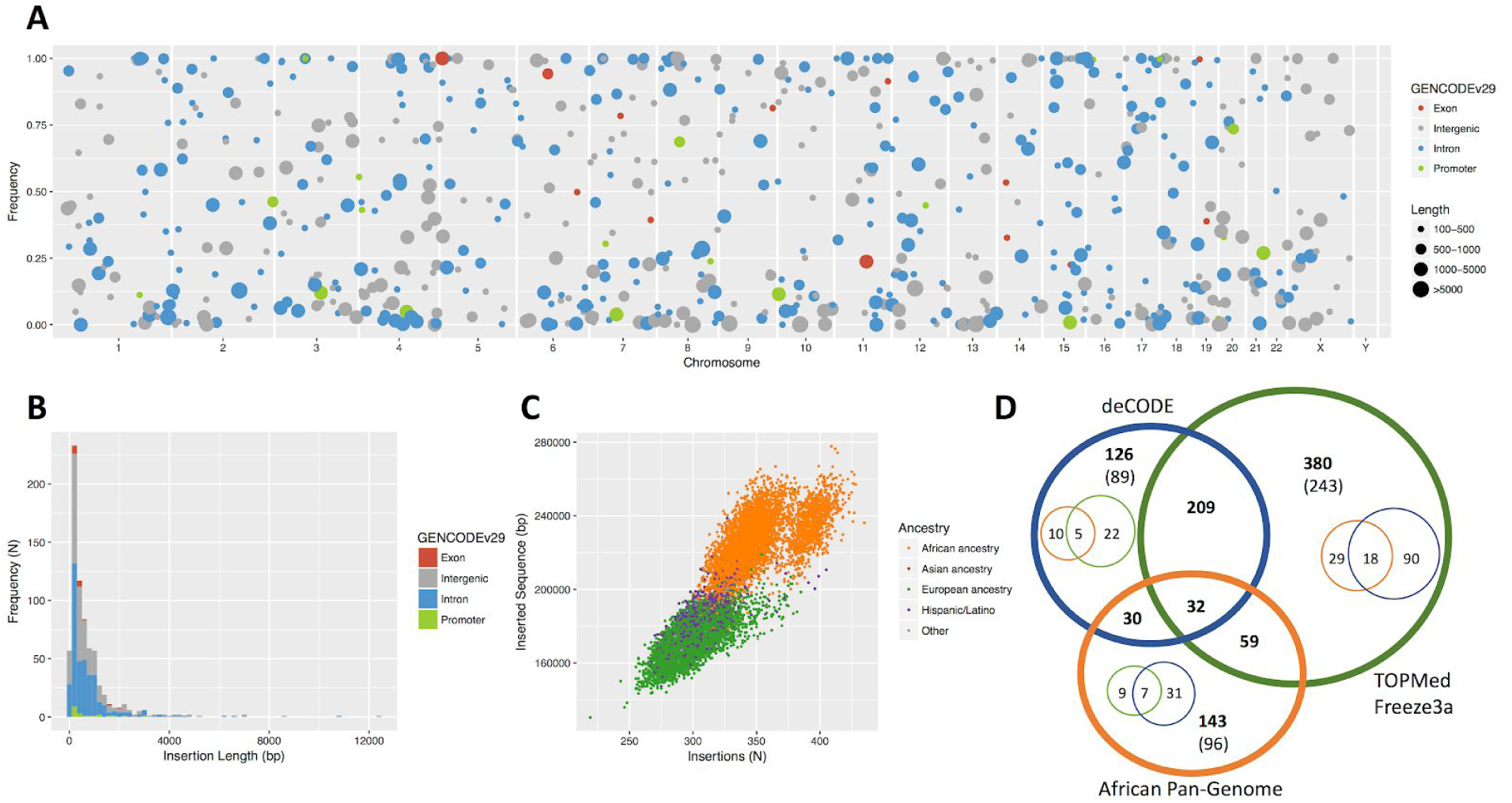
Retained ancestral sequences (>=100bp) discovered from unmapped reads. A) Cohort population frequency plotted over genomic location. Size of dot indicates insertion size, color indicates GENCODE annotation. B) Insertion length distribution. C) Number of insertion events vs. number of inserted bases, per individual by ancestry. D) Venn diagram showing positional concordance with insertions (small circles: one-ended breakends) identified in previous studies on different datasets^48,49^.

On average, African ancestry individuals have both more non-reference sequences (350.4 sequences/sample vs 297.7 sequences/sample) and greater total assembly size (226.2kb/sample vs 171.6kb/sample, respectively) than European-ancestry individuals (Figure 3C). This is consistent with a loss of genetic variation in the out-of-Africa bottleneck. Overall, 5 sequences spanning 15,653bp are found only in African samples, whereas no single fully resolved sequence is unique to Europeans; we do identify one partially resolved 605bp break end in a single European. All these ancestry-specific events are rare (present in <0.5% of samples), whereas the overwhelming majority of events are shared by multiple continental ancestry groups.

### Variation in CYP2D6

A complementary approach to *de novo* genome assembly is to develop approaches that combine multiple types of information — including previously observed haplotype variation, SNVs, indels, copy number, and homology information — to identify and classify haplotypes in interesting regions of the genome. One such region is the *CYP2D6* gene locus, which encodes an enzyme that metabolizes approximately 25% of drugs and whose activity varies substantially among individuals^50–52^. More than 100 *CYP2D6* haplotypes have been described to date, some involving a gene conversion with its nearby nonfunctional but highly similar paralog *CYP2D7*. These haplotypes are typically labelled using star alleles (e.g., *CYP2D6*1*, *\*2*), each defined by SNVs, indels, structural variants (SVs), or some combination of these.

We were particularly interested in understanding whether current variation catalogs for *CYP2D6* were complete, especially in non-European individuals. Therefore, we performed *CYP2D6* haplotype analysis for 3,418 African American individuals from the Jackson Heart Study (JHS) using the Stargazer program^50^. We called a total of 56 star alleles (51 known and 5 novel) representing increased function, decreased function, and loss of function (Supplementary Table 7). The five novel star alleles were comprised of gene duplications (*CYP2D6*29×3*, *\*34×2*, and *\*42×2*) and multiplications (*CYP2D6*1×3* and *\*2×3*). We observed various *CYP2D6/CYP2D7* hybrids and extensive copy number variation ranging from zero to five gene copies (Supplementary Figure 12). Based on the *CYP2D6* genotypes from 1,923 unrelated JHS individuals, we estimated that 26% of these individuals carry SVs, and that 2%, 9%, 83%, 4%, and 2% of the individuals are poor, intermediate, normal, ultrarapid, and unknown metabolizers^53^, respectively.

### Heterozygosity and rare variant sharing among ancestrally diverse sampled individuals

The TOPMed variation data also present an opportunity to examine expectations about rare variation studies — for example, which types of studies share the same rare variants and might be expected to provide the most concordant results? Which studies show unique or distinct patterns of variation and might be expected to provide unique insights? We grouped TOPMed participants by study and by self-reported ancestry and calculated genetically determined ancestry components, heterozygosity, number of singletons, and rare variant sharing (Figure 4, Supplementary Table 8).

**Figure 4.**
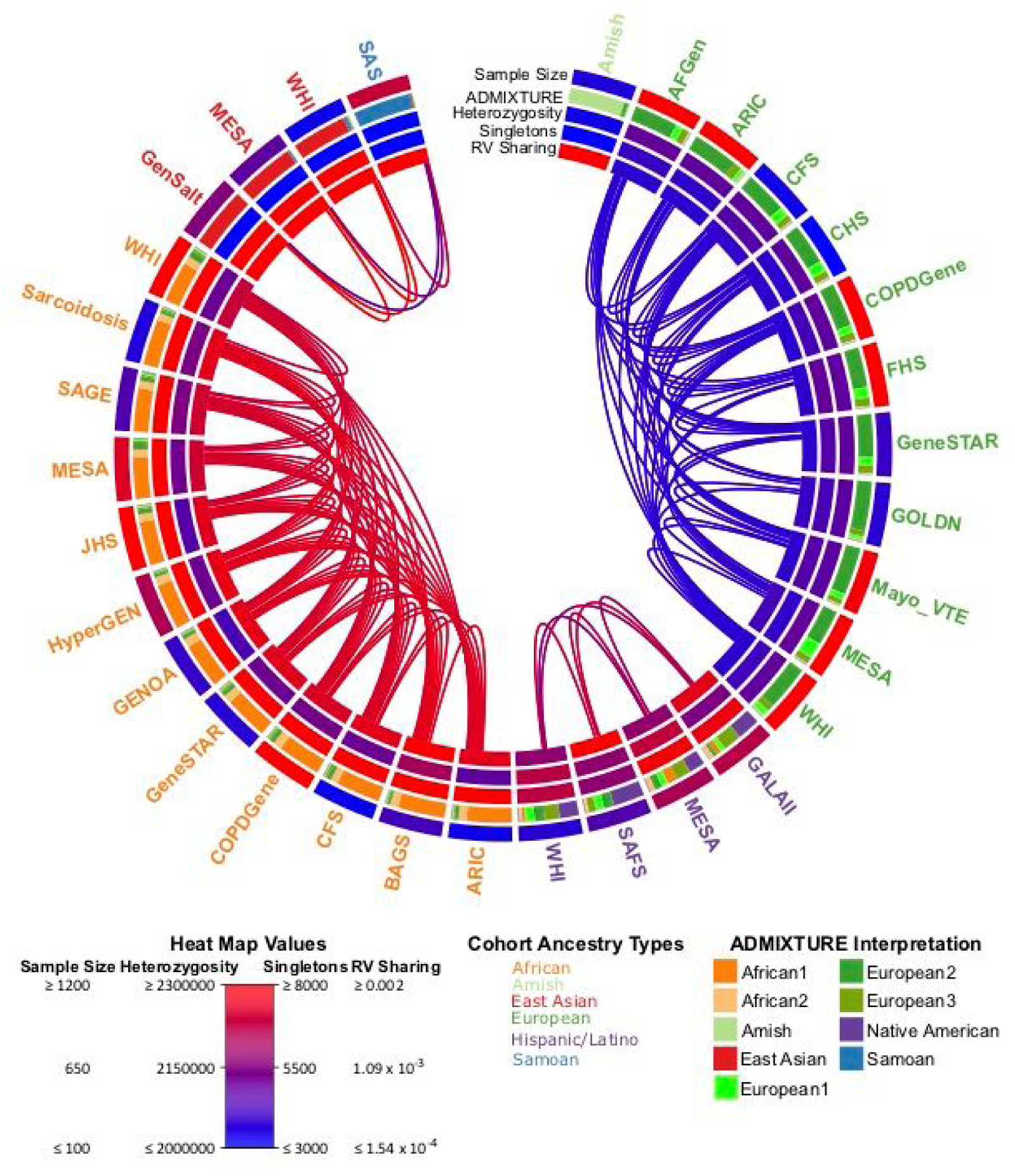
Ancestry, genetic diversity and rare variant genetic relatedness across the TOPMed studies. Each study label is shaded based on their self-described ancestry. From the outside moving inwards each track represents: the unrelated sample size of each study used in these calculations, average ADMIXTURE values, average number of heterozygous sites in each individual’s genome, average number of singleton variants in each individual’s genome, and the average within study rare variant sharing (RV Sharing). The links depict the 75th percentile of between study rare variant sharing. All between study rare variant sharing comparisons can be found in Supplementary Figure 13. The sample size, heterozygosity, and singleton average values can be found in Supplementary Table 8.

We find that African ancestry studies have the greatest heterozygosity^7,54^, followed by Hispanic/Latino, European, Amish, East Asian, and then Samoan studies -- consistent with a gradual loss of heterozygosity tracking recent African origin of modern humans and subsequent migration from Africa to the rest of the globe. Interestingly, while the East Asian ancestry studies have among the lowest heterozygosity in our sample (even lower than the Amish, a European ancestry founder population with notably low heterozygosity^55,56^), the East Asian ancestry studies have the greatest singleton counts (in contrast to the Amish, who have the lowest). These observations are consistent with a prolonged bottleneck in East Asian populations followed by extreme recent exponential growth, but could also reflect a more modest sampling of East Asian variation in TOPMed. The Amish on the other hand have a more recent bottleneck and have not experienced as great a growth since founding^55,56^ (see Supplementary Information section 1.5).

Using rare variation, we are also able to distinguish fine-scale patterns of population structure (Figure 4, Supplementary Figure 13; Supplementary Information section 1.6). Broadly, we observe sharing between studies with shared continental ancestry (whether African, European, Asian, or Hispanic/Latino). Nevertheless, additional patterns emerge. The Amish are unique among the included studies: they exhibit little rare variant sharing with outside groups and also the greatest rare variant sharing within study - consistent with a marked founder effect. Further, we observe ~4x greater rare variant sharing between African ancestry studies than between European ancestry studies, even after correcting for sample size differences (Supplementary Figure 14). We also observe that the MESA and GALAII Hispanic/Latino studies share ~2x more rare variants with the African ancestry studies than the SAFS and WHI Hispanic/Latino studies, consistent with ADMIXTURE analysis results and the expectation that these studies contain individuals from populations with greater African admixture (Figure 4, Supplementary Figure 13).

### Haplotype sharing

A corollary to rare variant sharing is rare haplotype sharing through segments inherited from a recent common ancestor. These identical-by-descent (IBD) segments show similar patterns of within and between study sharing to the rare variants (Supplementary Figures 15 and 16). The Amish study shows the greatest average within study IBD sharing levels (Supplementary Figure 15), consistent with a founder event 14 generations ago and a subsequently closed society^56,57^. In the Amish and other samples, the distribution of IBD segments allows estimates of effective population sizes over the last 300 generations (Figure 5). The demographic histories are broadly similar between ancestry groups, with the exception of the Amish, who experienced a more extreme bottleneck when moving to the New World, and the Samoans, who have had a smaller effective population size than the East Asian populations from which they separated ~5000 years ago^58,59^. Both non-Amish European ancestry and African ancestry populations appear to have experienced a bottleneck ~5-10 generations ago, consistent with moving to the New World, whether through colonization or forced migration.

**Figure 5.**
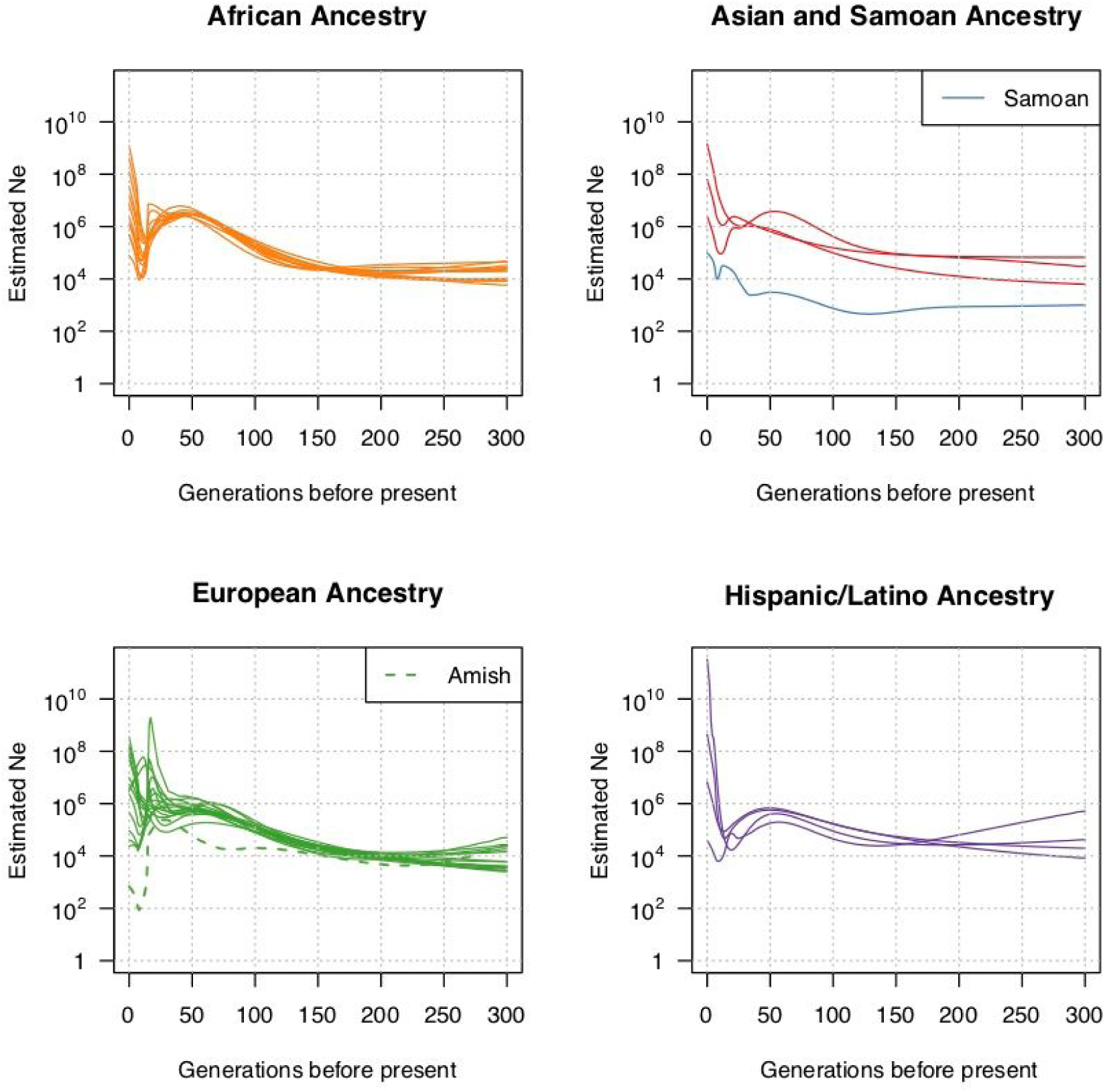
Estimates of recent effective population size by ancestry. Each line represents the estimate from a single study, considering only individuals self-reporting the given ancestry. The included studies are the same as those in Supplementary Figure 15. The Amish and Samoan results are individually identified in the plots due to their distinct recent population size trajectories. N_e_ indicates effective population size.

### Large sample sizes alleviate impact of selection at linked sites

The relative numbers of singletons, doubletons and other very rare variants contained at neutral sites can be used to infer human demographic history^13,60,61^. While most demographic inference is currently based on fourfold degenerate sites in protein sequences, these sites evolve under the influence of strong selection at nearby linked protein coding sites^62,63^, which can affect the inferred timing and magnitude of population size changes. WGS enabled us to access intergenic regions of the genome that are minimally affected by selection and complement the analyses of fourfold degenerate sites in protein coding regions. In addition, the recent expansion of the human population has led to an increase in the number of rare mutations in the human genome^12,64^ that may not have had time to be influenced by selection at linked sites^64^. While these recent mutations will dominate rare variant portions of the site frequency spectrum (SFS) and may lessen differences caused by selection at linked sites, they can only be identified with extremely large sample sizes. We measured how the site frequency spectrum and demographic inference changes as a function of sample size and an index of selection at linked sites (McVicker’s *B* statistic^65^). Increasing the sample size demonstrated a more limited impact of selection at linked sites on singleton and doubleton bins of the SFS (Figure 6A; Supplementary Figure 17). Estimates of European effective population size based on the 1% of the genome with the weakest effect of selection at linked sites consistently yielded ~1.1M million individuals. In contrast, inference at fourfold degenerate synonymous sites was sensitive to sample size until >3,000 chromosomes were included in the analysis, at which point estimates using the two datasets converged (Figure 6B; Supplementary Figure 18; Supplementary Table 9). These results suggest demographic inference of recent human history will be facilitated by large sample size and the ability to focus on regions of the genome least affected by selection at linked sites.

**Figure 6.**
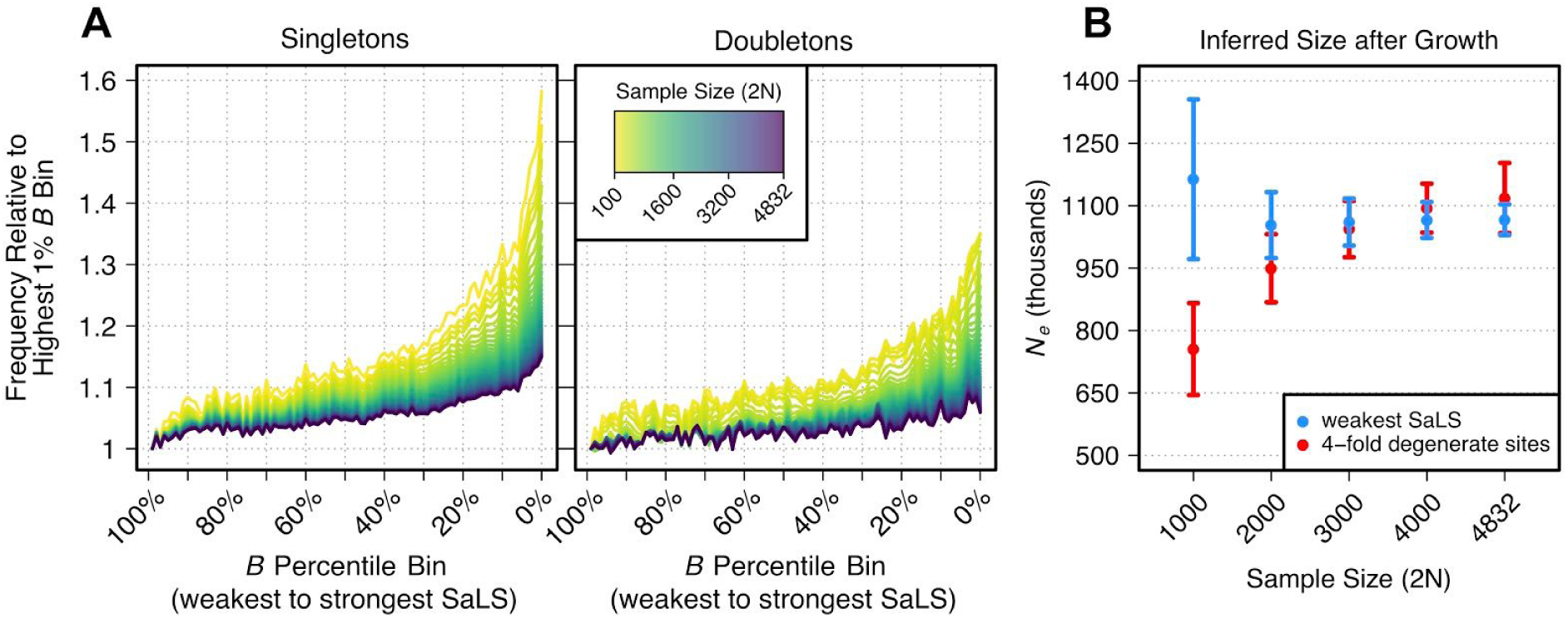
Relative increase in singletons and doubletons of the SFS across McVicker’s *B* and the population size inferred from demographic inference using various sample sizes. A) The relative increase in the singleton and doubleton bins of the SFS for decreasing percentile bins of McVicker’s *B* compared with the highest percentile bin of *B* (higher percentiles of *B* indicate weaker effects of selection at linked sites [SaLS]). These relative increases are plotted for different sample sizes (see legend). B) The population size inferred in the last generation of an exponential growth model for Europeans. Demographic inference was conducted with different sample sizes and by using different parts of the genome (see legend). Whiskers show 95% confidence intervals (see Supplementary Table 9 for parameter values).

### Human adaptations

When adaptive mutations occur their frequencies quickly rise, bringing surrounding linked variation to higher frequencies as well. If selection is strong enough, we expect to see regions of low diversity haplotypes flanking these events. We used the Integrated Haplotype Score^66^ (iHS) to search for regions where positive selection occurred or is ongoing. Overall, we find three regions with strong evidence of selection in all ancestry groups: one on chromosome 3 and two on chromosome 4. These regions center on three protein coding genes: *PIGG^67^*, *SLC30A9^68^*, and *STXBP5L^69^* (Supplementary Figure 19, Supplementary Table 10).

### TOPMed imputation resource

In addition to enabling detailed analysis of the combined genomic and health data for sequenced samples, TOPMed can enhance analyses of any genotyped samples. We constructed a TOPMed-based imputation reference panel of 60,039 individuals, including 239,756,147 SNVs and indels (see Supplementary Table 11 for breakdown by frequency). It is the first imputation reference panel based exclusively on deep WGS in diverse samples and greatly exceeds prior alternatives such as the Haplotype Reference Consortium (HRC)^8^ (39,635,008 autosomal SNVs) and the 1000 Genomes Project^7^ phase 3 (49,143,605 SNVs and indels) in resolution and accuracy (Supplementary Figure 21). For example, the average imputation quality for variants with frequency 0.001 increased from ~0.15 (1000 Genomes, HRC) to 0.91 (TOPMed) in African ancestry genomes and from 0.26 (1000 Genomes) and 0.59 (HRC) to 0.92 (TOPMed) for European ancestry genomes. Similar improvements were observable in other ancestries we considered except South Asians, which would be expected to improve in a future version of the panel that has more representation from this group. The minimum allele frequency at which variants could be well-imputed (r^2^>0.3) decreased from ~0.13-0.21% (1000 Genomes, European or African ancestry) to 0.05% (HRC, Europeans only) to ~0.004 - 0.006% (TOPMed, European or African ancestry) (Supplementary Figure 20). This means that 93% of the ~84,000 rare variants with minor allele frequency < 0.5% in an average African ancestry genome can be recovered through genotype imputation using the TOPMed panel (in comparison to 47% using the 1000G panel). Similarly, 95% of the ~69,000 variants with frequency <0.5% in an average European ancestry genome can be imputed using the TOPMed panel (in comparison to 49% using 1000G and 72% using HRC), enabling many human genetic studies to approximate the benefits of WGS for a modest investment in array genotyping and computation.

To illustrate the possibilities, we imputed TOPMed variants in 409,694 White British UK Biobank participants^1^, and performed association analyses for >1,400 binary phenotypes, defined as composites of ICD-10 billing codes^70^. We focused an initial analysis on 49,892 rare (frequency ≤0.5%) pLoF variants that were imputed (compared with 17,727 pLoF variants available from HRC imputation and 3,759 pLoF variants examined by Emdin et al.^71^). Among other findings, our analysis enabled several rare variant associations with breast cancer to be observed: we found a frameshift variant in *CHEK2* (chr22:28695868:AG:A; OR=2.04; P=6.98×10^−22^; minor allele count=2,111) and a stop gain variant in *PALB2* (chr16:23621362:C:T; OR=4.48; P=6.92×10^−14^; minor allele count=316) to be associated with breast cancer (12,636 cases and 200,417 controls). In addition to these two individual rare variants, we also found that a burden of rare pLoF variants in *BRCA2* (comprised of 23 rare pLoF variants; P=2×10^−8^; cumulative allele frequency cases vs. controls= 0.11% vs. 0.05%), the well-established risk gene for breast cancer, was significantly associated. Variants in these three genes are present in the ClinVar^32^ database as potentially pathogenic for familial breast cancer, but this is the first time these variants have been identified through a population of generally healthy adults not ascertained for cancer (Supplementary Table 12). In addition, identifying variant carriers in this sample of 500,000 well-characterized adults will help carefully dissect and understand the natural history and consequences of these variants. Other examples of rare variant association signals included associations with the burden of rare pLoF variants in *USH2A* and retinal dystrophies (83 cases; 34 rare pLoF variants; P=4×10^−8^; cumulative allele frequency cases vs. controls= 4% vs. 0.2%) and *IFT140* and acquired kidney cysts (1,257 cases; 15 rare pLoF variants; P=3×10^−8^; cumulative allele frequency cases vs. controls= 0.6% vs. 0.1%). These are all examples of signals that could previously only be studied through direct sequencing.

### Conclusion and future prospects

The first set of 53,831 sequenced genomes from TOPMed is now available to the community. The samples are deeply phenotyped and enable discovery of important biology^72–76^ for heart, lung, blood, and sleep disorders. In addition, they provide a rich resource for developing and testing methods for surveying human variation, for inference of human demography, and for exploring functional constraint in the genome. Beyond these uses, we expect TOPMed data will improve nearly all ongoing studies of common and rare disorders by providing both a deep catalog of variation in healthy individuals and an imputation resource that enables array-based studies to achieve completeness previously only attainable through direct sequencing.

The WGS and phenotypic resources for TOPMed studies are currently being enriched by applying transcriptomic, epigenomic, metabolomic, and proteomic assays to tens of thousands of samples selected through an open, peer-reviewed <u>process</u>^77^. Investigators affiliated with TOPMed studies share data and collaborate on cross-study analyses in Working Groups, from which many abstracts^78^ and publications^79^ are emerging. Members of the broader scientific community are using TOPMed resources through the WGS data available on dbGaP, the BRAVO variant server, and soon as an imputation reference panel on the Michigan Imputation Server.

Full utilization of the program’s resources by the scientific community will require new approaches to deal with the large size of the “omics” data (petabytes for read alignments, terabytes for genotype call sets); the diversity of phenotypic data types and structures; and the need for data sharing in an environment that supports privacy of participant data and respect for consent. These issues are currently being addressed in partnerships with the NIH Data Commons^80^ and NHLBI Data STAGE^81^ cloud-computing programs, which have both selected TOPMed as an initial data resource to be used for developing infrastructure and applications. Through these partnerships, investigators will be able to access the full range of TOPMed datasets, search and retrieve specific data types, and perform integrated analyses within a secure cloud environment that also supports data sharing among investigators within collaborative working groups.

## Methods

### DNA samples

Whole genome sequencing (WGS) for the 53,831 samples reported here was performed on samples previously collected and consented from research participants in 33 NHLBI funded research projects. All sequencing was done from DNA extracted from whole blood, with the exception of 17 Framingham samples (cell lines), and HapMap samples NA12878 and NA19238 (cell line) used periodically as sequencing controls.

### Whole genome sequencing

WGS targeting a mean depth of at least 30X (paired end, 150-bp reads) using Illumina HiSeq X Ten instruments occurred over several years at six sequencing centers (Supplementary Table 13). All sequencing used PCR-free library preparation kits purchased from KAPA Biosystems, equivalent to the protocol in the Illumina TruSeq PCR-Free Sample Preparation Guide (Illumina cat# FC-121-2001). Center-specific details are available from the TOPMed website^82^. In addition, 30X coverage WGS for 1,606 samples from four contributing studies were sequenced prior to the start of the TOPMed sequencing project and are included in this data set. These were sequenced at Illumina using HiSeq 2000 or 2500 instruments, have 2 x 100 bp or 2 x 125 bp paired end reads and sometimes used PCR amplification.

### Sequence data processing and variant calling

Sequence data processing was performed periodically to produce genotype data “Freezes” that include all samples available at a given time. All sequence was remapped using BWA-MEM^83^ to the hs38DH 1000 Genomes build 38 human genome reference including decoy sequences, following the <u>protocol</u>^84^ published by Regier et al. 2018^85^. Variant discovery and genotype calling was performed jointly, across TOPMed Parent studies, for all samples in a given Freeze using the <u>GotCloud</u>^86^ pipeline. This procedure results in a single, multi-study, genotype call set. A support vector machine (SVM) quality filter for variant sites was trained using a large set of site specific quality metrics and known variants from arrays and the 1000 Genomes Project as positive controls and variants with Mendelian inconsistencies in multiple families as negative controls (see <u>online documentation</u>^87^ for more details). After removing all sites with minor allele count less than 2, the genotypes with minimal depth >10X (minDP10) were phased using Eagle 2.4^88^. Sample-level quality control included checks for pedigree errors, discrepancies between self-reported and genetic sex, and concordance with prior genotyping array data. Any errors detected were addressed prior to dbGaP submission. Details regarding WGS data acquisition, processing, and quality control vary among the TOPMed data Freezes. Freeze-specific methods are described on the TOPMed <u>website</u>^89^ and in documents included in each TOPMed accession released on dbGaP (e.g., see document <u>phd007493.2</u>^90^ in phs000956.v3.p1).

### Sample sets

Several sample sets derived from three different WGS data Freezes were used in the analyses presented here: Freeze 3 (GRCh37 alignment, ~18,000 samples jointly called in 2016), Freeze 5 (GRCh38 alignment, ~65,000 samples jointly called in 2017), and Freeze 6 (GRCh38 alignment, ~107,000 samples jointly called in 2018). Supplementary Table 14 indicates which TOPMed study-consent groups were used in each of several different types of analyses described in this paper. Most analyses were performed on a set of 53,831 samples derived from Freeze 5 (column “General variant analyses” in Supplementary Table 14) or on a subset thereof approved for population genetic studies (column “Population genetics”). The set of 53,831 was selected from Freeze 5 by samples eligible for dbGaP sharing at the time of analysis, excluding (a) duplicate samples from the same participant (b) one member of each monozygotic twin pair; (c) samples with questionable identity or low read depth (<98% of variant sites at depth ≥10X); and (d) samples with consent types inconsistent with analyses presented here. The “unrelated” sample set consisting of 40,273 samples refers to a subset of the 53,831 samples who are unrelated with a threshold of third degree (less related than first cousins), identified using the PC-AiR method^91^.

### Identifying putative loss of function variants

Putative loss of function (pLoF) variants were identified using Loss Of Function Transcript Effect Estimator (LOFTEE) v0.3-beta^92^ and Variant Effect Predictor (VEP) v94^93^. The genomic coordinates of coding elements were based on GENCODE v29^27^. Only stop-gained, frameshift, and splice site disturbing variants annotated as high-confidence (HC) pLoF were used in the analysis. The pLoF variants with AF > 0.5% or within regions masked due to poor accessibility (Supplementary Information 1.2).

We evaluated enrichment and depletion of pLoF variants (AF < 0.5%) in gene sets (i.e. terms) from Gene Ontology (GO)^94,95^. For each gene annotated with a particular GO term we computed number of pLoF variants per protein coding base pair, *L*, and proportion of singletons, *S*. Then, we tested for lower/higher average *L* and *S* in a GO term using bootstrapping (1,000,000 samples) with adjustment for protein-coding gene length (CDS): (1) sort all genes by their CDS length in ascending order and divide them into equal-size bins (1,000 genes each); (2) count how many genes from a GO term are in each bin; (3) from each bin, sample with replacement same number of genes and compute average *L* and *S*; (4) count how many times sampled *L* and *S* were lower/higher than observed values. The p-values were computed as the proportion of bootstrap samples exceeding observed values. The fold change of average *L* and *S* was computed as a ratio of observed values to the average of sampled values. We tested 12,563 GO terms, those which included >1 gene. The p-value significance threshold was thus ~2×10^−6^. The enrichment and depletion of pLoF variants in public gene databases was tested in a similar way.

### Sequencing depth at protein-coding regions

We compared sequencing depth at protein-coding regions in TOPMed WGS and ExAC WES. The ExAC WES depth at each sequenced base pair on human genome build GRCh37 was downloaded from the ExAC browser <u>website</u>^96^. When sequencing depth summary statistics for a base pair was missing, we assumed depth <10X for this base pair. Only protein-coding CCDS genes were analyzed and the protein-coding regions (CDS) were extracted from GENCODE v29. When analyzing ExAC sequencing depth, we used GENCODE v29 lifted to human genome build GRCh37. When comparing sequencing depth for each gene individually in TOPMed and ExAC, we used only genes present in both GRCh38 and GRCh37 versions of GENCODE v29.

### Low coverage WGS and high coverage WES in Framingham Heart Study

Investigators in the Framingham Heart Study (FHS) evaluated WGS from TOPMed in comparison with sequencing data from CHARGE Consortium WGS and Whole Exome Sequencing (WES). Supplementary Table 15 provides the counts and depth of each sequencing effort. The overlap of these three groups is 430 FHS study participants, on whom we report here. We use a subset of 263 unrelated study participants to calculate the numbers of singletons and doubletons, minor allele frequency (MAF), heterozygosity, and all rates, to avoid bias from the family structure. Supplementary Information section 1.3 provides detailed description of the comparison results.

### Identification of *CYP2D6* alleles using Stargazer’s genotyping pipeline

Details of the Stargazer genotyping pipeline were described previously^50^. Briefly, SNVs and indels in *CYP2D6* were assessed from a VCF file generated using GATK-HaplotypeCaller^97^. The VCF file was phased using the program Beagle^98^ and the 1000 Genomes Project haplotype reference panel. Phased SNVs and indels were then matched to star alleles. In parallel, read depth was calculated from BAM files using GATK-DepthOfCoverage^97^. Read depth was converted to copy number by performing intra-sample normalization^50^. Following normalization, SVs were assessed by testing all possible pairwise combinations of pre-defined copy number profiles against the sample’s observed copy number profile. For novel SVs, breakpoints were statistically inferred using changepoint^99^. Information regarding novel SVs was stored and used to identify subsequent SVs in copy number profiles. Output data included individual diplotypes, copy number plots, and a VCF of SNVs and indels that were not used to define star alleles.

### Novel genetic variants in unmapped reads

Analysis of unmapped reads was performed using ~18,000 samples from TOPMed data Freeze 3. From each sample, we extracted and filtered all read-pairs with at least one umapped mate and used them to discover human sequences absent from the reference. The pipeline included four steps: (i) per-sample *de novo* assembly of unmapped reads and selection of hominid contigs using the *Pan paniscus*, *Pan troglodyte*s, *Gorilla gorilla* and *Pongo abelii* genome references; (ii) hominid-reference-based merging and scaffolding of sequences pooled together from all samples; (iii) reference placement and breakpoint calling; and (iv) variant calling. The detailed description of each step is provided in Supplementary Information section 1.4.

### Genome-wide distribution of genetic variation

#### Contiguous segment analysis

We excluded indels and multi-allelic variants, and categorized remaining variants as common (AF≥0.005) or rare (AF<0.005), and as coding or noncoding based on protein coding exons from Ensembl 94^100^. Variant counts were analyzed across 2,739 non-empty (i.e. with at least one variant) contiguous 1 Mbp chromosomal segments, and counts in segments at the end of chromosomes with length *L*<10^6^ bp were scaled up proportionally by the factor 10^6^ x *L*^−1^. For each segment, the coding proportion, *C*, was calculated as the proportion of bases overlapping protein coding exons. The distribution of *C* is fairly narrow, with 80% of segments having *C*≤ 0.0195, 99% of segments have *C*≤0.067, and only 3 segments having *C*≥0.10. Due to the significant negative correlation between *C* and the number of variants in a segment, and potential mapping effects, we use linear regression to adjust the variant counts per segment according to the model count = ß x *C* + *A* + count_adj, where *A* is the proportion of segment bases overlapping accessibility mask (Supplementary Information 1.2). Unless otherwise noted, we present analyses and results that use these adjusted count values.

#### Concatenated segment analysis

Distinct base classifications were defined by both coding and noncoding annotation (based on Ensembl 94^100^) and CADD in silico prediction scores^101^ (downloaded from the CADD server for all possible SNVs). For each base we used maximum possible CADD score (when using the minimum CADD score, results were qualitatively the same). Bases beyond the final base with a CADD score per chromosome were excluded. This resulted in six distinct types of concatenated segments: high (CADD≥20), intermediate (10≤CADD<20), and low (CADD<10) CADD scores for coding and similarly for non-coding. Common (AF≥0.005) and rare (AF<0.005) variant counts were then calculated across these concatenated segments. Multi-allelic variants and those in regions masked due to accessibility were excluded. Counts in segments at the end of chromosomes were scaled up as in the contiguous analysis.

#### Multi-nucleotide mutations

##### Data

From the TOPMed Freeze 5 dataset, we selected a subset of 1,000 unrelated individuals of African ancestry and 1,000 unrelated individuals of European ancestry, based on the global ancestry proportions inferred using RFMIX^102^. In each sample of 1,000 individuals, we recalculated the allele counts of each SNV and extracted SNVs that were singletons within that sample. Note that a singleton defined here is not necessarily a singleton in the entire TOPMed freeze 5 dataset. We chose to limit the size of each sample to N=1,000 for two reasons: first, to ensure each subsample shared homogenous ancestry, and second, to limit the incidence of recurrent mutations at hypermutable sites, which can alter the underlying mutational spectrum of singleton SNVs in large samples^103^.

##### Mixture model parameter estimation

For each individual *i*, we collected the set of *S* singletons unique to that individual (with singleton status determined relative to other individuals from the same population subsample). Assuming singletons occur independently at a constant rate ϕ_*i*_, we can model the probability of observing *S* singletons in individual *i* as a Poisson variable with mean ϕ_*i*_*G*,

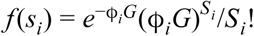

where *G* is the size in bp of the mappable autosomal regions.

Then, for singleton *j*, let *d*_*i,j*_ be the distance in base pairs to its nearest neighboring singleton in individual *i*. These distances thus follow an exponential distribution with rate θ_*i*_ = 1/ϕ_*i*_:

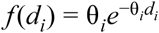

Now suppose the set of *S*_*i*_ singletons are generated by *K* > 1 independent Poisson processes, each with a different rate. Then the distribution of inter-singleton distances across all *S*_*i*_ singletons is parameterized as a mixture of *K* exponential component distributions, given by:

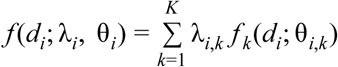

where θ_*i*,1_ < θ_*i*,2_ < … < θ_*i*,*K*_ and λ_*i*,*k*_ = *S*_*i*,*k*_/*S*_*i*_ is the proportion of singletons resulting from process *k*, such that 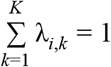.

We estimate the parameters of this mixture (λ_*i*,1_, …, λ_*i*,*K*_, θ_*i*,1_, …, θ_*i*,*K*_) using the expectation-maximization (EM) algorithm as implemented in the mixtools R package^104^. To identify an optimal number of mixture components, we iteratively fit mixture models for increasing values of K and calculated the log-likelihood of observed data D given the parameter estimates 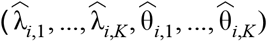, stopping at K components if the p-value of the likelihood ratio test between K-1 and K components was >0.01 (chi-squared test with 2 degrees of freedom). The goodness-of-fit plateaued at four components for the majority of individuals, so we used the 4-component parameter estimates from each individual in all subsequent analyses.

Now let *k*_*i,j*_ indicate which of the four processes generated singleton *j* in individual *i*. We calculated the probability of being generated by process *k* as:

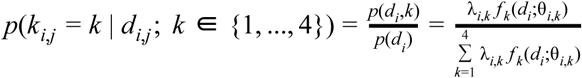

Where we then classified the process-of-origin for each singleton per the following optimal decision rule:

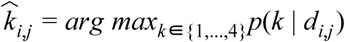

### Identification of cluster class 2 hotspots

After assigning singletons to the most likely cluster class, we counted the frequencies in non-overlapping 1Mb windows throughout the genome, and defined hotspots as the top 5% of 1Mb bins containing the most singletons within each ancestry group.

### Simulations

To quantify the effects of external branch length heterogeneity on singleton clustering patterns, we simulated singletons under the following model. First, we can assume the number *N*_*i*_ of singletons in individual i follow a Poisson(ϕ*_i_*) distribution, where 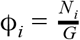(here, G indicates the total number of mutable bases in the mappable autosomal regions of the reference genome). Consequently, the distances between successive singletons in individual i are expected to follow an exponential distribution with rate θ_*i*_ = 1/ϕ_*i*_. For each individual i, we randomly drew *N*_*i*_ inter-singleton distances from the corresponding *exp*(θ_*i*_) probability distribution.

We used msprime^105^ to simulate 2,000 European chromosomes (100Mbp in length) using a demographic model with parameter estimates reported by Fu *et al.* 2013^12^. We performed simulations using a per-site, per-generation mutation rate ranging from 1×10^−8^ to 2×10^−8^. Because our aim was to compare these simulated singletons to unphased singletons in the TOPMed data, we randomly assigned each of the 2,000 haploid samples into one of 1,000 diploid pairs, and recalculated the inter-singleton distances per diploid sample, ignoring the chromosome on which each simulated singleton originated.

### Evolutionary genetics of diverse ancestry individuals

#### Rare variant sharing

We used 39,722 unrelated individuals that had consent for population genetics research. Each individual was grouped into their TOPMed study, except for individuals from the AFGen project, which were treated as one study (Supplementary Tables 1 and 2). FHS and ARIC individuals, which overlapped with the AFGen project, remained in their respective studies and were not grouped into the AFGen project. Individuals for whom self-described ancestry was either missing or “other” were removed from the analysis. We then removed all indels, multi-allelic variants, and singletons from the remaining 39,168 individuals. Each study was then split by self-described ancestry. We excluded studies that had <19 samples from the analysis, however all 39,168 samples were used to define singleton filtering. We used the Jaccard Index^106^, *J*, to determine the intersection of rare variants (2 ≤ sample count ≤ 100) between two individuals divided by the union of that pairs’ rare variants, where sample count are the number of individuals with either heterozygote or homozygote. We then determined the average *J* value between and within each study.

To confirm that *J* is not biased by sample size, we randomly sampled 500 individuals from each of two European (AFGen, FHS) and two African (COPDGene, JHS) ancestry studies in TOPMed data Freeze 3, without replacement. We then recalculated *J* between and within these randomly sampled studies from AC = 2 to 100, with AC calculated from these 2,000 individuals.

#### Identity-by-descent sharing

We used the RefinedIBD program^107^ to call segments of IBD having length ≥2 cM on the autosomes using passing SNVs with MAF>5%. All 53,831 samples were included in this analysis, and we used genotype data which had been phased with Eagle2^88^. Since IBD LOD scores are often deflated in populations with strong founding bottlenecks, such as the Amish, we used a LOD score threshold of 1.0 instead of the default 3.0. To account for possible phasing and genotyping errors, we filled gaps between IBD segments for the same pair of individuals if the gap had length at most 0.5 cM and at most one discordant genotype. As a result of the lower LOD threshold, regions with low variant density can have an excess of apparent IBD segments. We therefore identified regions with highly elevated levels of detected IBD using the procedure of Browning and Browning (2015)^108^, and removed any IBD segments falling wholly within these regions.

We divided the data by study and by self-identified ancestry within study. In the analyses of IBD sharing levels and recent effective size, we did not include studies without appropriate consent or ancestry groups with <80 individuals within a study. We calculated the total length of IBD segments for each pair of individuals, and we averaged these totals within each ancestry group in a study and between each pair of ancestry-by-study groups. We also estimated recent effective population size for each group using IBDNe^108^.

#### Demographic estimation under selection at linked sites

We selected 2,416 samples from the TOPMed data Freeze 3 which (a) had a high percentage of European ancestry; (b) were unrelated; and (c) gave consent for performing population genetics research. More detailed information about ancestry estimation and filters is provided in Supplementary Information 1.7.

We also performed several steps to filter the genome for high-quality neutral sites, which were based off of the ascertainment scheme used by Torres et al. 2018^109^ (Supplementary Information 1.7). After filtering, positions in the genome were annotated for how strongly affected they are by selection at linked sites by using the background selection (BGS) coefficient McVicker’s *B* statistic^65^. We used all sites annotated with a *B* value for performing general analyses. However, when performing demographic inference, we limited our analyses to regions of the genome within the top 1% of the genome-wide distribution of *B* (*B* >= 0.994). These sites correspond to regions of the genome inferred to be under the weakest amount of BGS (i.e., under the weakest effects of selection at linked sites). Sites in the genome were also polarized to ancestral and derived states using ancestral annotations called with high-confidence from the GRCh37 e71 ancestral sequence. After keeping only polymorphic di-allelic sites, we had 20,324,704 sites, 191,631 with *B* >= 0.994. We also identified 91,177 fourfold degenerate synonymous sites that were polymorphic (di-allelic) and had high-confidence ancestral/derived states.

We performed demographic inference with the *moments*^*110*^ program by fitting a model of exponential growth with three parameters (*N*_*Eur0*_, *N*_*Eur*_, *T*_*Eur*_) to the site-frequency spectrum (SFS). This included two free parameters: the starting time of exponential growth (*T*_*Eur*_) and the ending population size after growth (*N*_*Eur*_). The ancestral size parameter (i.e, the population size when growth begins), *N*_*Eur0*_, was kept constant in our model such that the relative starting size of the population was always 1. We applied the inference procedure to either fourfold degenerate sites or sites with *B* >= 0.994. The SFS used for inference was unfolded and based on the polarization step described above. The inference procedure was fit using sample sizes (2N) of 1,000, 2,000, 3,000, 4,000, and 4,832 chromosomes. To convert the scaled genetic parameters output by the inference procedure to physical units, we used the resulting theta (also inferred by *moments*) and a mutation rate of 1.66 x 10^−8 111^ to generate corresponding effective population sizes (*N*_*e*_). To convert generations to years, we assumed a generation time of 25 years. 95% confidence intervals were generated by resampling the SFS 1,000 times and using the Godambe Information Matrix to generate parameter uncertainties^112^. A more detailed description is available in Supplementary Information 1.7.

#### Selection

We used 8,377 unrelated individuals selected from the TOPMed data Freeze 3 for which we had consent for population genetic analyses (Supplementary Table 16). We assigned each individual to one of six population labels using k-means clustering on the first 7 PCs as described in Supplementary Information 1.8. Then, we analyzed each population separately. Only bi-allelic sites with unambiguous ancestral state, inferred using WGSA pipeline, were used. Sites near chromosome boundaries and with minor allele frequency <0.05 were also excluded (Supplementary Table 17). We used the program selscan^113^ to perform all iHS scans^66^ in each population using default parameters. Then we normalized raw iHS scores within 20 frequency bins. We followed Voight et al. 2006^66^ to find regions with the largest proportion of extreme iHS scores -- putatively selected regions (Supplementary Information 1.8). A full list of genes found in significant regions, and the populations in which they were found significant, is given in Supplementary File 2. A full list of regions identified as significant in each population can be found in Supplementary Files 3-8.

### TOPMed Imputation Panel

#### Construction

We divided each autosomal chromosome and the X chromosome into overlapping chunks (with chunk size 1Mb each and with 0.1Mb overlap between consecutive chunks), and then phased each of the chunks using Eagle v2.4^88^. We removed all singleton sites and compressed the haplotype chunks into m3vcf format^114^. Afterwards, we ligated the compressed haplotype chunks for each chromosome to generate the final reference panel.

#### Evaluation of imputation accuracy

For all TOPMed individuals, genetic ancestries were estimated using the top four principal components (PCs) projected onto the PC space of 938 Human Genome Diversity Project (HGDP) individuals using verifyBamID2^115^. For each TOPMed individual, we identified the 20 closest individuals from 2,504 individuals from the 1000 Genomes Project Phase 3 (1000G) based on Euclidean distances in the PC space estimated by verifyBamID2. If all of the 20 closest 1000G individuals belonged to the same super-population - among African (AFR), Admixed American (AMR), East Asian (EAS), European (EUR), South Asian (SAS) - we estimated that the TOPMed individual also belongs to that super-population. Among the 60,039 reference panel individuals, 55,365 (92%) were assigned to a super-population, with the following breakdown: AFR=14,764, AMR=4,347, EUR=31,686, EAS=4,429, SAS=139. Of 5,504 additionally sequenced individuals for the BioMe study but not included in the TOPMed reference panel, 4,725 were assigned to a single super-population using this method. We randomly selected 100 individuals from each super-population, and selected markers on chromosome 20 present on the Illumina HumanOmniExpress (8v1-2_A) array. The selected genotypes were phased with Eagle 2.4.1^88^, using the 1000G (n=2,504), Haplotype Reference Consortium (HRC, n=32,470)^8^, and TOPMed (n=60,039) reference panels. The phased genotypes were imputed using Minimac4^116^ from each reference panel, and the imputation accuracy was estimated as the squared correlation coefficient (r^2^) between the imputed dosages and the genotypes calls from the sequence data. The allele frequencies were estimated among all TOPMed individuals estimated to belong to the same super-population, and the r^2^ values were averaged across variants in each MAF category. Variants present in 100 sequenced individuals but absent from the reference panels were assumed to have r^2^=0 for the purposes of computing the average r^2^. The minimum MAF to achieve r^2^>0.3 was calculated from the average r^2^ in each MAF category by finding the MAF that crosses r^2^=0.3 using linear interpolation. The average number of rare variants (MAF<0.5%) and the fraction of imputable rare variants (r^2^>0.3) were calculated based on the number of non-reference alleles in imputed samples above and below the minimum MAF, assuming HWE.

#### Imputation of the UK Biobank to the TOPMed panel and association analyses

After phasing the UK Biobank genetic data (carried out on 81 chromosomal chunks using Eagle v2.4.), the phased data were converted from GRCh37 to GRCh38 using LiftOver^117^. Imputation was performed using Minimac4^116^.

We tested single putative loss of function (pLoF) nonsense, frameshift or essential splice site variants (determined as described previously) for association with 1,403 traits: PheCodes constructed from composites of ICD-10 codes to define cases and controls. Construction of the PheCodes has been previously described^70^. To perform the association analyses, we used a logistic mixed model test implemented in SAIGE^70^ with birth year and the top four PCs (computed from the White British subset) as covariates. For the pLoF burden tests, for each gene with at least two rare pLoF variants (N=8,636 genes) a burden variable was created in which dosages of rare pLoF variants were summed for each individual. This sum of dosages was tested for association with the 1,403 traits using SAIGE. The same covariates used in the single-variant tests were included. For both the single-variant and the burden tests we used 5.0×10^−8^ as the genome-wide significance threshold.

## Acknowledgements

Whole genome sequencing (WGS) for the Trans-Omics in Precision Medicine (TOPMed) program was supported by the National Heart, Lung and Blood Institute (NHLBI). Specific funding sources for each study and genomic center are given in Supplementary Table 18. Centralized read mapping and genotype calling, along with variant quality metrics and filtering were provided by the TOPMed Informatics Research Center (3R01HL-117626-02S1). Phenotype harmonization, data management, sample-identity QC, and general study coordination, were provided by the TOPMed Data Coordinating Center (3R01HL-120393-02S1). We gratefully acknowledge the studies and participants who provided biological samples and data for TOPMed. The full study specific acknowledgments are detailed in Supplementary Information section 2.

The UK Biobank analyses were conducted using the UK Biobank Resource under application number 24460.

Other acknowledgments are detailed in Supplementary Information section 3.

The views expressed in this manuscript are those of the authors and do not necessarily represent the views of the National Heart, Lung, and Blood Institute; the National Institutes of Health; or the U.S. Department of Health and Human Services.

## Author Contributions

T.W.B., Q.W., F.A., K.G.A., P.L.A., R.G.B., R.L.B., J.Bl., M.B., E.G.B., J.F.C., Y.I.C., M.H.C., A.Co., J.E.C., D.L.D., P.T.E., M.F., N.F., S.M.F., D.J.G., M.E.H., J.H., S.R.H., M.R.I., A.D.J., S.K., D.P.K., C.K., A.K., L.A.L., J.La., D.L., C.L., K.L.L., A.M., A.K.M., R.A.M., S.T.M., J.B.M., J.L.M., M.A.M., B.M., M.E.M., C.M., A.C.M., J.M.M., P.N., K.E.N., N.P., G.M.P., W.S.P., B.M.P., D.C.R., S.R., A.P.R., J.I.R., I.R., C.S., S.Se., V.A.S., N.L.S., N.S., K.D.T., R.S.V., S.V., D.E.W., B.S.W., S.T.W., C.J.W., D.K.A., A.E.A., K.C.B., E.B., S.Ga., R.Gi., K.M.R., S.S.R., E.S., P.Q., W.G., G.J.P., D.A.N., S.Z., J.G.W., L.A.C., C.C.L., C.E.J., R.D.H., T.D.O., and G.R.A. contributed to the conception or design of the TOPMed program and its operations. A.A., S.A., L.C.B., E.J.B., L.F.B., J.Bl., D.W.B., E.G.B., B.E.C., B.Ch., Y.I.C., M.K.C., A.Co., J.E.C., B.Cu., D.D., M.D., P.T.E., D.F., M.T.G., X.G., J.H., N.L.H., S.R.H., J.M.J., S.L.R.K., S.K., E.E.K., D.P.K., B.A.K., C.K., L.A.L., J.La., R.J.F.L., L.G., R.Ge., S.A.L., K.L.L., A.C.Y.M., R.A.M., D.D.M., S.T.M., D.A.M., B.M., S.M., C.M., A.N., N.D.P., P.A.P., W.S.P., B.M.P., D.C.R., S.R., J.I.R., S.Se., V.A.S., J.A.S., K.D.T., M.T., D.J.V.D.B., R.S.V., D.E.W., S.T.W., Y.Z., D.K.A., A.E.A., K.C.B., E.B., S.S.R., E.S., J.G.W., L.A.C., R.D.H. provided phenotypic data and/or biosamples. F.A., K.G.A., L.C.B., J.Bl., B.E.C., C.B.C., J.E.C., S.K.D., P.T.E., S.Ge., X.G., D.L., R.J.F.L., S.T.M., J.I.R., J.S., K.D.T., D.J.V.D.B., D.E.W., A.E.A., K.C.B., E.B., S.Ga., R.Gi., G.J.P., D.A.N. acquired WGS and/or other omics data. D.T., D.N.H., M.D.K., J.C., Z.A.S., R.T., S.A.G.T., A.Co., S.M.G., H.M.K., A.N.P., J.Le., S.L., X.T., B.L.B., S.D., A.E., W.E.C., D.P.L., A.C.S., T.W.B., Q.W., L.S.E., L.F., C.F., S.Ge., X.L., K.L., S.C.N., S.Sc., A.M.S., X.Z., E.B., and D.A.N. created software, processed, and/or analyzed WGS or other data for data summaries in this paper. D.T., D.N.H., M.D.K., J.C., Z.A.S., R.T., S.A.G.T., A.Co., S.D., S.Ge., S.R.B., L.A.C., C.C.L., C.E.J., R.D.H., T.D.O., G.R.A. drafted the manuscript and revised according to co-author suggestions. All authors critically reviewed the manuscript, suggested revisions as needed, and approved the final version.

