## Supplementary Text for "Sequencing of 53,831 diverse genomes from the NHLBI TOPMed Program"

1 - Department of Biostatistics and Center for Statistical Genetics, University of Michigan School of Public Health, Ann Arbor, MI; 2 - Institute for Genome Sciences, University of Maryland School of Medicine, Baltimore, MD; 3 - Program in Personalized and Genomic Medicine, University of Maryland School of Medicine, Baltimore, MD; 4 - Department of Medicine, University of Maryland School of Medicine, Baltimore, MD; 5 - Department of Computational Medicine and Bioinformatics, University of Michigan, Ann Arbor, MI; 6 - Department of Genome Sciences, University of Washington, Seattle, WA; 7 - Department of Bioengineering and Therapeutic Sciences, University of California, San Francisco, CA; 8 - Department of Biological Sciences, Auburn University, Auburn, AL; 9 - Biomedical Sciences Graduate Program, University of California, San Francisco, CA; 10 - New York Genome Center, New York, NY; 11 - Department of Biostatistics, University of Washington, Seattle, WA; 12 - Department of Biostatistics, Boston University School of Public Health, Boston, MA; 13 - Department of Medicine, Division of Medical Genetics, University of Washington, Seattle, WA; 14 - The Broad Institute of MIT and Harvard, Cambridge, MA; 15 - Massachusetts General Hospital, Boston, MA; 16 - Department of Epidemiology, Rollins School of Public Health, Emory University, Atlanta, GA; 17 - University of Alabama, Birmingham, AL; 18 - Zilber School of Public Health, University of Wisconsin Milwaukee, Milwaukee, WI; 19 - Cleveland Clinic, Cleveland, OH; 20 - Department of Medicine, Columbia University Medical Center, New York, NY; 21 - Department of Epidemiology, Columbia University Medical Center, New York, NY; 22 - Johns Hopkins University, Baltimore, MD; 23 - National Heart, Lung, and Blood Institute, National Institutes of Health, Bethesda, MD; 24 - Department of Medicine, Boston University School of Medicine, Boston, MA; 25 - Department of Epidemiology, Boston University School of Public Health, Boston, MA; 26 - Framingham Heart Study, Framingham, MA; 27 - Department of Epidemiology, University of Michigan School of Public Health, Ann Arbor, MI; 28 - Department of Human Genetics, University of Texas Rio Grande Valley School of Medicine, Brownsville, TX; 29 - South Texas Diabetes and Obesity Institute, University of Texas Rio Grande Valley School of Medicine, Brownsville, TX; 30 - Department of Biochemistry, Wake Forest School of Medicine, Winston-Salem, NC; 31 - Department of Medicine, University of Washington, Seattle, WA; 32 - Cardiovascular Health Research Unit, University of Washington, Seattle, WA; 33 - Department of Medicine, University of California, San Francisco, CA; 34 - Department of Medicine, Harvard Medical School, Boston, MA; 35 - Department of Medicine, Brigham and Women's

Hospital, Boston, MA; 36 - Department of Pediatrics, Johns Hopkins University, Baltimore, MD; 37 - Division of Pediatric Hematology, Johns Hopkins University, Baltimore, MD; 38 - Department of Medical Genetics, University of British Columbia, Vancouver, BC; 39 - Department of Pediatrics, The Institute for Translational Genomics and Population Sciences, Los Angeles Biomedical Research Institute at Harbor-UCLA Medical Center, Torrance, CA; 40 - Channing Division of Network Medicine, Department of Medicine, Brigham and Women's Hospital, Boston, MA; 41 - Department of Cardiovascular Medicine, Heart & Vascular Institute, Cleveland Clinic, Cleveland, Ohio, USA; 42 - Department of Cardiovascular and Metabolic Sciences, Lerner Research Institute, Cleveland Clinic, Cleveland, Ohio, USA; 43 - Cleveland Clinic Lerner College of Medicine of Case Western Reserve University, Cleveland, Ohio, USA; 44 - Metabolomics Platform, The Broad Institute of MIT and Harvard, Cambridge, MA; 45 - Department of Medicine, University of Mississippi Medical Center, Jackson, MS; 46 - Department of Pediatrics, University of Mississippi Medical Center, Jackson, MS; 47 - Department of Population Health Science, University of Mississippi Medical Center, Jackson, MS; 48 - Vitalant Research Institute, San Francisco, CA, USA; 49 - Department of Laboratory Medicine, UCSF, San Francisco, CA, USA; 50 - Department of Medicine, University of Illinois at Chicago, Chicago, IL; 51 - School of Medicine, University of Colorado, Aurora, CO; 52 - Mayo Clinic, Rochester, MN; 53 - McDonnell Genome Institute and Department of Genetics, Washington University, St Louis, MO; 54 - Program in Medical and Population Genetics, The Broad Institute of MIT and Harvard, Cambridge, MA; 55 - Molecular Cardiology Division, Victor Chang Cardiac Research Institute, Darlinghurst, NSW, Australia; 56 - Faculty of Medicine, University of New South Wales, Kensington, NSW, Australia; 57 - Cardiology Department, St. Vincent’s Hospital, Darlinghurst, NSW, Australia; 58 - Division of Genetic Epidemiology, Department of Medical Genetics, Molecular and Clinical Pharmacology, Medical University of Innsbruck, Innsbruck, Austria; 59 - Institute of Molecular Medicine, University of Texas Health Science Center at Houston, Houston, TX; 60 - Department of Epidemiology, University of North Carolina, Chapel Hill, NC; 61 - Institute for Biomedicine, Eurac Research, Bolzano, Italy; 62 - Department of Bioethics & Humanities, University of Washington School of Medicine, Seattle, WA; 63 - Pittsburgh Heart, Lung, Blood and Vascular Medicine Institute, University of Pittsburgh, Pittsburgh, PA; 64 - Pulmonary, Allergy and Critical Care Medicine, University of Pittsburgh, Pittsburgh, PA; 65 - Department of Medicine, University of Pittsburgh, Pittsburgh, PA; 66 - VA Boston Healthcare System, Boston, MA; 67 - Division of Sleep and Circadian Disorders, Brigham and Women's Hospital, Boston, MA; 68 - Department of Epidemiology, Tulane University, New Orleans, LA; 69 - Tulane University Translational Science Institute, Tulane University, New Orleans, LA; 70 - Department of Neurology, Boston University School of Medicine, Boston, MA; 71 - Department of Epidemiology, University of Washington, Seattle, WA; 72 - Department of Epidemiology, University of Alabama at Birmingham, Birmingham, AL; 73 - Bloodworks Northwest Research Institute, Seattle, WA; 74 - Population Sciences Branch, National Heart, Lung, and Blood Institute, National Institutes of Health, Framingham, MA; 75 - Department of Epidemiology, Vitalant Research Institute, San Francisco, CA; 76 - Department of Pediatrics, UCSF Benioff Children's Hospital, Oakland, CA; 77 - Division of Pediatric Hematology, UCSF Benioff Children's Hospital, Oakland, CA; 78 - Icahn School of Medicine at Mount Sinai, New York, NY; 79 - Marcus Institute for Aging Research/Harvard Medical School, Hebrew SeniorLife, Boston, MA; 80 - Department of Medicine, Beth Israel Deaconess Medical Center, Boston, MA; 81 - Division of Public Health Sciences, Fred Hutchinson Cancer Research Center, Seattle, WA; 82 - Department of Epidemiology, Johns Hopkins University, Baltimore, MD; 83 - Institute of Genetic Epidemiology, Medical Center - University of Freiburg, Freiburg, Germany; 84 - Department of Medicine, University of Colorado at Denver, Aurora, CO; 85 - Brigham and Women's Hospital, Boston, MA; 86 - Biostatistics and Statistics, Harvard University, Boston, MA; 87 - The Charles Bronfman Institute for Personalized Medicine, Icahn School of Medicine at Mount Sinai, New York, NY, USA; 88 - The Mindich Child Health and Development Institute, Icahn School of Medicine at Mount Sinai, New York, NY, USA; 89 - Division of Genomics and Data Science, Department of Arthritis and Clinical Immunology, Oklahoma Medical Research Foundation, Oklahoma City, OK; 90 - Beth Israel Deaconess Medical Center, Boston, MA; 91 - Center for Public Health Genomics, University of Virginia, Charlottesville, VA; 92 - Department of Public Health Sciences, University of Virginia, Charlottesville, VA; 93 - Department of Medicine/Clinical and Translational Epidemiology Unit, Massachusetts General Hospital, Boston, MA; 94 - Department of Medicine, Metabolism Program, Massachusetts General Hospital, Boston, MA; 95 - Metabolism Program, The Broad Institute of MIT and Harvard, Cambridge, MA; 96 - Department of Medicine, Johns Hopkins University, Baltimore, MD; 97 - Cardiovascular Medicine, University of Massachusetts Medical School, Worcester, MA, USA; 98 - International Health Institute, Brown University, Providence, RI; 99 - Department of Epidemiology, Brown University, Providence, RI; 100 - Department of Anthropology, Brown University, Providence, RI; 101 - Division Of General Internal Medicine, Massachusetts General Hospital, Harvard Medical School, The Broad Institute of MIT and Harvard, Boston, MA; 102 - University of Arizona; 103 - Geriatrics Research and Education Clinical Center, Baltimore Veterans Administration Medical Center, Baltimore, MD; 104 - Texas Cardiac Arrhythmia Institute, St. David's Medical Center, Austin, TX, USA; 105 - Department of Internal Medicine, Dell Medical School, Austin, TX, USA; 106 - Human Genetics Center, Department of Epidemiology, Human Genetics, and Environmental Sciences, School of Public Health, University of Texas Health Science Center at Houston, Houston, TX; 107 - Cardiovascular Research Center, Massachusetts General Hospital, Boston, MA; 108 - Center for Genomic Medicine, Massachusetts General Hospital, Boston, MA; 109 - Department of Laboratory Medicine and Pathology, University of Minnesota, Minneapolis, MN; 110 - Division of Cardiology, Department of Medicine, Johns Hopkins University, Baltimore, MD; 111 - Department of Health Services, University of Washington, Seattle, WA; 112 - Kaiser Permanente Washington Health Research Institute, Seattle, WA; 113 - Division of Biostatistics, Washington University in St. Louis, St. Louis, MO; 114 - Vanderbilt University, Nashville, TN; 115 - Department of Medicine, The Institute for Translational Genomics and Population Sciences, Los Angeles Biomedical Research Institute at Harbor-UCLA Medical Center, Torrance, CA; 116 - Department of Biostatistics, Johns Hopkins Bloomberg School of Public Health, Baltimore, MD; 117 - Precision Medicine Center, Seoul National

University Bundang Hospital, Seongnam, Republic of Korea; 118 - Macrogen Inc., Seoul, Republic of Korea; 119 - Gong Wu Genomic Medicine Institute, Seoul National University Bundang Hospital, Seongnam, Republic of Korea; 120 - Glenn Biggs Institute for Alzheimer's and Neurodegenerative Diseases, University of Texas Health Sciences Center at San Antonio, San Antonio, TX; 121 - Department of Pediatrics, Division of Hematology/Oncology, Baylor College of Medicine, Houston, TX; 122 - Seattle Epidemiologic Research and Information Center, Veterans Administration Puget Sound Health Care System, Seattle, WA; 123 - Survey Research Center, Institute for Social Research, University of Michigan, Ann Arbor, MI; 124 - Division of Epidemiology and Community Health, School of Public Health, University of Minnesota, Minneapolis, Minnesota, USA; 125 - Duke University; 126 - University of Vermont, Burlington, VT; 127 - University of Southern California, Center for Genetic Epidemiology, Department of Preventive Medicine, Los Angeles, CA; 128 - Illumina Clinical Genomics Laboratory Services, San Diego, CA; 129 - Department of Psychology, University of Minnesota, Minneapolis, MN; 130 - Department of Human Genetics, Graduate School of Public Health, University of Pittsburgh, Pittsburgh, PA; 131 - Department of Biostatistics, Graduate School of Public Health, University of Pittsburgh, Pittsburgh, PA; 132 - Department of Internal Medicine, University of Michigan, Ann Arbor, MI; 133 - Department of Human Genetics, University of Michigan, Ann Arbor, MI; 134 - Department of Biostatistics, University of Michigan, Ann Arbor, MI; 135 - Department of Epidemiology, University of Kentucky, Lexington, KY; 136 - Duke Molecular Physiology Institute, Duke University Medical Center, Durham, NC; 137 - University of Texas Health Science Center at Houston, Houston, TX; 138 - Baylor College of Medicine Human Genome Sequencing Center, Houston, TX; 139 - Northwest Genomics Center, Seattle, WA; 140 - Brotman Baty Institute, Seattle, WA; 141 - Department of Psychiatry, University of Michigan, Ann Arbor, MI; 142 - Department of Physiology and Biophysics, University of Mississippi Medical Center, Jackson, MS; 143 - Department of Human Genetics, McGill University, Montreal, Canada

* These authors contributed equally to this work
† https://www.nhlbiwgs.org/topmed-banner-authorship; See “Additional Authors from the Trans-Omics for Precision Medicine Program” for full banner author list (excluding primary authors above)

#### Additional Authors from the Trans-Omics for Precision Medicine Program

Namiko Abe[144], Laura Almasy[145], Seth Ament[146], Peter Anderson[147], Pramod Anugu[148], Deborah Applebaum-Bowden[149], Dan Arking[150], Tim Assimes[151], Dimitrios Avramopoulos[150], Emily Barron-Casella[150], Terri Beaty[150], Diane Becker[150], Ferdouse Begum[150], Amber Beitelshees[146], Marcos Bezerra[152], Joshua Bis[147], Ingrid Borecki[147], Russell Bowler[153], Ulrich Broeckel[154], Jai Broome[147], Karen Bunting[144], Jonathan Cardwell[155], Cara Carty[156], Richard Casaburi[157], Mark Chaffin[158], Christy Chang[146], Daniel Chasman[159], Sameer Chavan[155], Bo-Juen Chen[144], Wei-Min Chen[160], Lee-Ming Chuang[161], Elaine Cornell[162], Carolyn Crandall[157], James Crapo[153], Jeffrey Curtis[163], Coleen Damcott[146], Sean David[151], Colleen Davis[147], Michael DeBaun[164], Ranjan Deka[165], Scott Devine[146], Ron Do[166], Qing Duan[167], Ravi Duggirala[168], Peter Durda[162], Charles Eaton[169], Lynette Ekunwe[148], Charles Farber[160], Leanna Farnam[159], Tasha Fingerlin[170], Matthew Flickinger[163], Mao Fu[146], Lucinda Fulton[171], Yan Gao[148], Margery Gass[172], Bruce Gelb[166], Xiaoqi (Priscilla) Geng[163], Chris Gignoux[151], David Glahn[173], Da-Wei Gong[146], Harald Goring[174], C. Charles Gu[171], Yue Guan[146], Jeff Haessler[175], Nicola Hawley[173], Ben Heavner[176], David Herrington[177], Craig Hersh[178], Bertha Hidalgo[179], James Hixson[180], John Hokanson[155], Elliott Hong[146], Karin Hoth[181], Chao (Agnes) Hsiung[182], Haley Huston[183], Chii Min Hwu[184], Rebecca Jackson[185], Deepti Jain[147], Min A Jhun[163], Craig Johnson[147], Rich Johnston[186], Kimberly Jones[150], Robert Kaplan[187], Sekar Kathiresan[158], Laura Kaufman[159], Alyna Khan[147], Greg Kinney[155], Holly Kramer[188], Stephanie Krauter[147], Christoph Lange[189], Ethan Lange[155], Cecelia Laurie[147], Meryl LeBoff[159], Jiwon Lee[159], Seunggeun Shawn Lee[163], Wen-Jane Lee[184], David Levine[147], Joshua Lewis[146], Yun Li[167], Honghuang Lin[190], Simin Liu[191], Yongmei Liu[177], James Luo[192], Michael Mahaney[193], Barry Make[150], JoAnn Manson[159], Lauren Margolin[158], Lisa Martin[194], Susan Mathai[155], Patrick McArdle[146], Merry-Lynn McDonald[179], Sean McFarland[195], Hao Mei[148], Nancy Min[148], Ryan L Minster[196], Solomon Musani[197], Stanford Mwasongwe[148], Josyf C Mychaleckyj[160], Girish Nadkarni[166], Rakhi Naik[150], Take Naseri[198], Sergei Nekhai[199], Heather Ochs-Balcom[200], James Pankow[201], Margaret Parker[159], Afshin Parsa[146], Sara Penchev[153], Juan Manuel Peralta[168], Marco Perez[151], James Perry[146], Ulrike Peters[202], Lawrence S Phillips[186], Sam Phillips[147], Toni Pollin[146], Julia Powers Becker[203], Meher Preethi Boorgula[155], Michael Preuss[166], Dmitry Prokopenko[195], Dandi Qiao[159], Zhaohui Qin[186], Nicholas Rafaels[155], Laura Raffield[167], Laura Rasmussen-Torvik[204], Aakrosh Ratan[160], Robert Reed[146], Elizabeth Regan[153], Muagututi‘a Sefuiva Reupena[205], Carolina Roselli[158], Pamela Russell[155], Sarah Ruuska[183], Kathleen Ryan[146], Ester Cerdeira Sabino[206], Phuwanat Sakornsakolpat[159], Shabnam Salimi[146], Steven Salzberg[150], Kevin Sandow[207], Vijay Sankaran[195], Christopher Scheller[163], Ellen Schmidt[163], Karen Schwander[171], David Schwartz[155], Frank Sciurba[196], Christine Seidman[208], Jonathan Seidman[208], Aniket Shetty[155], Wayne Hui-Heng Sheu[184], Brian Silver[209], Josh Smith[147], Tanja Smith[144], Sylvia Smoller[187], Beverly Snively[210], Tamar Sofer[159], Elizabeth Streeten[146], Yun Ju Sung[171], Jody Sylvia[159], Adam Szpiro[147], Carole Sztalryd[146], Hua Tang[151], Margaret Taub[150], Simeon Taylor[146], Lesley Tinker[156], David Tirschwell[147], Hemant Tiwari[179], Michael Tsai[201], Dhananjay Vaidya[150], Peter VandeHaar[163], Tarik Walker[155], Robert Wallace[181], Avram Walts[155],

Emily Wan[159], Fei Fei Wang[147], Heming Wang[211], Karol Watson[157], Kayleen Williams[147], L. Keoki Williams[212], Carla Wilson[159], Huichun Xu[146], Lisa Yanek[150], Ivana Yang[155], Rongze Yang[146], Norann Zaghloul[146], Maryam Zekavat[158], Snow Xueyan Zhao[153], Wei Zhao[163], Xiuwen Zheng[147], Degui Zhi[180], Xiang Zhou[163], Xiaofeng Zhu[213]

144 - New York Genome Center, New York, NY, US; 145 - Children's Hospital of Philadelphia, University of Pennsylvania, Philadelphia, PA, US; 146 - University of Maryland, Baltimore, MD, US; 147 - University of Washington, Seattle, WA, US; 148 - University of Mississippi, Jackson, MS, US; 149 - National Institutes of Health, Bethesda, MD, US; 150 - Johns Hopkins University, Baltimore, MD, US; 151 - Stanford University, Stanford, CA, US; 152 - Fundação de Hematologia e Hemoterapia de Pernambuco - Hemope, Recife, BR; 153 - National Jewish Health, Denver, CO, US; 154 - Medical College of Wisconsin, Milwaukee, WI, US; 155 - University of Colorado at Denver, Denver, CO, US; 156 - Women's Health Initiative, Seattle, WA, US; 157 - University of California, Los Angeles, CA, US; 158 - Broad Institute, Cambridge, MA, US; 159 - Brigham & Women's Hospital, Boston, MA, US; 160 - University of Virginia, Charlottesville, VA, US; 161 - National Taiwan University, National Taiwan University Hospital, Taipei, TW; 162 - University of Vermont, Burlington, VT, US; 163 - University of Michigan, Ann Arbor, MI, US; 164 - Vanderbilt University, Nashville, TN, US; 165 - University of Cincinnati, Cincinnati, OH, US; 166 - Icahn School of Medicine at Mount Sinai, New York, NY, US; 167 - University of North Carolina, Chapel Hill, NC, US; 168 - University of Texas Rio Grande Valley School of Medicine, Edinburg, TX, US; 169 - Brown University, Providence, RI, US; 170 - National Jewish Health, Center for Genes, Environment and Health, Denver, CO, US; 171 - Washington University in St Louis, St Louis, MO, US; 172 - Fred Hutchinson Cancer Research Center, Seattle, WA, US; 173 - Yale University, New Haven, CT, US; 174 - University of Texas Rio Grande Valley School of Medicine, San Antonio, TX, US; 175 - Fred Hutchinson Cancer Research Center, Women's Health Initiative, Seattle, WA, US; 176 - University of Washington, Biostatistics, Seattle, WA, US; 177 - Wake Forest Baptist Health, Winston-Salem, NC, US; 178 - Brigham & Women's Hospital, Channing Division of Network Medicine, Boston, MA, US; 179 - University of Alabama, Birmingham, AL, US; 180 - University of Texas Health at Houston, Houston, TX, US; 181 - University of Iowa, Iowa City, IA, US; 182 - National Health Research Institute Taiwan, Institute of Population Health Sciences, NHRI, Miaoli County, TW; 183 - Blood Works Northwest, Seattle, WA, US; 184 - Taichung Veterans General Hospital Taiwan, Taichung City, TW; 185 - Ohio State University Wexner Medical Center, Internal Medicine, DIvision of Endocrinology, Diabetes and Metabolism, Columbus, OH, US; 186 - Emory University, Atlanta, GA, US; 187 - Albert Einstein College of Medicine, New York, NY, US; 188 - Loyola University, Public Health Sciences, Maywood, IL, US; 189 - Harvard School of Public Health, Biostats, Boston, MA, US; 190 - Boston University, Boston, MA, US; 191 - Brown University, Women's Health Initiative, Epidemiology, Providence, RI, US; 192 - National Heart, Lung, and Blood Institute, National Institutes of Health, Bethesda, MD, US; 193 - University of Texas Rio Grande Valley School of Medicine, Brownsville, TX, US; 194 - George

Washington University, Washington, DC, US; 195 - Harvard University, Cambridge, MA, US; 196 - University of Pittsburgh, Pittsburgh, PA, US; 197 - University of Mississippi, Medicine, Jackson, MS, US; 198 - Ministry of Health, Government of Samoa, Apia, WS; 199 - Howard University, Washington, DC, US; 200 - University at Buffalo, Buffalo, NY, US; 201 - University of Minnesota, Minneapolis, MN, US; 202 - Fred Hutchinson Cancer Research Center, University of Washington, Seattle, WA, US; 203 - University of Colorado at Denver, Medicine, Denver, CO, US; 204 - Northwestern University, Chicago, IL, US; 205 - Lutia I Puava Ae Mapu I Fagalele, Apia, WS; 206 - Universidade de Sao Paulo, Faculdade de Medicina, Sao Paulo, BR; 207 - Los Angeles Biomedical Research Institute, Los Angeles, CA, US; 208 - Harvard Medical School, Boston, MA, US; 209 - UMass Memorial Medical Center, Worcester, MA, US; 210 - Wake Forest Baptist Health, Biostatistical Sciences, Winston-Salem, NC, US; 211 - Brigham & Women's Hospital, Partners.org, Boston, MA, US; 212 - Henry Ford Health System, Detroit, MI, US; 213 - Case Western Reserve University, Cleveland, Ohio, US

**Table of Contents**

**1 Supplementary Text** **11**
1.1 TOPMed program description 11
1.1.1 Organizational components 11
1.1.2 Study designs 12
1.1.3 Phenotypic data 13
1.1.4 Participant diversity 14
1.1.5 Multi-omics assays 15
1.1.6 Resources available 17
1.2 Whole Genome Sequencing Accessibility Mask 18
1.3 Comparison to low coverage WGS and high coverage WES in Framingham Heart Study 18
1.4 Novel genetic variants in unmapped reads 21
1.4.1 Assembly of hominid non-human-reference sequences from unmapped reads 21
1.4.2 Reference-based contig merging and scaffolding 21
1.4.3 Reference placement and breakpoint calling 22
1.4.4 Presence/absence calling 23
1.5 Site Frequency Spectrum 23
1.6 Admixture 24
1.7 Demographic estimation under selection at linked sites 25
1.7.1 Sample selection 25
1.7.2 Site filtering/ascertainment 25
1.7.3 Demographic inference 27
1.8 Selection 28
**2 Study Acknowledgments** **30**

**3 Other Acknowledgments** **40**

**4 References** **43**

### 1 Supplementary Text

#### 1.1 TOPMed program description

##### 1.1.1 Organizational components

TOPMed has several organizational components: the NHLBI program office, an External Advisory Panel, an Informatics Research Center (IRC), a Data Coordinating Center (DCC), several Omics Centers (Supplementary Table 13), several internal committees (including Executive; Steering; Analysis; Publications; and Ethical, Legal, and Social Implications (ELSI)), 32 investigator-led Working Groups, and >80 participating studies. The IRC (Gonçalo Abecasis, Principal Investigator) is located in the Biostatistics Department at the University of Michigan; it has primary responsibility for WGS data processing, including read alignments harmonized across sequencing centers, variant discovery, and genotype calling. The DCC (Ken Rice, Principal Investigator) is located in the Biostatistics Department at the University of Washington; it has primary responsibility for coordinating data flow and activities within the TOPMed program, as well as coordination with external entities, such as the NHLBI program office, dbGaP, and other genomics programs. Both the IRC and DCC perform quality control, develop analytical methods, perform analyses, and advise investigators on analytical issues.

The Working Groups are each focused on a particular phenotypic area such as atherosclerosis or asthma, or other general scientific area such as analysis methods or population genetics. Much of the scientific work of the program is performed in these Working Groups where TOPMed investigators collaborate on designing, performing, and reporting multi-study analyses for publication. TOPMed data are being shared with the general scientific community through periodic releases on dbGaP, after completion of quality control and dbGaP curation (in total, approximately 6 to 12 months after release to study investigators).

A key priority for the TOPMed program is the development of methodology and software for WGS data in genotype-phenotype association and other types of analyses. Therefore, several methods development projects[1] are being supported in areas such as omics data integration and annotation, pedigree based analyses, machine learning, and high performance statistical modeling with a focus on heart, lung, blood and sleep (HLBS) diseases and risk factors. TOPMed groups have been supported to develop and maintain association analysis pipelines based on the GENESIS[2] and EPACTS[3] statistical software suites. Furthermore, training of young investigators has been supported by periodic analysis workshops[4] focused on TOPMed data.

##### 1.1.2 Study designs

The TOPMed program consists of multiple “Projects”, composed of one or more “Parent” studies, each of which had previously recruited participants and obtained consent, phenotypic data, and biosamples. Projects were considered for inclusion in TOPMed based largely on peer-reviewed responses to an NHLBI Funding Opportunity Announcement[5] for X01 applications. Study selection criteria included ancestral and ethnic diversity; richness of HLBS phenotypes and environmental risk factors; and design features facilitating the detection of rare variant effects. Within each study, participants were selected for WGS according to similar criteria, described more specifically in the TOPMed dbGaP accession for each study. In all cases, participants provided informed consent for genetic studies and data sharing with the scientific community via controlled access.

TOPMed includes multiple study designs with different strengths: (1) prospective cohorts provide broad phenotypic characterization and enable analyses of incident disease, longevity, and mortality; (2) cross-sectional or case-control studies increase precision in comparison of risk factors between cases and controls; (3) families and population isolates facilitate the study of rare variants by enrichment of private variants[6–8]; (4) family-based designs provide robustness to population structure[9]; and (5) case-only studies optimize analyses of disease severity and response to treatment.

Supplementary Tables 1 and 2 provide summaries of Projects and Parent studies in the first two phases of TOPMed, for which data from ~55,000 samples have been released on dbGaP. Projects and Parent studies from all five Phases are described on the TOPMed website[10]. The following examples illustrate some of the variety in the design of TOPMed Projects and their component studies.

1. “The Jackson Heart Study” (JHS; Adolfo Correa, PI) Project derives from a single Parent study of African Americans, with a prospective cohort design and extensive clinical characterization across many different phenotypic areas.

2. “The Barbados Asthma Genetics Study” (BAGS; Kathleen Barnes, PI) Project derives from a single family-based genetic Parent study focused on asthma. Pediatric probands with asthma were initially recruited through local clinics, followed by recruitment of parents and other family members, and expansion to independent asthma cases and controls.
3. The “Atrial Fibrillation Genetics Consortium” (Patrick Ellinor, PI) Project is a consortium of several Parent studies of largely European ancestry, each contributing early-onset atrial fibrillation cases. Controls were derived from other TOPMed Projects.

##### 1.1.3 Phenotypic data

Parent studies have measured thousands of phenotypic and environmental risk factors. The phenotypic foci of the current studies include heart (38%), lung (33%), blood (8%), or sleep phenotypes (1%), while the remaining 20% are cohort studies with many HLBS phenotypes (Supplementary Figure 1 and Supplementary Table 2). There is considerable overlap in phenotypes among studies. For example, many lung-focused studies have heart phenotypes and vice versa. Many of the studies also have a wide range of other biomedical phenotypes including diabetes, kidney disease, and osteoporosis.

Common phenotypic measures across TOPMed Parent studies provide opportunities for cross-study analyses to gain power in detecting genetic effects. However, these studies differ in how their phenotypic data were collected, annotated, and structured. Creating harmonized phenotypic data sets for cross-study analyses is therefore a challenging and largely manual process.

Many TOPMed Working Groups harmonize phenotypes within their focus area. Generally, the writing group for a given paper develops an analysis plan that defines the primary outcome and any necessary covariates. Subsequently, selected Parent studies each contribute source data and modify or transform it to fit these definitions. The individual-level data are then pooled and evaluated for homogeneity across Parent studies.

The TOPMed Data Coordinating Center (DCC) also harmonizes phenotypes centrally, in collaboration with phenotype-domain experts in the Working Groups and data managers from the Parent studies. The DCC process uses study data from existing dbGaP accessions to enable provenance-tracking and reproducibility by the scientific community, and includes the following steps:

1. Precisely define the harmonized phenotype concept.
2. Identify dbGaP phenotype variables in each participating study that appear to fit the definition (with or without modification).
3. Determine which of these dbGaP variables are sufficiently equivalent to allow a meaningful harmonization[11].

4. Perform quality control of the selected dbGaP variables.
5. Define and implement algorithms to modify or transform the dbGaP variables as needed to produce a harmonized version.
6. Perform quality control on the harmonized variables across multiple studies to evaluate homogeneity, and refine previous steps as needed.
7. Document the process to ensure full reproducibility.
8. Submit harmonized phenotypes with documentation to dbGaP for distribution to the scientific community.

Depending on the availability of study phenotypes, their heterogeneity, and the results of data QC, this process is often iterative before a harmonized phenotype is finalized.

The DCC-harmonized data for an initial set of frequently-used phenotypes are being submitted to each study's TOPMed dbGaP accession (Supplementary Table 2) for release to the scientific community. These variables currently include subcohort, sex, race, ethnicity, recruitment site, height, weight, BMI, and smoking behavior (ever smoker and current smoker). Age at measurement is paired with each variable, except for basic demographics.

In addition, the TOPMed Lung Working Groups are utilizing an extensive lung phenotype harmonization project by the NHLBI Pooled Cohorts Study for lung function[12].

##### 1.1.4 Participant diversity

As of 2009, approximately 96% of participants in genome-wide association studies (GWAS) were of primarily European ancestry; by 2016, this figure was 80% mainly due to an increase in the study of East Asian populations, but the number of GWAS studies involving participants of African ancestry, Amerindian ancestry, and Hispanic or Latino ethnicity had not substantially increased[13]. Thus, in TOPMed, a concerted effort is being made to create a resource well-suited for investigating health issues affecting diverse populations. The populations sampled in Parent studies are mainly from the United States, although ~18% are from non-U.S. locations, including Brazil, Samoa, Costa Rica, Barbados, Taiwan, Pakistan, and multiple countries in Africa and Europe.

Race, ethnicity, and ancestry information was obtained from participant questionnaires and/or study inclusion criteria. This information was consolidated across studies into five broad categories. The entire current set of ~145,000 selected participants consists of approximately 40% European, 32% African, 16% Hispanic/Latino, 10% Asian and 2% 'Other' ancestries/ethnicities (Supplementary Figure 2. For the Freeze 5 genotype call set, sample numbers by study are provided for ancestry/ethnicity group and for sex in Supplementary Figures 21 and 22, respectively. In Freeze 5, the overall percentage of females is 60%.

Population structure and relatedness were evaluated in order to account for their effects in association studies, as well as to detect pedigree errors. We estimated principal components of genotypic data in the Freeze 5 call set using PC-AiR[14] and PC-Relate[15], which distinguish close relatedness from more distant relatedness due to population structure, while accounting for admixture. Supplementary Figure 23 (left panel) shows the expected separation among the major continental ancestral groups. Supplementary Figure 23 (right panel) shows differentiation among subgroups, including distinct differences between participants in the Old Order Amish study and other European ancestry groups (PC6), and between participants in the Costa Rican Asthma study and other Central Americans (see also Supplementary Figure 24). Kinship coefficient estimates show that a majority of Parent studies have substantial numbers of first and second degree relatives (Supplementary Figures 25 and 26). A small number of individuals were participants in more than one TOPMed study, resulting in cross-study duplicates and some cross-study relatives (see the Circos[16] plot in Supplementary Figure 26).

##### 1.1.5 Multi-omics assays

The TOPMed program is adding multi-omic assays to samples from participants with sequenced genomes. Initially, a multi-omics pilot study was performed on ~2,000 blood samples from ~1,000 participants in the Multi-Ethnic Study of Atherosclerosis (MESA), an on-going cohort study with extensive longitudinal phenotype data. The pilot measured DNA methylation, RNA-seq, metabolomics, and proteomics, using technologies described below. Subsequently, TOPMed initiated assays of DNA methylation on ~30,500 samples, RNA-seq on ~17,000 samples, metabolomics on ~7,800 samples and proteomics on ~950 samples. These samples derive from 15 different studies and multiple tissue types, including blood, nasal, lung, and cardiovascular tissues, as well as participant-specific, induced pluripotent stem cells. These TOPMed omics data are scheduled for release on dbGaP beginning in 2019. Several TOPMed Parent studies have omics data sets collected outside of the TOPMed program, some of which are currently available on dbGaP (e.g. the Framingham Heart Study; see "Molecular Data" under Parent study accession numbers in Supplementary Table 2).

TOPMed performed RNA-seq, proteomic, metabolomic, and methylation assays in a pilot project, using blood samples from each of ~1,000 participants in the Multi-Ethnic Study of Atherosclerosis (MESA). One goal of the project was to assess changes in omics values over time, in conjunction with changes in biomedical phenotypes. To this end, the pilot included peripheral blood mononuclear cells (PBMCs), plasma, and DNA samples drawn at each of two time points approximately 10 years apart, Exam 1 (2000-2002) and Exam 5 (2010-2012), from each participant. Each omics assay type was performed on largely the same set of paired samples from participants whose DNA had already been whole genome sequenced. Multiple quality control features were included in each assay to evaluate reproducibility and batch effects. The data from this pilot project are scheduled for release on dbGaP in 2019.

The following assay methods were used:

1. RNA-seq: PolyA+ mRNA libraries were prepared from PBMCs (Illumina TruSeq™) from the ~1,000 pairs of blood samples, and sequenced on Illumina HiSeq 4000 with target coverage of >40M reads at two genomic centers. In addition, RNA-seq was performed on cell-sorted CD19+ monocytes and on CD4+ T cells from ~400 participants obtained at Exam 5. The assays were performed and the resulting data processed at two different genomic centers using a harmonized pipeline[17].
2. Metabolomics: Three methods were used for assaying metabolites in plasma samples, each based on chromatography and mass spectroscopy. These methods included non-targeted lipids (228 known and 2,662 unknown), non-targeted polar metabolites (253 known and 3,966 unknown) and 84 targeted central metabolites.
3. Proteomics: DNA aptamer technology[18] (SOMAscan®[19]) was used to assay more than 1,300 proteins in plasma samples. In this technology, single-stranded DNAs composed of modified nucleotides are selected from libraries based on affinity to specific proteins. These reagents capture proteins in a complex mixture and their concentrations are estimated by DNA detection technology.
4. Methylation: Microarray technology was used to assay methylation at CpG sites in genomic DNA samples. The assays were performed using the Illumina EPIC array[20], which targets over 850,000 CpG sites in the genome. All DNA samples were treated with bisulfite to assay a combination of 5-methylcytosine (5-mC) and 5-hydroxymethylcytosine (5-hmC), and ~200 of these matched samples were also treated with an oxidizing reagent to distinguish between 5-mC and 5-hmC.

Another pilot study was carried out in 2017 using 950 plasma samples from Generation 3 participants in the Framingham Heart Study. These samples were used for proteomic profiling using the SOMAscan® aptamer technology described above.

Supplementary Table 13 provides information about the omics centers that performed assays for these pilot studies. The same centers are providing assays for omics assays in 2018 and 2019, using the same basic technologies described above.

#### 1.1.6 Resources available

Following quality control and dbGaP curation, TOPMed data are available in dbGaP accessions with accompanying study documents that describe the methods of phenotypic and omics data collection. WGS data include read alignments, genotype call sets, and related quality metrics. Methods of sequence data acquisition and processing are provided for each data Freeze. Metadata also include linking of DNA sample and participant identifiers; DNA sample attributes; participant consent group; and, for many studies, pedigree structures. A detailed guide to finding and using TOPMed data in dbGaP accessions is provided on the TOPMed website[21].

The TOPMed variant browser (BRAVO[22]) provides the chromosome location, alleles, TOPMed-wide allelic frequencies, and other characteristics of all single nucleotide or short indel variants called in the current data Freeze. A detail page for each variant shows the distributions of sequencing depth and genotype quality scores for carriers and non-carriers of the non-reference allele, as well as QC metrics that were used for site-level filtering. A variant site list deposited in NCBI dbSNP is available for download[23].

Phased genotypes from TOPMed data Freeze 5 will soon be made publicly available as an imputation reference panel in the Michigan Imputation Server[24]. This panel is a significant improvement upon existing panels such as 1000 Genomes[25] and the Haplotype Reference Consortium[26] because of the high quality of TOPMed WGS data, the ancestral/ethnic diversity in its samples, and the large sample size. In particular, these improvements allow more accurate imputation of low frequency variants, which will significantly leverage the value of sample sets with array-based genotypes.

The TOPMed Ethical, Legal, and Social Implications (ELSI) committee moderates discussion of matters regarding access to and use of TOPMed data. The committee has considered consent, participant privacy, and implementation issues regarding: (1) public sharing of variant summary data in the TOPMed variant server; (2) public access to use of a TOPMed reference panel in a genotype imputation server (without sharing individual-level data); and (3) return of results to study participants. Summary reports on the TOPMed website[27] contain points for study investigators to consider in making decisions about each issue.

#### 1.2 Whole Genome Sequencing Accessibility Mask

For the population genetic analysis it is important to exclude genomic regions with elevated false positive and false negative variants discovery rates due to pure accessibility by next-generation sequencing methods in these regions. We used per sample coverage information in order to create the accessible genome mask of GRCh38 for our analysis. First, for each of 1,000 randomly selected individuals from TOPMed freeze 5, we computed base-pair coverage using reads with mapping quality greater than 20 and base quality greater than 20. Second, for each base-pair we aggregated coverage across all individuals and computed summary statistics: average coverage, percent of individuals with coverage >1x, >5x, >10x, >50x, and >100x. These coverage summary statistics are available from BRAVO variant browser at bravo.sph.umich.edu. Base-pairs with N reference allele, low base quality, or where all reads had low map quality were not considered when aggregating. Finally, a base-pair was declared not accessible if: i) reference allele was N; ii) base quality was below threshold or all map qualities were below threshold across all individuals; iii) computed summary statistics didn't fall between 1 and 99 percentiles for autosomal chromosomes and pseudoautosomal regions on chromosome X, and between 1 and 99.9 percentiles for non-pseudoautosomal regions on chromosome X.

#### 1.3 Comparison to low coverage WGS and high coverage WES in Framingham Heart Study

Investigators in the Framingham Heart Study (FHS) evaluated WGS data from TOPMed in comparison with sequencing data from CHARGE Consortium WGS and Whole Exome Sequencing (WES). Supplementary Table 15 provides the counts and depth of each sequencing effort. The overlap of these three groups is 430 FHS study participants, on whom we report here. We use a subset of 263 unrelated study participants to calculate the numbers of singletons and doubletons, minor allele frequency (MAF), heterozygosity, and all rates, in order to avoid bias from the family structure. There have been other sequencing efforts in FHS, such as the Exome Sequencing Project, but there was insufficient overlap to include them here.

Supplementary Table 19 shows the number of non-monomorphic variants in each sequencing set. Note that the sets containing all sequenced subjects had more variants that the subgroup of 430 subjects. For the two WGS efforts, the overlap among the 430 subjects of common variants with MAF > 20% is nearly 3 million (Supplementary Figure 27). The CHARGE low coverage WGS has more variants with MAF > 20% than TOPMed; it is likely that different calling and quality control strategies explain this circumstance. Among the 263 study participants, the proportion of variants observed in TOPMed that are also seen in the CHARGE WGS varies from ~67% for rare MAF to ~85% for common variants. Hence, there is a great deal of overlap between the two sequencing efforts, even though CHARGE WGS is low coverage and TOPMed is deep WGS.

Using a set of 263 unrelated individuals, the number of SNV singletons and doubletons found in TOPMed is 9,624,993 and 1,901,162 respectively, while these numbers are lower in the CHARGE WGS at 5,379,585 and 1,344,916. These counts resulted in an average of 36,596.9 singletons and 7,228.7 doubletons per person in TOPMed and 20,454.7 singletons and 5,113.7 doubletons per person in the CHARGE WGS. Dividing the total number for each person by the number of non-monomorphic variants in a dataset (constant for all people in a dataset) yields rates of singletons and doubletons in TOPMed as $1.65 \times 10^{-3}$ and $3.27 \times 10^{-4}$ and in CHARGE WGS $1.16 \times 10^{-3}$ and $2.9 \times 10^{-4}$. Hence, the rates of singletons and doubletons are fairly similar between the two WGS efforts.

The rate of bi-allelic SNVs is similar in the two WGS efforts (Supplementary Table 20). The numbers in parentheses are the average number of SNVs seen per person, summing the heterozygous and homozygous calls for the minor allele of each SNV. The rates indicate the average proportion of SNVs that are heterozygous or homozygous for the minor allele. These rates suggest that about 14-15% of variants are heterozygous or homozygous for the minor allele among all SNVs, while the rate in exonic SNVs is lower.

Across all three sequencing efforts (TOPMed, CHARGE WGS and CHARGE WES), rare variants predominated (Supplementary Table 21). Among the 263 unrelated study participants, TOPMed had the greatest number of variants with MAF < 0.005, constituting 47% of the total number. The percentage of variants in this MAF range in the CHARGE WGS was 38% and in the CHARGE WES 60%. Other than the rarest variants, the two WGS efforts have similar numbers in each MAF frequency bin, although the CHARGE WGS has somewhat more, especially in the MAF > 0.05 range.

To focus on the exome, we used Ensembl gene GRCh37.p13 to define exonic regions in the WGS data. The number of exonic SNVs called was greatest in TOPMed with ~17-18% more variants (Supplementary Table 3). The numbers of multi-allelic variants and indels were far fewer relative to SNVs. Fewer multi-allelic variants were seen in the CHARGE WES than in TOPMed (9,226 vs 11,078). The numbers of indels seen in CHARGE WES were about half the ones seen in TOPMed (3,209 vs 5,794). We do not have counts for multi-allelic variants and indels for CHARGE WGS.

We compared the concordance of the calls in the sequencing efforts of the exome with those on the Exome Chip (Illumina Infinium HumanExome BeadChip array v1.0). We found very high concordance with most having concordance > 99%. Only the very rare variants had lower concordance, but still high at > 96.5%. The concordance of the TOPMed exonic variants was the highest across the MAF range with mean concordance rates of 0.999 across the minor allele spectrum; the light-coverage CHARGE WGS had the lowest among the three sequencing sets with concordance rates ranging from 0.990 to 0.998.

In the exome, the average heterozygosity per person for SNVs was 7.97% per variant for TOPMed, 9.66% for CHARGE WGS and 6.85% for CHARGE WES. The average number of heterozygous SNVs per person was 13,208 for TOPMed, 14,601 for CHARGE WGS and 9,640 for CHARGE WES, matching expectations.

Comparing the number of exonic SNVs across the MAF range, TOPMed had the greatest number of rare variants, followed by the CHARGE WES with the low coverage CHARGE WGS having the lowest number (Supplementary Figure 28). In contrast, the light-coverage CHARGE WGS had a greater number of common variants in the 1-50% MAF range. These comparisons match what was observed for the total number of variants (Supplementary Table 21). The CHARGE WES consistently had higher numbers of multi-allelic variants across the minor allele spectrum than TOPMed, nearly double for more common variants of this type. In contrast, TOPMed had similar, but consistently more indels than the CHARGE WES, especially among indels with frequencies greater than 1%.

More than half of exonic variants were non-synonymous in the three sequencing efforts (Supplementary Table 3). In TOPMed 0.96% of variants were LOF while in the light-coverage CHARGE WGS this percentage was 0.87% and in CHARGE WES it was the same as TOPMed. The numbers of LOF, missense and non-synonymous SNVs by minor allele frequency were comparable across the three sequencing datasets with somewhat fewer among SNVs in the CHARGE WES with MAF > 1%.

In summary, the results suggest more common variants were seen in the light-coverage CHARGE WGS in comparison to TOPMed, while the deep-sequencing in TOPMed revealed many more rare variants compared to TOPMed. Regardless, the vast majority of variants seen in TOPMed were also seen in CHARGE WGS. While CHARGE WGS had somewhat higher rates of common SNVs, this circumstance could possibly be due to different calling and quality control procedures. Other characteristics were similar for the three sequencing datasets. In particular, the heterozygosity rates were similar and the numbers of variants by functional category were similar. Most important, the concordance of each sequencing set with genotypes on the Exome Chip were very high. Hence, we feel confident in using any of the three sequencing sets in data analyses.

#### 1.4 Novel genetic variants in unmapped reads

##### 1.4.1 Assembly of hominid non-human-reference sequences from unmapped reads

For each TOPMed data freeze 3 sample BAM (mapped against GRCh37), read-pairs with at least one unmapped mate (SAM flags 4 and/or 8) were extracted and converted into FASTQ format, excluding any read shorter than 30bp. Reads were then sorted by read name and any singleton filtered out. Next, duplicate read pairs identified based on two sets of alternating 12bp signatures in the beginning of each mate (from position 10) were removed. The remaining read-pairs were screened for adapter sequences and low quality bases (Q<20), that were trimmed using Cutadapt 1.8.1[28]. Read-pairs that, after this process, had either of the two mates shorter than 50bp were removed and the remaining pairs mapped against a phiX reference using GEM mapper[29] for spike-in filtering. Finally, samples with more than 500,000 processed fully unmapped read-pairs (1 million reads) at this stage were considered as highly contaminated and were discarded. On a per-sample basis, reads were then assembled into contigs using ABySS v.2.0.2[30], exploring different *K*-mer sizes (37, 47, 57, 67, 77, 87 and 97). All resulting contigs longer than 200bp from the seven assemblies were then pooled together. Contigs that fully aligned into other contigs using MegaBLAST[31] (coverage = 100%, termini slop <= 5bp, identity >= 99%) were removed and contigs that extended from other contigs (MegaBLAST, terminal overlap length >= 150bp, termini slop <= 5bp, identity >= 99%) were merged. This process was done iteratively, prioritizing on alignment length, until no actionable alignments were found. All assembled contigs were then queried against 5 hominid reference genomes (*Pan paniscus* – panPan1; *Pan troglodytes* – panTro5; *Gorilla gorilla* – gorGor5; *Pongo abelii* – ponAbe2; and *Homo sapiens* – hg38) downloaded from the UCSC Genome Browser[32], using MegaBLAST. Contigs for which no High-scoring Segment Pair (HSP) covering more than 80% of the sequence with more than 80% identity and no HSP longer than 1000bp was found, were discarded. Finally, only contigs with the best hit to any of the four non-human hominid references were kept.

##### 1.4.2 Reference-based contig merging and scaffolding

Filtered contigs from all samples were pooled together and aligned, using BWA-MEM[33], to an index consisting of the afore-mentioned non-human hominid genome references. Contigs aligning to the same genomic region are likely to represent the same ancestral sequence and were, therefore, merged together based on the alignment pileup. Given that four references were concurrently used, up to four contig clusters could be formed for each ancestral sequence. To address this, cluster contigs were further collapsed and extended using the method described above for merging multi K-mer assemblies. To address potential mis-assemblies resulting from this process, merged contigs were remapped and then split at alignment breakpoints. Low complexity contigs, identified using PRINSEQ lite (trinucleotide entropy < 70)[34], were removed and contigs that, at this stage, best-mapped to human when re-queried against the five hominid genomes or that virtually fully aligned to the human reference (reference alignment span >= 95% contig length; or reference alignment span >= contig length - 20 bp) were also discarded. The remaining contigs were joined into scaffolds based on alignment proximity (maximum distance <= 7.5 kb) in the hominid genomes, with gap sizes corresponding to the alignment distance. In situations, where alignments partially overlapped, the overlapping region of the shorter contig was eroded and a single-nucleotide gap was placed between the two contigs.

##### 1.4.3 Reference placement and breakpoint calling

To identify candidate regions for the location of each scaffold/contig in the human reference GRCh38, we took the terminal alignment positions of each contig against the four hominid genomes and transposed them into the human reference using liftOver[35]. When the alignment coordinates of a given contig were located within an alignment gap to the human reference – leading to a null human coordinate result, positions upstream of the alignment start and downstream of the alignment end were iteratively tried (in increments of 100bp, up to 60kb), until the coordinate was converted. A candidate region was defined when both the start and end of each contig are successfully converted into the same chromosome, and in the correct orientation. For scaffold placement, the outmost alignment coordinates of the terminal contigs were considered.

Given the intervals defined by liftOver, we next wanted to resolve the exact breakpoint position(s) for each contig. We reasoned that contigs would fall into one of the following three categories: a) insertion contigs extending into the reference sequence on both ends, b) “hanging” insertion (or *breakend*) contigs extending into the reference sequence on only one end, or c) other unanchorable contigs (not extending into reference sequence, or potential noise). We used AGE[36] to align contigs allowing for one long insertion, testing first for insertion contigs anchored on both ends. AGE uses a dynamic programming approach to finding the optimal breakpoint position between two partial alignments. If the two partial alignments identified by AGE fulfilled minimal criteria (at least 94% identity, at least 15bp aligned on each side, and within 15bp of contig ends), an insertion event was called. If no such alignment was identified, we checked whether, within the given liftOver interval, BWA-MEM found a soft-clipped alignment (within 30bp of one contig end) with more than 30 clipped bases. If so, we called a breakend.

All contigs without liftOver interval and contigs not yielding a valid insertion or breakend call in the previous local step were aligned to GRCh38 using Gustaf[37] (minimum match length 100bp, maximum error rate 0.04). Gustaf chains local alignments allowing (theoretically) for any type of structural rearrangement and making no assumption about what type of rearrangements or how many breakpoints might be present. All contigs with an insertion or breakend call were extracted and, to achieve higher confidence and consistency with the above local approach, were realigned with AGE using the genomic interval identified by Gustaf. If no insertion call was possible with AGE, we again checked for partial BWA alignments and potentially added a breakend call.
The described placement steps were applied to both contigs and scaffolds. We used the scaffold results whenever the resulting breakpoint calls were either the same as for the individual contigs or when two breakend calls were joined into a single insertion call. For unplaced scaffolds or scaffolds where a single breakend was called, while the individual contigs lead to multiple, sufficiently different (>100bp apart) breakpoints, we discarded the scaffold results and kept the individual contig placements.

##### 1.4.4 Presence/absence calling

In order to call presence or absence of each identified variant within the set of 17,443 individuals, we aligned the originally extracted read pairs onto an index of GRCh38 plus our breakpoint-resolved insertions/breakends (extending putative insertion sequence 150bp into the reference – i.e. on both sides for insertion contigs, one side for breakend contigs). For a variant to be present in an individual, at least 50% of the putative inserted sequence needed to be covered with read depth >= 3, or at least 2 breakpoint crossing reads (mapQ>10, breakpoint overlap >= 10bp) needed to be observed on either breakpoint.

#### 1.5 Site Frequency Spectrum

One way of gaining inference into a population's demographic history is to study the site frequency spectrum (SFS), which shows the proportion of all variants found at each minor allele count.

Using a subsetted sample of 1,370 unrelated individuals each from East Asian, Europeans, and African ancestry we calculated the allele count of each site and generated a log-log histogram of their frequency. 1,370 constitutes all the East Asian samples, and we have subset the other populations randomly to control for the rarity of the variants. A similar procedure was done for the analysis that includes the Amish, except with 225 individual per cohort, which represents the total number of unrelated Amish individuals in the dataset. Using AC and AN values for each subsampled cohort, we used all variants passing typical variant QC (filter==“PASS”) to calculate the frequency of each allele count, as well as its proportion of the total number of variants in the cohort.

Comparing the SFS for 1,370 unrelated individuals from European, African, and East Asian ancestry individuals, we find that all three major groups show an excess of rare variation indicative of recent population expansions[38–40] and purifying selection, with European and East Asian ancestry populations exhibiting the greatest excess of rare variants (Supplementary Figure 29). In contrast, the Amish founder population has experienced a very severe and recent bottleneck, and their SFS exhibits a shift toward common variants (Supplementary Figure 30). This then gives a basis for the observation of increased heterozygosity in the Amish compared to East Asians and a higher singleton count in the East Asian groups (Figure 4)

#### 1.6 Admixture

Average ancestry proportions were calculated using a stratified random sample from participating TOPMed studies consented for population genetics research and a standard ADMIXTURE analysis[41,42]. 100 unrelated subjects were randomly sampled from each self-reported ancestry group within each TOPMed study. If the ancestry group contained fewer than 100 subjects, the entire unrelated group was included. In total, 4,444 subjects were included in the analysis. Sequencing data was filtered, removing indels, using a minimum MAF of 0.05 and a maximum $r^2$ value of 0.2. Ancestry proportions were calculated using ADMIXTURE v1.3.0. The value for K was chosen by a cross-validation procedure. The cross-validation errors of Ks 1 through 15 showed two minima, one at a K of 9 and another at a K of 13. K of 9 was chosen as the more conservative value. 20 replications were done; the replication with the highest log-likelihood was used.

ADMIXTURE analysis[41] revealed nine clusters in the dataset, three of which resemble different European ancestries and two of which represent African ancestries. Amish, East Asian, Native American, and Samoan ancestries are all represented by their “own” cluster. The first European cluster contributes similarly to European American and Hispanic/Latino cohorts, the second European cluster is at highest frequency in the European American cohorts, and a third European cluster is at highest frequency in the Hispanic/Latino cohorts. The European ancestry in the African American cohorts is predominantly represented by the second European cluster. We find shared ancestry between the Samoan and the MESA and WHI Asian American subsets, likely due to the recent shared ancestry of Samoan and East Asian populations[43,44].

#### 1.7 Demographic estimation under selection at linked sites

##### 1.7.1 Sample selection

In order to sample individuals with a high percentage of European ancestry and to prevent confounds introduced by population structure, we used two separate ascertainment schemes for selecting individuals in our study. First, the program RFMix[45] was run on 18,436 samples from the TOPMed data freeze 3 using the following parameter settings: PopPhased --num-threads 1 --min-node-size 5. For the reference panel, 938 samples from the Human Genome Diversity Panel (HGDP) were used. The 53 populations of HGDP were condensed into 7 super-populations: 1) Sub-Saharan African (n=104), 2) Central and South Asian (n=200), 3) East Asian (n=229), 4) European (n=154), 5) Native American (n=63), 6) Oceanian (n=28), and 7) Middle Eastern (n=160). After running RFMix, we summed local ancestries assigned for each TOPMed sample to create a vector of global ancestries corresponding to the 7 HGDP super-populations. We then selected individuals that had greater than or equal to 90% global European ancestry (Supplementary Figure 31). To further limit potential population structure, we also filtered individuals for those that belonged to the population 1 ('European A') cluster from the PCA/k-means clustering methodology conducted for the iHS/EHH analyses (Supplementary Information 1.8). Finally, we limited our selected individuals to those that were unrelated and gave consent for performing population genetics research. This resulted in a total of 2,416 samples. When measuring the site-frequency spectrum across these samples as a function of sample size, we sampled progressively larger random samples of 50 individuals (100 chromosomes) each, generating 49 discrete sample sizes (2N=100, 200, 300…4800, 4832).

##### 1.7.2 Site filtering/ascertainment

In order to perform inference using a high-quality set of neutral sites that are least influenced by the direct effects of natural selection and putative selective sweeps and to avoid potential sequence/mapping error, we performed several steps to filter the genome. Many of these filtering steps were based off of the ascertainment scheme used by Torres et al. 2018[46]. Specifically, the following filters were applied (all filters are in hg19 and only autosomes were kept for analyses):

1. Coding regions: coding exons annotated in the UCSC known genes track (table: knownGene, track: UCSC Genes) were removed.

2. phyloP: Sites with phyloP[47] scores > 1.2 or < -1.2 were removed to limit the effects of natural selection due to conservation or accelerated evolution. Scores were downloaded from http://hgdownload.cse.ucsc.edu/goldenPath/hg19/phyloP46way/.
3. phastCons: Regions in the UCSC conservation 46-way track (table: phastCons46wayPlacental)[48] were removed to limit the effects of natural selection due to conservation.
4. CpG: CpG islands in the UCSC CpG islands track were removed because of their potential role in gene regulation and/or being conserved.
5. ENCODE blacklist: Regions with high signal artifacts from next-generation sequencing experiments discovered during the ENCODE project[49] were removed.
6. Simple repeats: Regions in the UCSC simple repeats track were removed due to potential misalignments with outgroups and/or being under natural selection.
7. Gaps/centromeres/telomeres: Regions in the UCSC gap track were removed, including centromeres and telomeres.
8. Segmental duplications: Regions in the UCSC segmental dups track[50] were removed to limit potential effects of natural selection.
9. Transposons: Active transposons (HERVK retrotransposons, the AluY subfamily of Alu elements, SVA elements, and L1Ta/L1pre-Ta LINEs) in the human genome were removed.
10. Recent positive selection: Regions inferred to be under hard and soft selective sweeps (using iHS and iHH12 regions from selscan v1.2.0[46,51]); within Thousand Genomes phase 3[25] European and African populations were removed.
11. Non-coding transcripts: Non-coding transcripts from the UCSC genes track were removed to limit potential effects of natural selection.
12. GC-biased gene conversion (gBGC): Regions in UCSC phastBias track[52] from UCSC genome browser were removed to limit regions inferred to be under strong GC-biased gene conversion.
13. Recombination hotspots: All sites within 1.5 kb (i.e., 3 kb windows) of sites with recombination rates >= 10 cM/Mb in the 1000G OMNI genetic maps for non-admixed populations (downloaded from ftp://ftp.1000genomes.ebi.ac.uk/vol1/ftp/technical/working/20130507_omni_recombination_rates/) and the HapMap II genetic map[53] were removed. 1.5 kb flanking regions surrounding the center of hotspots identified by Pratto et al. 2014[54] (downloaded from http://science.sciencemag.org/content/sci/suppl/2014/11/12/346.6211.1256442.DC1/1256442_DatafileS1.txt) were also removed, except for the cases in which the entire hotspot site was greater than 3 kb in length (in which case just the hotspot was removed).

Positions in the genome were then annotated for how strongly affected they are by selection at linked sites by using the background selection (BGS) coefficient, $B$[55] (McVicker's $B$ statistic; downloaded from http://www.phrap.org/othersoftware.html). BGS is a process by which neutral variation in the genome is affected by purifying selection via genetic linkage to deleterious sites[56–58]. $B$ represents the fraction of neutral genetic variation at a particular site in a population suffering BGS relative to a neutrally evolving population and varies between 0 and 1, with BGS increasing in strength as values of $B$ approach 0. Positions for $B$ were lifted over from hg18 to hg19 using the UCSC liftOver tool. Sites that failed to uniquely map from hg18 to hg19 or failed to uniquely map in the reciprocal direction were excluded. Sites lacking a $B$ value were also ignored. We used all sites annotated with a $B$ value for performing general analyses. However, when performing demographic inference, we only focused our analyses on those regions of the genome within the top 1% of the genome-wide distribution of $B$ ($B >= 0.994$). These sites correspond to regions of the genome inferred to be under the weakest amount of BGS (i.e., under the weakest effects of selection at linked sites).

Sites in the genome were also polarized to ancestral and derived states using ancestral annotations called with high-confidence from the GRCh37 e71 ancestral sequence (downloaded from: ftp://ftp.ensembl.org/pub/release-71/fasta/ancestral_alleles/homo_sapiens_ancestor_GRCh37_e71.tar.bz2) from Ensembl[55,59], which used a multiple species alignment of 6 primates to infer the ancestral state using the Enredo-Pecan-Ortheus (EPO) pipeline[60,61]. All of the filtering steps described, including the annotation for $B$ and polarization for ancestral/derived state, left 1,377,691,456 sites within the genome for use in our study, including 10,977,437 sites with $B >= 0.994$. Finally, we filtered polymorphic sites within the filtered genome on being di-allelic only. This left a total 20,324,704 polymorphic sites across the 2,416 European samples, including 191,631 polymorphic sites that had $B >= 0.994$. To generate a set of fourfold degenerate synonymous sites, all coding sites within the genome were annotated using the program ANNOVAR[62] using Gencode V19 annotations. This resulted in 5,188,972 total sites. 4,718,653 sites were left after filtering for high-confidence ancestral/derived states, of which 91,177 were polymorphic (di-allelic) across the 2,416 European samples.

##### 1.7.3 Demographic inference

We performed demographic inference using the program *moments*[63], which fits a specified demographic model to an observed site-frequency spectrum. For our study, we specified a model of exponential growth with three total parameters ($N_{Eur0}$, $N_{Eur}$, $T_{Eur}$). This included two free parameters: the starting time of exponential growth ($T_{Eur}$) and the ending population size after growth ($N_{Eur}$). The ancestral size parameter (i.e, the population size when growth begins), $N_{Eur0}$, was kept constant in our model such that the relative starting size of the population was always 1. We applied the inference procedure to the 2,416 European samples using either fourfold degenerate sites or sites where *B* >= 0.994 (highest 1% *B* bin). The site-frequency spectrum used for inference was unfolded and based on the polarization step described above. The inference procedure was fit using sample sizes (2N) of 1000, 2000, 3000, 4000, and 4832 samples (i.e., chromosomes). The inference procedure was run from different initial starting points hundreds of times for each sample size and dataset to ensure convergence on a global optimum. Attempts at using samples sizes smaller than 2N=1000 for inference resulted in convergence issues, likely because of poor model fit.

To convert the scaled genetic parameters output by the inference procedure moments to physical units, we used the resulting theta (also inferred by *moments*) and a mutation rate of $1.66 \times 10^{-8}$ [64] to generate corresponding effective population sizes ($N_e$). In order to account for the fact that fourfold degenerate sites and sites from regions within the highest 1% *B* bin are ascertained from different effective sequence lengths, we had to first normalize theta by their corresponding lengths. These lengths were 4,718,653 sites and 10,977,437 sites for fourfold degenerate sites and highest 1% *B* sites, respectively. To convert time to years, we used a generation time of 25 years. 95% confidence intervals were generated by resampling the SFS 1,000 times and using the Godambe Information Matrix to generate parameter uncertainties[65].

#### 1.8 Selection

From the previously performed principal components analyses (PCA) on a set of 18,234 individuals from the TOPMed data freeze 3. We used k-means clustering to cluster individuals on the first 7 PCs. Clustering was initially run on various values of k from 1 to 20, with 25 restarts each. We plotted the total within class sum of squares from this initial clustering round and chose k=9 since large decreases in the sum of squares diminish after this point (Supplementary Figure 32). We then re-ran the clustering for k=9 with 250 restarts to finalize the cluster assignments for the 18,234 samples. This left us with population clusters that roughly aligned with several self-report categories, and we assigned population labels based on these (see Supplementary Table 16).

We identified individuals from these populations for which we had clear consent and were left with six populations with a non-zero numbers of individuals (Supplementary Table 16). Once we had our set of individuals identified for analysis, we partitioned each population and analyzed each population separately.

We used only biallelic sites, and the WGSA pipeline to infer ancestral state. Any sites for which ancestral state was ambiguous were filtered. Alleles were then polarized such that 1 represents the derived allele and 0 represents the ancestral allele.

We used the program selscan[51] to perform all iHS scans[66] in each population using default parameters. This includes a filtering step where all sites with minor allele frequency less than 0.05 are filtered before analysis and sites near chromosome boundaries being excluded due to a truncation of the EHH integration curve. The final number of sites for which we calculated iHS scores, per population, is in Supplemental Table 17.

We then normalized raw iHS scores within 20 frequency bins, a requirement since raw iHS is correlated with allele frequency. After normalization, iHS scores are approximately normally distributed under neutrality, and furthermore true selected regions tend to have clusters of extreme scores[66]. We therefore follow Voight et al. 2006 and partition the genome into 100kb windows and count the proportion of sites with $|iHS| > 2$. We then partition windows into 20 bins based on total number of sites within that window, and select the highest 1% of windows with the largest proportion of extreme iHS scores. These represent our putatively selected regions. A full list of genes found in significant regions, and the populations in which they were found significant, is given in Supplementary File 2. A full list of regions identified as significant in each population can be found in Supplementary Files 3-8.

### 2 Study Acknowledgments

**NHLBI TOPMed: Genetics of Cardiometabolic Health in the Amish**

The Amish studies upon which these data are based were supported by NIH grants R01 AG18728, U01 HL072515, R01 HL088119, R01 HL121007, and P30 DK072488. See publication: PMID: 18440328

**NHLBI TOPMed: Trans-Omics for Precision Medicine Whole Genome Sequencing Project: ARIC**

The Atherosclerosis Risk in Communities study has been funded in whole or in part with Federal funds from the National Heart, Lung, and Blood Institute, National Institutes of Health, Department of Health and Human Services (contract numbers HHSN268201700001I, HHSN268201700002I, HHSN268201700003I, HHSN268201700004I and HHSN268201700005I). The authors thank the staff and participants of the ARIC study for their important contributions.

**NHLBI TOPMed: The Genetics and Epidemiology of Asthma in Barbados**

The Genetics and Epidemiology of Asthma in Barbados is supported by National Institutes of Health (NIH) National Heart, Lung, Blood Institute TOPMed (R01 HL104608-S1) and: R01 AI20059, K23 HL076322, and RC2 HL101651. For the specific cohort descriptions and descriptions regarding the collection of phenotype data can be found at: https://www.nhlbiwgs.org/group/bags-asthma. The authors wish to give special recognition to the individual study participants who provided biological samples and or data, without their support in research none of this would be possible.

**NHLBI TOPMed: Cleveland Clinic Atrial Fibrillation Study**

The research reported in this article was supported by grants from the National Institutes of Health (NIH) National Heart, Lung, and Blood Institute grants R01 HL090620 and R01 HL111314, the NIH National Center for Research Resources for Case Western

Reserve University and the Cleveland Clinic Clinical and Translational Science Award (CTSA) UL1-RR024989, the Department of Cardiovascular Medicine philanthropic research fund, Heart and Vascular Institute, Cleveland Clinic, the Fondation Leducq grant 07-CVD 03, and The Atrial Fibrillation Innovation Center, State of Ohio.

**NHLBI TOPMed: The Cleveland Family Study (WGS)**

Support for the Cleveland Family Study was provided by NHLBI grant numbers R01 HL46380, R01 HL113338 and R35 HL135818.

**NHLBI TOPMed: Cardiovascular Health Study**

This research was supported by contracts HHSN268201200036C, HHSN268200800007C, HHSN268201800001C, N01-HC85079, N01-HC-85080, N01-HC-85081, N01-HC-85082, N01-HC-85083, N01-HC-85084, N01-HC-85085, N01-HC-85086, N01-HC-35129, N01-HC-15103, N01-HC-55222, N01-HC-75150, N01-HC-45133, and N01-HC-85239; grant numbers U01 HL080295, U01 HL130114 and R01 HL059367 from the National Heart, Lung, and Blood Institute, and R01 AG023629 from the National Institute on Aging, with additional contributions from the National Institute of Neurological Disorders and Stroke. A full list of principal CHS investigators and institutions can be found at https://chs-nhlbi.org/pi. Its content is solely the responsibility of the authors and does not necessarily represent the official views of the National Institutes of Health.

**NHLBI TOPMed: Genetic Epidemiology of COPD (COPDGene) in the TOPMed Program**

This research used data generated by the COPDGene study, which was supported by NIH Award Number U01 HL089897 and Award Number U01 HL089856 from the National Heart, Lung, and Blood Institute. The content is solely the responsibility of the authors and does not necessarily represent the official views of the National Heart, Lung, and Blood Institute or the National Institutes of Health.

The COPDGene project is also supported by the COPD Foundation through contributions made to an Industry Advisory Board comprised of AstraZeneca, Boehringer Ingelheim, GlaxoSmithKline, Novartis, Pfizer, Siemens and Sunovion.

**NHLBI TOPMed: The Genetic Epidemiology of Asthma in Costa Rica**

This study was supported by NHLBI grants R37 HL066289 and P01 HL132825. We wish to acknowledge the investigators at the Channing Division of Network Medicine at Brigham and Women's Hospital, the investigators at the Hospital Nacional de Niños in San José, Costa Rica and the study subjects and their extended family members who contributed samples and genotypes to the study, and the NIH/NHLBI for its support in making this project possible.

**NHLBI TOPMed: Diabetes Heart Study African American Coronary Artery Calcification (AA CAC)**

This work was supported by R01 HL92301, R01 HL67348, R01 NS058700, R01 AR48797, R01 DK071891, the General Clinical Research Center of the Wake Forest University School of Medicine (M01 RR07122, F32 HL085989), the American Diabetes Association, and a pilot grant from the Claude Pepper Older Americans Independence Center of Wake Forest University Health Sciences (P60 AG10484).

**NHLBI TOPMed: Boston Early-Onset COPD Study in the TOPMed Program**

The Boston Early-Onset COPD Study (dbGaP accession number phs000946) was supported by the following NIH grants: R01 HL075478, U01 HL089856, and R01 HL113264.

**NHLBI TOPMed: Whole Genome Sequencing and Related Phenotypes in the Framingham Heart Study**

The Framingham Heart Study (FHS) acknowledges the support of contracts NO1-HC-25195 and HHSN268201500001I from the National Heart, Lung, and Blood Institute and grant supplement R01 HL092577-06S1 for this research. We also acknowledge the dedication of the FHS study participants without whom this research would not be possible. Dr. Vasan is supported in part by the

Evans Medical Foundation and the Jay and Louis Coffman Endowment from the Department of Medicine, Boston University School of Medicine.

**NHLBI TOPMed: Genes-environments and Admixture in Latino Asthmatics (GALA II) Study**

Supported by NIH and NHLBI grant # R01HL117004; study enrollment supported by NIEHS grant # R01ES015794, the Sandler Family Foundation, the American Asthma Foundation, the RWJF Amos Medical Faculty Development Program, Harry Wm. and Diana V. Hind Distinguished Professor in Pharmaceutical Sciences II.

All study collaborators: Shannon Thyne, UCSF; Harold J. Farber, Texas Children's Hospital; Denise Serebrisky, Jacobi Medical Center; Rajesh Kumar, Lurie Children's Hospital of Chicago; Emerita Brigino-Buenaventura, Kaiser Permanente; Michael A. LeNoir, Bay Area Pediatrics; Kelley Meade, Children's Hospital, Oakland; William Rodriguez-Cintron, VA Hospital, Puerto Rico; Pedro C. Avila, Northwestern University, Jose R. Rodriguez-Santana, Centro de Neumologia Pediatrica.

The authors acknowledge the families and patients for their participation and thank the numerous health care providers and community clinics for their support and participation in GALA II. In particular, the authors thank study coordinator Sandra Salazar; the recruiters who obtained the data: Duanny Alva, MD, Gaby Ayala-Rodriguez, Lisa Caine, Elizabeth Castellanos, Jaime Colon, Denise DeJesus, Blanca Lopez, Brenda Lopez, MD, Louis Martos, Vivian Medina, Juana Olivo, Mario Peralta, Esther Pomares, MD, Jihan Quraishi, Johanna Rodriguez, Shahdad Saeedi, Dean Soto, Ana Taveras.

See publication: PMID: 23750510

**NHLBI TOPMed: Genetic Epidemiology Network of Arteriopathy (GENOA)**

Support for GENOA was provided by the National Heart, Lung, and Blood Institute (HL054457, HL054464, HL054481, and HL087660) of the National Institutes of Health.

**NHLBI TOPMed: Genetic Epidemiology Network of Salt Sensitivity (GenSalt)**

The Genetic Epidemiology Network of Salt-Sensitivity (GenSalt) was supported by research grants (U01HL072507, R01HL087263, and R01HL090682) from the National Heart, Lung, and Blood Institute, National Institutes of Health, Bethesda, MD.

**NHLBI TOPMed: Genetics of Lipid Lowering Drugs and Diet Network (GOLDN)**

GOLDN biospecimens, baseline phenotype data, and intervention phenotype data were collected with funding from the National Heart, Lung and Blood Institute (NHLBI) grant U01 HL072524. Whole-genome sequencing in GOLDN was funded by NHLBI grant R01 HL104135 and supplement R01 HL104135-04S1.

**NHLBI TOPMed: Heart and Vascular Health Study (HVH)**

The research reported in this article was supported by grants HL068986, HL085251, HL095080, and HL073410 from the National Heart, Lung, and Blood Institute.

**NHLBI TOPMed: Hypertension Genetic Epidemiology Network (HyperGEN)**

The HyperGEN Study is part of the National Heart, Lung, and Blood Institute (NHLBI) Family Blood Pressure Program; collection of the data represented here was supported by grants U01 HL054472 (MN Lab), U01 HL054473 (DCC), U01 HL054495 (AL FC), and U01 HL054509 (NC FC). The HyperGEN: Genetics of Left Ventricular Hypertrophy Study was supported by NHLBI grant R01 HL055673 with whole-genome sequencing made possible by supplement -18S1.

**NHLBI TOPMed: The Jackson Heart Study**

The Jackson Heart Study (JHS) is supported and conducted in collaboration with Jackson State University (HHSN268201800013I), Tougaloo College (HHSN268201800014I), the Mississippi State Department of Health (HHSN268201800015I/HHSN26800001) and the University of Mississippi Medical Center (HHSN268201800010I, HHSN268201800011I and HHSN268201800012I) contracts from the National Heart, Lung, and Blood Institute (NHLBI) and the National Institute for Minority Health and Health Disparities (NIMHD). The authors also wish to thank the staff and participants of the JHS.

**NHLBI TOPMed: Multi-Ethnic Study of Atherosclerosis**

Whole genome sequencing (WGS) for the Trans-Omics in Precision Medicine (TOPMed) program was supported by the National Heart, Lung and Blood Institute (NHLBI). WGS for “NHLBI TOPMed: Multi-Ethnic Study of Atherosclerosis (MESA)” (phs001416.v1.p1) was performed at the Broad Institute of MIT and Harvard (3U54HG003067-13S1). Centralized read mapping and genotype calling, along with variant quality metrics and filtering were provided by the TOPMed Informatics Research Center (3R01HL-117626-02S1). Phenotype harmonization, data management, sample-identity QC, and general study coordination, were provided by the TOPMed Data Coordinating Center (3R01HL-120393-02S1). MESA and the MESA SHARe project are conducted and supported by the National Heart, Lung, and Blood Institute (NHLBI) in collaboration with MESA investigators. Support for MESA is provided by contracts HHSN268201500003I, N01-HC-95159, N01-HC-95160, N01-HC-95161, N01-HC-95162, N01-HC-95163, N01-HC-95164, N01-HC-95165, N01-HC-95166, N01-HC-95167, N01-HC-95168, N01-HC-95169, UL1-TR-000040, UL1-TR-001079, UL1-TR-001420. MESA Family is conducted and supported by the National Heart, Lung, and Blood Institute (NHLBI) in collaboration with MESA investigators. Support is provided by grants and contracts R01HL071051, R01HL071205, R01HL071250, R01HL071251, R01HL071258, R01HL071259, and by the National Center for Research Resources, Grant UL1RR033176. The provision of genotyping data was supported in part by the National Center for Advancing Translational Sciences, CTSI grant UL1TR001881, and the National Institute of Diabetes and Digestive and Kidney Disease Diabetes Research Center (DRC) grant DK063491 to the Southern California Diabetes Endocrinology Research Center.

**NHLBI TOPMed: Whole Genome Sequencing of Venous Thromboembolism (WGS of VTE)**

Funded in part by grants from the National Institutes of Health, National Heart, Lung, and Blood Institute (HL66216 and HL83141) and the National Human Genome Research Institute (HG04735).

**NHLBI TOPMed: MGH Atrial Fibrillation Study**

The research reported in this article was supported by NIH grants K23HL071632, K23HL114724, R21DA027021, R01HL092577, R01HL092577S1, R01HL104156, K24HL105780, U01HL65962, R01HL128914, and American Heart Association, 18SFRN34110082. The research has also been supported by an Established Investigator Award from the American Heart Association (13EIA14220013) and by support from the Fondation Leducq (14CVD01).

**NHLBI TOPMed: Partners HealthCare Biobank**

We thank the Broad Institute for generating high-quality sequence data supported by the NHLBI grant 3R01HL092577-06S1 to Dr. Patrick Ellinor. The datasets used in this manuscript were obtained from dbGaP at http://www.ncbi.nlm.nih.gov/gap through dbGaP accession number phs001024.

**NHLBI TOPMed: San Antonio Family Heart Study (WGS)**

Collection of the San Antonio Family Study data was supported in part by National Institutes of Health (NIH) grants R01 HL045522, MH078143, MH078111 and MH083824; and whole genome sequencing of SAFS subjects was supported by U01 DK085524 and R01 HL113323. We are very grateful to the participants of the San Antonio Family Study for their continued involvement in our research programs.

**NHLBI TOPMed: Study of African Americans, Asthma, Genes and Environment (SAGE) Study**

Supported by NIH and NHLBI grant # R01HL117004;study enrollment supported by the Sandler Family Foundation, the American Asthma Foundation, the RWJF Amos Medical Faculty Development Program, Harry Wm. and Diana V. Hind Distinguished Professor in Pharmaceutical Sciences II.

**NHLBI TOPMed: Genome-wide Association Study of Adiposity in Samoans**

Financial support from the U.S. National Institutes of Health Grant R01-HL093093. We acknowledge the assistance of the Samoa Ministry of Health and the Samoa Bureau of Statistics for their guidance and support in the conduct of this study. We thank the local village officials for their help and the participants for their generosity. The following publication describes the origin of the dataset: Hawley NL, Minster RL, Weeks DE, Viali S, Reupena MS, Sun G, Cheng H, Deka R, McGarvey ST. Prevalence of Adiposity and Associated Cardiometabolic Risk Factors in the Samoan Genome-Wide Association Study. Am J Human Biol 2014. 26: 491-501. DOI: 10.1002/jhb.22553. PMID: 24799123.

**NHLBI TOPMed: The Vanderbilt AF Ablation Registry**

The research reported in this article was supported by grants from the American Heart Association to Dr. Shoemaker (11CRP742009), Dr. Darbar (EIA 0940116N), and grants from the National Institutes of Health (NIH) to Dr. Darbar (R01 HL092217), and Dr. Roden (U19 HL65962, and UL1 RR024975). The project was also supported by a CTSA award (UL1 TR00045) from the National Center for Advancing Translational Sciences. Its contents are solely the responsibility of the authors and do not necessarily represent the official views of the National Center for Advancing Translational Sciences or the NIH.

**NHLBI TOPMed: The Vanderbilt Atrial Fibrillation Registry**

The research reported in this article was supported by grants from the American Heart Association to Dr. Darbar (EIA 0940116N), and grants from the National Institutes of Health (NIH) to Dr. Darbar (HL092217), and Dr. Roden (U19 HL65962, and UL1 RR024975). This project was also supported by CTSA award (UL1TR000445) from the National Center for Advancing Translational Sciences. Its contents are solely the responsibility of the authors and do not necessarily represent the official views of the National Center for Advancing Translational Sciences of the NIH.

**NHLBI TOPMed: Novel Risk Factors for the Development of Atrial Fibrillation in Women**

The Women's Genome Health Study (WGHS) is supported by HL 043851 and HL099355 from the National Heart, Lung, and Blood Institute and CA 047988 from the National Cancer Institute, the Donald W. Reynolds Foundation with collaborative scientific support and funding for genotyping provided by Amgen. AF endpoint confirmation was supported by HL-093613 and a grant from the Harris Family Foundation and Watkin's Foundation.

**NHLBI TOPMed: Rare Variants for Hypertension in Taiwan Chinese (THRV)**

The Rare Variants for Hypertension in Taiwan Chinese (THRV) is supported by the National Heart, Lung, and Blood Institute (NHLBI) grant (R01HL111249) and its participation in TOPMed is supported by an NHLBI supplement (R01HL111249-04S1). THRV is a collaborative study between Washington University in St. Louis, LA BioMed at Harbor UCLA, University of Texas in Houston, Taichung Veterans General Hospital, Taipei Veterans General Hospital, Tri-Service General Hospital, National Health Research Institutes, National Taiwan University, and Baylor University. THRV is based (substantially) on the parent SAPPHIRe study, along with additional population-based and hospital-based cohorts. SAPPHIRe was supported by NHLBI grants (U01HL54527, U01HL54498) and Taiwan funds, and the other cohorts were supported by Taiwan funds.

**NHLBI TOPMed: Women's Health Initiative (WHI)**

The WHI program is funded by the National Heart, Lung, and Blood Institute, National Institutes of Health, U.S. Department of Health and Human Services through contracts HHSN268201600018C, HHSN268201600001C, HHSN268201600002C, HHSN268201600003C, and HHSN268201600004C.

**NHLBI TOPMed: BioMe Biobank at Mount Sinai**

The Mount Sinai BioMe Biobank has been supported by The Andrea and Charles Bronfman Philanthropies and in part by Federal funds from the NHLBI and NHGRI (U01HG00638001; U01HG007417; X01HL134588). We thank all participants in the Mount Sinai Biobank. We also thank all our recruiters who have assisted and continue to assist in data collection and management and are grateful for the computational resources and staff expertise provided by Scientific Computing at the Icahn School of Medicine at Mount Sinai.

**NHLBI TOPMed: My Life, Our Future – Genotyping for Progress in Hemophilia**

The My Life, Our Future samples and data are made possible through the partnership of Bloodworks Northwest, the American Thrombosis and Hemostasis Network, the National Hemophilia Foundation, and Bioverativ. We gratefully acknowledge the hemophilia treatment centers and their patients who provided biological samples and phenotypic data.

**NHLBI TOPMed: Outcome Modifying Gene in SCD (OMG-SCD)**
The OMG-SCD study was administrated by Marilyn J. Telen, M.D. and Allison E. Ashley-Koch, Ph.D. from Duke University Medical Center and collection of the data set was supported by grants HL068959 and HL079915 from the National Heart, Lung, and Blood Institute (NHLBI) of the National Institute of Health (NIH).

**NHLBI TOPMed: Treatment of Pulmonary Hypertension and Sickle Cell Disease With Sildenafil Therapy (Walk-PHaSST)**
Special thanks to the volunteers who participated in the Walk-PHaSST study and the investigators of this study. This project was funded with federal funds from the NHLBI, NIH, Department of Health and Human Services, under contract HHSN268200617182C. We also thank the Walk-PHaSST Biorepository at the University of Pittsburgh for their support in this study.

### 3 Other Acknowledgments

Vivien A. Sheehan was supported in part by grant 5K08DK110448-03.
Eimear E. Kenny was supported in part by grant X01HL134588.
Gonçalo R. Abecasis was supported in part by grants U01HL117626, R01HG007022, HHSN268201800002I.
Jessica Lasky-Su was supported in part by grant 1 P01 HL13285.
Lori Garman and Courtney Montgomery were supported in part by grant HL113326.
R. Graham Barr was supported in part by grant R01 HL077612.
Deborah A. Nickerson was supported in part by grants HHSN268201600032I and S10OD021553.
Michelle Daya and Kathleen C. Barnes were supported in part by grant R01HL104608.
Cristen J. Willer was supported in part by grants R01-HL127564, R35-HL135824, and R01-HL142023.
Seung Hoan Choi was the recipient of analysis support program from the TOPMed.
May E. Montasser was supported in part by grants U01 HL137181 and AHA 17GRNT33661168.
Diane Fatkin was supported by the National Health and Medical Research Council of Australia (1074386), with additional support from the Victor Chang Cardiac Research Institute, Estate Late RT Hall, and the Simon Lee Foundation.
Anne-Katrin Emde and Soren Germer were supported in part by a Centers for Common Disease Genomics grant from the National Human Genome Research Institute (UM1HG008901).
Thomas W. Blackwell was supported in part by grants U01HL117626, HHSN268201800002I, and R01HG007022.
Mariza de Andrade was supported in part by grants R01 HL66216 an R01 HL83141.
David J. Van Den Berg was supported in part by grant HHSN268201500017C.
Stella Aslibekyan was supported in part by grant K01HL136700.
Achilleas N. Pitsillides and L Adrienne Cupples were supported in part by grants NO1-HC-25195, HHSN268201500001I, and R01 HL092577-06S1.
Donald W. Bowden and Nicholette D. Palmer were supported in part by grants R01 HL92301, R01 HL67348, R01 NS058700, R01 NS075107, R01 AR48797, R01 DK071891, M01 RR07122, F32 HL085989, and P60 AG10484.
James G. Wilson was supported in part by grant U54GM115428.

James B. Meigs was supported in part by grants U01DK078616 and K24 DK080140.
Daniel E Weeks was supported in part by grants R01 HL093093 and R01 HL1333040.
David D. McManus was supported in part by grant KL2RR031981.
Nancy L. Heard-Costa was supported in part by grants NO1-HC-25195, HHSN268201500001I, and R01HL092577-06S1.
Ani Manichaikul was supported in part by grant R01 HL131565.
Hyun Min Kang was supported in part by grant U01HL137182.
John Blangero and Joanne E. Curran were supported in part by grants HL045522, MH078143, MH078111, MH083824, DK085524, and HL113323.
Dawood Darbar was supported in part by grant R01 HL138737.
Weihong Tang was supported in part by grants R01HL059367, NHLBI contracts: HHSN268201100005C, HHSN268201100006C, HHSN268201100007C, HHSN268201100008C, HHSN268201100009C, HHSN268201100010C, HHSN268201100011C, and HHSN268201100012C.
Pradeep Natarajan was supported in part by grant K08HL140203.
Esteban G. Burchard was supported in part by Sandler Family Foundation, the American Asthma Foundation, the RWJF Amos Medical Faculty Development Program, the Harry Wm. and Diana V. Hind Distinguished Professor in Pharmaceutical Sciences II, and grants R01HL117004, R01HL128439, R01HL135156, X01HL134589, R01ES015794, R21ES24844, P60MD006902, R01MD010443, RL5GM118984, 24RT-0025.
Emelia J. Benjamin was supported in part by grants R01HL128914; 2R01 HL092577; 2U54HL120163, American Heart Association, 18SFRN34110082.
Scott Vrieze was supported in part by grants R01 DA 037904, R01 HG 008983, and R21 DA 040177
Edwin Silverman was supported in part by grants U01 HL089856, U01 HL089897, R01 HL113264, and P01 HL114501.
Sudha Seshadri was supported in part by grants R01 AG054076 and R56 AG029451.
Bruce M. Psaty was supported in part by grants HL120393 and HL130114.
Michael H. Cho was supported in part by grants U01 HL089856 and U01 HL089897.
Ramachandran S. Vasan was supported in part by grants NO1-HC-25195, HHSN268201500001I, R01 HL092577-06S1, Evans Medical Foundation and the Jay and Louis Coffman Endowment from the Department of Medicine, Boston University School of Medicine.

Mina K. Chung was supported in part by grants R01 HL 090620 and R01 HL 111314, NIH National Center for Research Resources for Case Western Reserve University and Cleveland Clinic Clinical and Translational Science Award UL1-RR024989, Cleveland Clinic Department of Cardiovascular Medicine philanthropy research funds, Tomsich Atrial Fibrillation Research Fund.
Lewis C. Becker was supported in part by grants U01 HL72518, HL087698, HL112064, HL11006, HL118356, and M01-RR000052.
Steven A. Lubitz was supported by NIH grant 1R01HL139731 and American Heart Association 18SFRN34250007.
Xiaowen Tian, Brian L. Browning and Sharon R. Browning were supported in part by grant HG005701.
Sebastian Zöllner was supported in part by grant R01 HG005855.
Patrick T. Ellinor was supported in part by grants from the National Institutes of Health (1RO1HL092577, R01HL128914, K24HL105780), American Heart Association (18SFRN34110082), Fondation Leducq (14CVD01), Carol and Roch Hillenbrand and the George L. Nardi, MD, funds at Massachusetts General Hospital.
D.C. Rao was supported in part by grants R01HL111249 and R01HL111249-04S1.
Lu-Chen Weng was supported in part by American Heart Association grant 17POST33660226.
Douglas P. Kiel was supported in part by grants R01 AR072199 and R01 AR041398.
Brian E. Cade was supported in part by grant K01HL135405.
Alexander P. Reiner was supported in part by grants R01HL132947, R01HL136574, R01HL129132, and RO1HL130733.
Adolfo Correa was supported in part by grants HHSN268201800010I, HHSN268201800011I, HHSN268201800012I, HHSN268201800013I, HHSN268201800014I, and HHSN268201800015I.
Kathryn L. Lunetta was supported in part by grants R01 HL092577-10 and 18SFRN34110082.
Stephen T. McGarvey was supported in part by grant HL093093.
Gina M. Peloso was supported in part by grants K01HL125751 and R03HL141439.
Alisa K. Manning was supported in part by grant K01 DK107836 and R03 DK118305.
Brian Custer and Shannon Kelly were supported in part by grant HHSN268201100007I.
Nora Franceschini was supported in part by grants R01-MD012765, R01 DK117445-01A1, and R21-HL140385.
Kenneth M. Rice was supported in part by grants OT3HL142478, R01 HL120393, U01 HL 137162, HHSN268201800001I, and HHSN26800001.
Lawrence F. Bielak, Sharon L.R. Kardia, Patricia A. Peyser, and Jennifer A. Smith were supported in part by grants U10 HL054457, U10 HL054464, U10 HL054481, R01 HL087660, R01 HL085571, and R01 HL119443.
Jiang He was supported in part by grants U01HL072507, R01HL087263, and R01HL090682.

Daniel N. Harris, Michael D. Kessler, Douglas P. Loesch, Amol C. Shetty, Braxton Mitchell, and Timothy D. O'Connor were supported in part by grants U01 HL137181-01, R01 HG002898-09A, R01 HL121007-01, OT3 OD025459-01.
Anna Köttgen was supported in part by grants DFG CRC 1140, CRC 992, and KO 3598/3-1.
Rasika A. Mathias was supported in part by grants U01 HL72518, HL087698, HL112064, HL11006, HL118356, and M01-RR000052.
Ruth J.F. Loos was supported in part by grants X01HL134588, R01DK110113, and R01DK107786.
Zachary A. Szpiech, Raul Torres, and Ryan D. Hernandez were supported in part by grant R01HG007644.
Susan Redline was supported in part by grants R35 HL 135818, HL 046389, and HL 113338.
Yingze Zhang was supported in part by grant HHSN268200617182C.
André Corvelo, Wayne E. Clarke, and Michael C. Zody were supported in part by a Centers for Common Disease Genomics grant from the National Human Genome Research Institute (UM1HG008901) and by the Alfred P. Sloan Foundation.
