## Supplementary Figures and Tables for "Sequencing of 53,831 diverse genomes from the NHLBI TOPMed Program"

**Supplementary Table 1. TOPMed Projects and participating Parent Studies included in genotype data Freeze 5**. See Supplementary Information 1.1.2 for definitions of TOPMed “Projects” and “Parent studies”.

| Project Abbreviation | Project Name | Phenotype Focus[1] | Participating TOPMed Parent Studies[2] |
|---|---|---|---|
| AA_CAC | African American Coronary Artery Calcification | CAC | DHS, GeneSTAR, GENOA, MESA |
| AFGen | Atrial Fibrillation Genetics Consortium | Afib | ARIC, CCAF, HVH, FHS, MGH_AF, Partners, VAFAR, VU_AF, WGHS |
| Amish | Genetics of Cardiometabolic Health in the Amish | HLB | Amish |
| BAGS | Barbados Genetics of Asthma Study | Asthma | BAGS |
| CFS | Cleveland Family Study | HLB, Sleep | CFS |
| COPD | Genetic Epidemiology of COPD | COPD | COPDGene, EOCOPD |
| CRA_CAMP | The Genetic Epidemiology of Asthma in Costa Rica and the Childhood Asthma Management Program | Asthma | CRA |
| FHS | Framingham Heart Study | HLB | FHS |
| GeneSTAR | Genetic Studies of Atherosclerosis Risk | Platelet Aggregation | GeneSTAR |
| GenSalt | Genetic Epidemiology Network of Salt Sensitivity | Hypertension | GenSalt |
| GOLDN | Genetics of Lipid Lowering Drugs and Diet Network | Lipids | GOLDN |
| HyperGEN_GENOA | Hypertension Genetic Epidemiology Network and Genetic Epidemiology Network of Arteriopathy | Hypertension | GENOA, HyperGEN |
| JHS | Jackson Heart Study | HLB | JHS |
| MESA | Multi-Ethnic Study of Atherosclerosis | HLB | MESA |
| PGX_Asthma | Pharmacogenomics of Bronchodilator Response in Minority Children with Asthma | Asthma | GALAII, SAGE |
| SAFS | San Antonio Family Studies | HLB | SAFS |
| Sarcoidosis | Genetics of Sarcoidosis in African Americans | Sarcoidosis | Sarcoidosis |
| SAS | Samoan Adiposity Study | Adiposity | SAS |
| THRV | Taiwan Study of Hypertension using Rare Variants | Hypertension | THRV |
| VTE | Venous Thromboembolism | VTE, HLB | ARIC, CHS, HVH, Mayo_VTE, WHI |
| WHI | Women's Health Initiative | HLB, Stroke, VTE | WHI |

1 - Primary phenotype focus for TOPMed samples. HLB = general heart, lung, and blood. CAC=coronary artery calcification, Afib=atrial fibrillation, VTE=venous thromboembolism. Note some case-only collections are included.

2 – Some TOPMed studies participate in more than one Project. See Supplementary Table 2 for study abbreviations and additional study information

**Supplementary Table 2. Studies that contributed to the Freeze 5 genotype call set being released on dbGaP**. Each study has a dbGaP accession for the TOPMed sequence data and genotypes, while some also have pre-existing Parent study accessions. Phenotypic data are mainly in the Parent accessions, although some are in the TOPMed accessions. See also Supplementary Figures 21 and 22 for information about the ancestral/ethnic and sex composition of each study. The relationships between these studies and their TOPMed project(s) are summarized in Supplementary Table 1. Users may search dbGaP for "NHLBI TOPMed" for an up-to-date view of which TOPMed accessions have been released on dbGaP. Parent study accessions have been released.

| Study/Cohort Abbreviation | TOPMed Accession | TOPMed Study Name[1] ("NHLBI TOPMed:") | Sample Size[2] | Parent Study Accession | Parent Study Design[3] |
|---|---|---|---|---|---|
| Amish | phs000956 | Genetics of Cardiometabolic Health in the Amish | 1,111 | | family/population sample |
| ARIC | phs001211 | Trans-Omics for Precision Medicine Whole Genome Sequencing Project: ARIC | 3,619 | phs000280 | prospective cohort |
| BAGS | phs001143 | The Genetics and Epidemiology of Asthma in Barbados | 1,022 | | family |
| CCAF | phs001189 | Cleveland Clinic Atrial Fibrillation Study | 360 | phs000820 | cross-sectional case-control |
| CFS | phs000954 | The Cleveland Family Study (WGS) | 994 | phs000284 | family |
| CHS | phs001368 | Cardiovascular Health Study | 69 | phs000287 | prospective cohort |
| COPDGene | phs000951 | Genetic Epidemiology of COPD (COPDGene) in the TOPMed Program | 8,909 | phs000179 | case-control, longitudinal follow-up |
| CRA | phs000988 | The Genetic Epidemiology of Asthma in Costa Rica | 1,142 | | family |
| DHS | phs001412 | Diabetes Heart Study African American Coronary Artery Calcification (AA CAC) | 337 | | family/population sample |
| EOCOPD | phs000946 | Boston Early-Onset COPD Study in the TOPMed Program | 74 | phs001161 | family |
| FHS | phs000974 | Whole Genome Sequencing and Related Phenotypes in the Framingham Heart Study | 4,166 | phs000007 | prospective cohort |
| GALAII | phs000920 | Genes-environments and Admixture in Latino Asthmatics (GALA II) Study | 999 | phs001180 | pharmacogenomic |
| GeneSTAR | phs001218 | GeneSTAR (Genetic Study of Atherosclerosis Risk) | 1,637 | phs001074 | family |
| GENOA | phs001345 | Genetic Epidemiology Network of Arteriopathy (GENOA) | 1,143 | phs001238 | family |
| GenSalt | phs001217 | Genetic Epidemiology Network of Salt Sensitivity (GenSalt) | 1,689 | phs000784 | family |

| | | | | | |
|---|---|---|---|---|---|
| GOLDN | phs001359 | Genetics of Lipid Lowering Drugs and Diet Network (GOLDN) | 899 | phs000741 | family |
| HVH | phs000993 | Heart and Vascular Health Study (HVH) | 625 | phs001013 | cross-sectional case-control |
| HyperGEN | phs001293 | HyperGEN - Genetics of Left Ventricular (LV) Hypertrophy | 1,776 | | cross-sectional case-control |
| JHS | phs000964 | Jackson Heart Study | 3,406 | phs000286 | prospective cohort |
| Mayo_VTE | phs001402 | Whole Genome Sequencing of Venous Thromboembolism (WGS of VTE) | 1,251 | phs000289 | cross-sectional case-control |
| MESA | phs001416 | MESA and MESA Family AA-CAC | 4,875 | phs000209 | prospective cohort |
| MGH_AF | phs001062 | MGH Atrial Fibrillation Study | 984 | phs001001 | family |
| Partners | phs001024 | Partners HealthCare Biobank | 128 | | cross-sectional case-control |
| SAFS | phs001215 | San Antonio Family Heart Study (WGS) | 1,508 | | family |
| SAGE | phs000921 | Study of African Americans, Asthma, Genes and Environment (SAGE) Study | 499 | | pharmacogenomic |
| Sarcoidosis | phs001207 | African American Sarcoidosis Genetics Resource | 606 | | family and cross-sectional case-control |
| SAS | phs000972 | Genome-wide Association Study of Adiposity in Samoans | 1,232 | phs000914 | population sample |
| THRV | phs001387 | Rare Variants for Hypertension in Taiwan Chinese (THRV) | 1,525 | | case families and controls |
| VAFAR | phs000997 | The Vanderbilt AF Ablation Registry | 163 | | cases with longitudinal follow-up |
| VU_AF | phs001032 | The Vanderbilt Atrial Fibrillation Registry | 1,110 | | families with longitudinal follow-up |
| WGHS | phs001040 | Novel Risk Factors for the Development of Atrial Fibrillation in Women | 115 | | prospective cohort |
| WHI | phs001237 | Women's Health Initiative (WHI) | 10,047 | phs000200 | prospective cohort |

1 – Study name as it appears in dbGaP, with “NHLBI TOPMed:” prepended
2 – Approximate sample size for freeze4 and freeze5 releases combined

**Supplementary Table 3: Number of exonic SNVs in Framingham Heart Study.**
Categories are not mutually exclusive.

| Study | Exonic SNVs | LOF | Missense | Non-synonymous |
|---|---|---|---|---|
| TOPMed | 212,603 | 2,047 | 125,862 | 127,834 |
| CHARGE WGS | 182,165 | 1,581 | 105,588 | 107,283 |
| CHARGE WES | 180,520 | 1,742 | 108,481 | 109,849 |

**Supplementary Table 4. Count and percent of singletons and indels in regions of different function.**

| Category | Singletons (%, CI) | Indels (%) | All Variants |
|---|---|---|---|
| CTCF Binding Sites | 4,348,892 (45.42, [45.40, 45.43]) | 570,081 (5.95, [5.95, 5.96]) | 9,575,270 |
| TF Binding Sites | 1,161,714 (45.68, [45.65, 45.71]) | 164,715 (6.48, [6.46, 6.49]) | 2,543,261 |
| Intergenic | 80,323,798 (45.89, [45.89, 45.90]) | 11,877,557 (6.79, [6.78, 6.79]) | 175,023,248 |
| Enhancers | 803,583 (45.91, [45.87, 45.95]) | 119,949 (6.85, [6.83, 6.87]) | 1,750,311 |
| Genome | 188,947,391 (46.05, [46.05, 46.05]) | 28,980,753 (7.06, [7.06, 7.06]) | 410,323,831 |
| Intronic | 78,225,277 (46.06, [46.06, 46.06]) | 12,576,976 (7.41, [7.40, 7.41]) | 169,833,412 |
| 3' UTR | 2,491,324 (46.10, [46.08, 46.12]) | 516,744 (9.56, [9.55, 9.57]) | 5,404,168 |
| Open Chromatin | 4,619,783 (46.22, [46.21, 46.24]) | 652,279 (6.53, [6.52, 6.53]) | 9,994,905 |
| CDS | 2,235,254 (46.79, [46.77, 46.82]) | 167,427 (3.50, [3.50, 3.51]) | 4,776,820 |
| 5' UTR | 872,250 (47.28, [47.25, 47.32]) | 121,980 (6.61, [6.59, 6.63]) | 1,844,774 |
| Promoters | 515,168 (47.72, [47.67, 47.77]) | 74,684 (6.92, [6.89, 6.94]) | 1,079,553 |

**Supplementary Table 5. Putative loss of function variants per individual in TOPMed Freeze 5 and ExAC data sets.** To compare number of putative loss of function (pLoF) variants per individual, we used only rare (AF < 0.5%) bi-allelic variants which were not present in dbSNP build 142 (last dbSNP database version without ExAC variants).

| | ExAC | | | | TOPMed Freeze 5 | | | |
|---|---|---|---|---|---|---|---|---|
| | % Singletons | Per Individual | Singletons | Total | % Singletons | Per Individual | Singletons | Total |
| LoF | | | | | | | | |
| all | 73.00 | 5.51 | 114,621 | 157,006 | 63.90 | 7.83 | 110,815 | 173,428 |
| frameshift | 70.94 | 2.81 | 46,027 | 64,883 | 62.27 | 4.51 | 53,861 | 86,498 |
| splice | 76.25 | 1.09 | 28,596 | 37,501 | 66.76 | 1.39 | 24,167 | 36,198 |
| stop_gained | 73.23 | 1.61 | 39,998 | 54,622 | 64.63 | 1.94 | 32,787 | 50,732 |
| Coding | | | | | | | | |
| all | 66.71 | 116.96 | 2,088,105 | 3,130,064 | 59.43 | 129.63 | 1,700,286 | 2,861,087 |
| inframe | 59.30 | 1.49 | 13,972 | 23,561 | 51.41 | 4.13 | 23,083 | 44,899 |
| missense | 67.62 | 72.88 | 1,390,186 | 2,056,022 | 60.24 | 79.17 | 1,109,809 | 1,842,262 |
| synonymous | 64.24 | 38.37 | 604,628 | 941,154 | 57.48 | 39.30 | 478,827 | 833,050 |

**Supplementary Table 6. Enrichment and depletion of putative loss-of-function (pLoF) variants in gene sets.** For each genes set we computed number of rare (AF < 0.5%) pLoF variants per coding sequence base pair (pLoF/bp) and proportion of singletons. We compared observed pLoF/bp and proportion of singletons to 1,000,000 randomly sampled gene sets of same size and coding sequence length. P-value significance threshold is $2x10^{-6}$. BP - biological process, MF - molecular function, CC - cellular component.

| Gene Sets | | No. of Genes | pLoF / bp | | | | Proportion of Singletons | | | |
|---|---|---|---|---|---|---|---|---|---|---|
| | | | Observed | Sample Mean | Ratio | P-value | Observed | Sample Mean | Ratio | P-value |
| *Gene Ontology (GO)* | | | | | | | | | | |
| Class | Term | | | | | | | | | |
| MF | (GO:0043565) sequence-specific DNA binding | 596 | 0.0033 | 0.0072 | 0.46 | $<1x10^{-6}$ | 0.4956 | 0.4767 | 1.04 | $<1x10^{-6}$ |
| BP | (GO:0006413) translational initiation | 142 | 0.0045 | 0.0083 | 0.55 | $<1x10^{-6}$ | 0.5171 | 0.4803 | 1.08 | $3x10^{-6}$ |
| BP | (GO:0008380) RNA splicing | 283 | 0.0048 | 0.0071 | 0.67 | $<1x10^{-6}$ | 0.5056 | 0.4772 | 1.06 | $<1x10^{-6}$ |
| BP | (GO:0006397) mRNA processing | 359 | 0.0048 | 0.0071 | 0.67 | $<1x10^{-6}$ | 0.5085 | 0.4772 | 1.07 | $<1x10^{-6}$ |
| BP | (GO:0006357) regulation of transcription by RNA polymerase II | 768 | 0.0043 | 0.0069 | 0.62 | $<1x10^{-6}$ | 0.4911 | 0.4766 | 1.03 | $<1x10^{-6}$ |
| MF | (GO:0003700) DNA binding transcription factor activity | 996 | 0.0048 | 0.0070 | 0.68 | $<1x10^{-6}$ | 0.4900 | 0.4763 | 1.03 | $<1x10^{-6}$ |
| CC | (GO:0005654) nucleoplasm | 3,135 | 0.0055 | 0.0070 | 0.79 | $<1x10^{-6}$ | 0.4914 | 0.4768 | 1.03 | $<1x10^{-6}$ |
| CC | (GO:0030529) intracellular ribonucleoprotein complex | 351 | 0.0063 | 0.0082 | 0.77 | $<1x10^{-6}$ | 0.5000 | 0.4783 | 1.05 | $<1x10^{-6}$ |
| MF | (GO:0003723) RNA binding | 1,491 | 0.0057 | 0.0071 | 0.79 | $<1x10^{-6}$ | 0.4930 | 0.4771 | 1.03 | $<1x10^{-6}$ |
| BP | (GO:0006351) transcription, DNA-templated | 2,364 | 0.0052 | 0.0068 | 0.75 | $<1x10^{-6}$ | 0.4875 | 0.4764 | 1.02 | $<1x10^{-6}$ |
| BP | (GO:0006355) regulation of transcription, DNA-templated | 2,537 | 0.0052 | 0.0069 | 0.76 | $<1x10^{-6}$ | 0.4874 | 0.4764 | 1.02 | $<1x10^{-6}$ |
| CC | (GO:0005634) nucleus | 6,262 | 0.0061 | 0.0071 | 0.86 | $<1x10^{-6}$ | 0.4863 | 0.4769 | 1.02 | $<1x10^{-6}$ |
| CC | (GO:0005886) plasma membrane | 4,620 | 0.0065 | 0.0070 | 0.94 | $<1x10^{-6}$ | 0.4713 | 0.4766 | 0.99 | $<1x10^{-6}$ |
| BP | (GO:0055114) oxidation-reduction process | 704 | 0.0094 | 0.0074 | 1.27 | $<1x10^{-6}$ | 0.4620 | 0.4772 | 0.97 | $<1x10^{-6}$ |
| MF | (GO:0016491) oxidoreductase activity | 577 | 0.0093 | 0.0072 | 1.29 | $<1x10^{-6}$ | 0.4618 | 0.4767 | 0.97 | $<1x10^{-6}$ |
| *Public Databases* | | | | | | | | | | |
| COSMIC genes | | 916 | 0.0048 | 0.0069 | 0.69 | $<1x10^{-6}$ | 0.4846 | 0.4761 | 1.018 | $9x10^{-5}$ |
| GWAS Catalog upstream genes | | 1,917 | 0.0068 | 0.0075 | 0.91 | $<1x10^{-6}$ | 0.4773 | 0.4766 | 1.001 | $3x10^{-1}$ |
| GWAS Catalog downstream genes | | 1,944 | 0.0069 | 0.0075 | 0.92 | $<1x10^{-6}$ | 0.4764 | 0.4767 | 0.999 | $4x10^{-1}$ |

| | | | | | | | | | |
|---|---|---|---|---|---|---|---|---|---|
| GWAS Catalog genes | 5,179 | 0.0067 | 0.0070 | 0.96 | $4x10^{-5}$ | 0.4736 | 0.4761 | 0.995 | $3x10^{-3}$ |
| ClinVar genes with ≥1 pathogenic variants | 3,893 | 0.0068 | 0.0071 | 0.96 | $8x10^{-4}$ | 0.4717 | 0.4762 | 0.990 | $2x10^{-5}$ |
| ClinVar genes without pathogenic variants | 2,103 | 0.0072 | 0.0070 | 1.02 | $7x10^{-2}$ | 0.4703 | 0.4762 | 0.988 | $3x10^{-5}$ |
| OMIM genes | 4,335 | 0.0069 | 0.0072 | 0.96 | $5x10^{-5}$ | 0.4713 | 0.4763 | 0.990 | $<1x10^{-6}$ |

**Supplementary Table 7. Unique *CYP2D6* alleles detected by Stargazer, both known and novel, in 1,923 unrelated African American individuals (3,846 allele calls) from the Jackson Heart Study.**

| # | Allele | Activity score | Major variation | Count | % |
|---|---|---|---|---|---|
| 1 | **1* | 1 | Reference | 1155 | 30.03 |
| 2 | **1x2* | 2 | Gene duplication | 31 | 0.81 |
| 3 | **1x3*[1] | 3 | Gene multiplication | 1 | 0.03 |
| 4 | **2* | 1 | 1661G>C, 2850C>T, 4180G>C | 628 | 16.33 |
| 5 | **2x2* | 2 | Gene duplication | 38 | 0.99 |
| 6 | **2x3*[1] | 3 | Gene multiplication | 1 | 0.03 |
| 7 | **3* | 0 | 2549delA | 14 | 0.36 |
| 8 | **4* | 0 | 1846G>A | 119 | 3.09 |
| 9 | **4x2* | 0 | Gene duplication | 124 | 3.22 |
| 10 | **4N+*4* | 0 | *CYP2D6/CYP2D7* hybrid | 5 | 0.13 |
| 11 | **5* | 0 | Gene deletion | 211 | 5.49 |
| 12 | **6* | 0 | 1707delT | 4 | 0.1 |
| 13 | **7* | 0 | 2935A>C | 1 | 0.03 |
| 14 | **9* | 0.5 | 2615_2617delAAG | 19 | 0.49 |
| 15 | **10* | 0.5 | 100C>T | 160 | 4.16 |
| 16 | **10x2* | 1 | Gene duplication | 1 | 0.03 |
| 17 | **12* | 0 | 124G>A | 4 | 0.1 |
| 18 | **15* | 0 | 137_138insT | 1 | 0.03 |
| 19 | **17* | 0.5 | 1023C>T | 610 | 15.86 |
| 20 | **17x2* | 1 | Gene duplication | 1 | 0.03 |
| 21 | **19* | 0 | 2539_2542delAACT | 1 | 0.03 |
| 22 | **22* | Unknown | 82C>T | 3 | 0.08 |
| 23 | **27* | 1 | 3853G>A | 6 | 0.16 |
| 24 | **28* | Unknown | 19G>A, 1704C>G, 2850C>T, 4180G>C | 2 | 0.05 |
| 25 | **29* | 0.5 | 1659G>A, 2850C>T, 3183G>A, 4180G>C | 294 | 7.64 |
| 26 | **29x2* | 1 | Gene duplication | 12 | 0.31 |
| 27 | **29x3*[1] | 1.5 | Gene multiplication | 1 | 0.03 |
| 28 | **30* | Unknown | 1863_1864insTTTCGCCCC | 1 | 0.03 |
| 29 | **31* | 0 | 2850C>T, 4042G>A, 4180G>C | 1 | 0.03 |
| 30 | **32* | Unknown | 1661G>C, 2850C>T, 3853G>A, 4180G>C | 2 | 0.05 |

| | | | | | |
|---|---|---|---|---|---|
| 31 | **33* | 1 | 2483G>T | 7 | 0.18 |
| 32 | **34* | 1 | 2850C>T | 11 | 0.29 |
| 33 | **34x2*[1] | 2 | Gene duplication | 7 | 0.18 |
| 34 | **35* | 1 | 31G>A, 1661G>C, 2850C>T, 4180G>C | 36 | 0.94 |
| 35 | **36+*10* | 0.5 | *CYP2D6/CYP2D7* hybrid | 7 | 0.18 |
| 36 | **39* | 1 | 1661G>C, 4180G>C | 4 | 0.1 |
| 37 | **40* | 0 | 1023C>T, 1863_1864insTTTCGCCCCTTTCGCCCC, 2850C>T, 4180G>C | 20 | 0.52 |
| 38 | **41* | 0.5 | 2988G>A | 109 | 2.83 |
| 39 | **41x2* | 1 | Gene duplication | 4 | 0.1 |
| 40 | **42* | 0 | 3260_3261insTG | 11 | 0.29 |
| 41 | **42x2*[1] | 0 | Gene duplication | 1 | 0.03 |
| 42 | **43* | 1 | 77G>A | 31 | 0.81 |
| 43 | **43x2* | 2 | Gene duplication | 2 | 0.05 |
| 44 | **46* | 1 | 77G>A, 1716G>A, 2850C>T, 4180G>C | 19 | 0.49 |
| 45 | **50* | 0.5 | 1720A>C | 1 | 0.03 |
| 46 | **53* | 1 | 1611T>A, 1617G>T | 1 | 0.03 |
| 47 | **56* | 0 | 3201C>T | 10 | 0.26 |
| 48 | **59* | 0.5 | 1661G>C, 2291G>A, 2850C>T, 2939G>A, 3384A>C, 3584G>A, 4180G>C | 2 | 0.05 |
| 49 | **66* | 0 | *CYP2D6/CYP2D7* hybrid | 3 | 0.08 |
| 50 | **68+*4* | 0 | *CYP2D6/CYP2D7* hybrid | 31 | 0.81 |
| 51 | **76* | 0 | *CYP2D6/CYP2D7* hybrid | 1 | 0.03 |
| 52 | **77+*2* | 1 | *CYP2D6/CYP2D7* hybrid | 1 | 0.03 |
| 53 | **78+*2* | 1 | *CYP2D6/CYP2D7* hybrid | 2 | 0.05 |
| 54 | **81* | 0 | 2579C>T | 2 | 0.05 |
| 55 | **84* | 0.5 | 2574C>A, 2850C>T, 4180G>C | 13 | 0.34 |
| 56 | **108* | Unknown | 3226A>G, 3235A>G | 1 | 0.03 |
| 57 | Unknown | - | - | 58 | 1.51 |

[1]Novel gene duplication or multiplication.

**Supplementary Table 8. Average heterozygosity and number of singletons by TOPMed study and self-reported ancestry.**

| Study | Ancestry | Sample Size | Average Heterozygosity | Average Singletons |
|---|---|---|---|---|
| Amish | Amish | 225 | 2022666.44 | 425.22 |
| AFGen | European | 2572 | 2115260.32 | 4105.61 |
| ARIC | European | 3150 | 2109648.87 | 3950.55 |
| CFS | European | 182 | 2114570.49 | 4648.35 |
| CHS | European | 53 | 2109758.51 | 3876.19 |
| COPDGene | European | 5653 | 2115083.01 | 4359.24 |
| FHS | European | 1262 | 2114087.25 | 4929.14 |
| GeneSTAR | European | 340 | 2113552.12 | 4634.87 |
| GOLDN | European | 280 | 2107030.61 | 4385.82 |
| Mayo_VTE | European | 1161 | 2108846.87 | 4626.79 |
| MESA | European | 1621 | 2117335.35 | 3972.58 |
| WHI | European | 8101 | 2112381.22 | 4240.63 |
| GALAII | Hispanic/Latino | 891 | 2279532.13 | 5426.62 |
| MESA | Hispanic/Latino | 828 | 2293654.33 | 6827.96 |
| SAFS | Hispanic/Latino | 443 | 2167178.99 | 5863.18 |
| WHI | Hispanic/Latino | 273 | 2225397.04 | 6685.21 |
| ARIC | African | 186 | 2815360 | 4820.82 |
| BAGS | African | 383 | 2814073.26 | 6248.38 |
| CFS | African | 157 | 2809913.25 | 5113.15 |
| COPDGene | African | 2646 | 2814045.84 | 5320.36 |
| GeneSTAR | African | 270 | 2808915.3 | 5145.25 |
| GENOA | African | 331 | 2816393.91 | 4420.97 |
| HyperGEN | African | 855 | 2820505.11 | 5157.84 |
| JHS | African | 1897 | 2813758.33 | 4644.06 |
| MESA | African | 1061 | 2800932.63 | 5268.29 |
| SAGE | African | 437 | 2811047.94 | 5477.55 |
| Sarcoidosis | African | 236 | 2811252.67 | 5413.35 |
| WHI | African | 1286 | 2792439.3 | 5075.41 |
| GenSalt | Asian | 677 | 1972061.56 | 13954.51 |
| MESA | Asian | 509 | 1968481.72 | 13874.66 |
| WHI | Asian | 176 | 1980241.1 | 14752.07 |
| SAS | Samoan | 962 | 1907790.71 | 2415.64 |

**Supplementary Table 9. Resulting fitted parameters from performing demographic inference with 4-fold degenerate sites and sites under the weakest effects of selection at linked sites (SaLS).** Weakest SaLS represent sites from the highest 1% *B* bin (McVicker's *B* statistic). Parameters include the starting population size before growth ($NEur_0$), the ending population size after growth (*NEur*), and the time span over which exponential growth occurred (*TEur*). The rate of growth is shown in the last column (rEur). Population size is in given in units of $N_e$ and time is given in years assuming a generation time of 25 years.

| sample size (2N) | data | $NEur_0$ (95% CI) | *NEur* (95% CI) | *TEur* (95% CI) | rEur*100 (95% CI) |
|---|---|---|---|---|---|
| 1000 | weakest SaLS | 11,689 (11,586-11,792) | 1,163,554 (971,534-1,355,573) | 5,741 (5,566-5,916) | 2.02 (1.94-2.09) |
| | 4-fold degenerate | 10,059 (9,902-10,217) | 755,334 (644,947-865,722) | 8,505 (8,131-8,880) | 1.28 (1.23-1.32) |
| 2000 | weakest SaLS | 11,762 (11,658-11,866) | 1,053,407 (974,047-1,132,768) | 5,813 (5,682-5,943) | 1.95 (1.92-1.98) |
| | 4-fold degenerate | 10,313 (10,159-10,466) | 949,761 (867,911-1,031,612) | 8,058 (7,802-8,314) | 1.41 (1.38-1.44) |
| 3000 | weakest SaLS | 11,835 (11,735-11,936) | 1,060,540 (1,003,916-1,117,164) | 5,793 (5,687-5,899) | 1.96 (1.93-1.98) |
| | 4-fold degenerate | 10,487 (10,334-10,639) | 1,043,842 (976,228-1,111,456) | 7,853 (7,634-8,072) | 1.48 (1.45-1.5) |
| 4000 | weakest SaLS | 11,895 (11,790-12,000) | 1,065,695 (1,022,542-1,108,848) | 5,778 (5,676-5,879) | 1.96 (1.95-1.98) |
| | 4-fold degenerate | 10,619 (10,462-10,775) | 1,094,072 (1,035,292-1,152,852) | 7,738 (7,524-7,952) | 1.51 (1.49-1.53) |
| 4832 | weakest SaLS | 11,937 (11,716-12,158) | 1,066,503 (1,029,592-1,103,414) | 5,770 (5,600-5,940) | 1.97 (1.95-1.98) |
| | 4-fold degenerate | 10,709 (9,469-11,948) | 1,118,240 (1,033,993-1,202,487) | 7,677 (6,416-8,938) | 1.53 (1.5-1.55) |

**Supplementary Table 10.** The genes found associated with strong evidence for positive selection in all six analyzed populations, along with their location, function, and reported disease associations.

| Gene | Location (hg19) | Function | Disease Associations |
|---|---|---|---|
| *PIGG* | chr4:492988-533710 | encodes an enzyme involved in glycosylphosphatidylinositol-anchor biosynthesis (Shishioh et al. 2005) | intellectual disability, hypotonia, and early-onset seizures |
| *SLC30A9* | chr4:41992522-42089551 | zinc transporter (Sim and Chow 1999) | Birk-Landau-Perez syndrome |
| *STXBP5L* | chr3:120627049-121143608 | vesicular trafficking, neurotransmitter release (Kumar et al. 2015) | neurodegenerative disorder |

**Supplementary Table 11.** Distribution of variants in the TOPMed imputation panel in non-reference allele frequency bins.

| Variation type | Non-reference allele frequency bins | | | | |
|---|---|---|---|---|---|
| | (0, 0.005] | (0.005, 0.01] | [0.01, 0.05) | [0.05, 1) | Totals |
| **SNVs** | 208,013,118 | 3,258,174 | 5,010,445 | 6,730,611 | 223,012,348 |
| **Insertions** | 4,234,716 | 53,957 | 78,732 | 100,158 | 4,467,563 |
| **Deletions** | 11,557,580 | 173,315 | 257,782 | 287,559 | 12,276,236 |
| **Totals** | 223,805,414 | 3,485,446 | 5,346,959 | 7,118,328 | 239,756,147 |

**Supplementary Table 12**. Rare LoF disease-associated variants identified in the UK Biobank not present in the HRC-imputed UK Biobank public release. AN - non-reference allele frequency. $R^2$ - Minimac4 imputation quality metric. OR and corresponding 95% confidence interval are reported from using the Firth test on the unrelated subset of the UK Biobank white British (no third degree relative pairs or closer) because SAIGE's estimates of effect size are unstable for rare variants for traits with a small number of cases.

| Trait | Cases | | Controls | | CHR:BP:REF:ALT | $R^2$ | P-value | OR | 95% CI | Status in ClinVar | Gene |
|---|---|---|---|---|---|---|---|---|---|---|---|
| | N | AF (%) | N | AF (%) | | | | | | | |
| Breast cancer | 12,636 | 0.55% | 200,417 | 0.21% | 22:28695868:AG:A | 0.68 | 2.4E-21 | 2.09 | 1.52-2.87 | Pathogenic for familial breast cancer | *CHEK2* |
| Breast cancer | 12,636 | 0.15% | 200,417 | 0.04% | 16:23621362:C:T | 0.96 | 1.9E-13 | 4.39 | 2.90-6.63 | Pathogenic/ likely pathogenic for familial breast cancer | *PALB2* |
| Hematuria | 16,379 | 0.34% | 379,181 | 0.05% | 2:227052367:G:C | 0.99 | 8.2E-49 | 7.03 | 5.57-8.87 | Pathogenic for Alport's disease (note: key symptom is hematuria) | *COL4A4* |
| Hereditary hemolytic anemias | 156 | 0.96% | 389,249 | 0.00167 % | 11:5227001:CT:C | 0.73 | 9.2E-09 | 706 | 201-2480 | Pathogenic for Beta thalassemia | *HBB* |

**Supplementary Table 13. Centers that have provided genomic assays to TOPMed.**

| Center name | Principal Investigator | Assay type(s) |
|---|---|---|
| Baylor Human Genome Sequencing Center | Richard Gibbs | WGS |
| Broad Institute Genomics Platform | Stacey Gabriel | WGS, RNA-seq |
| McDonnell Genome Institute | Susan K. Dutcher | WGS |
| Illumina | Karine A. Viaud-Martinez | WGS |
| Macrogen | Sal Situ | WGS |
| New York Genome Center | Soren Germer | WGS |
| Northwest Genomics Center | Deborah Nickerson | WGS, RNA-seq |
| Beth Israel Deaconess Medical Center | Robert E. Gerszten | Proteomics, Metabolomics |
| Broad Institute Metabolomics Platform | Clary Clish | Metabolomics |
| Keck Molecular Genomics Core Facility | David Van Den Berg | Methylomics |

**Supplementary Table 14. TOPMed study-consent groups used in analyses/tools**. Consent group data use limitations: GRU - general research use; HMB - limited to health/medical/biomedical purposes; DS - use of the data must be related to specified disease. Consent group data use limitation modifiers: IRB - requestor must provide documentation of local IRB approval; PUB - requestor agrees to make results of studies using the data available to the larger scientific community; COL - requestor must provide a letter of collaboration with the primary study investigator(s); NPU - use of the data is limited to not-for-profit organizations; MDS - use of the data includes methods development research; GSO - use of the data is limited to genetic studies only. Consent group disease abbreviations: AF - Atrial Fibrillation; ASC-RF - Arteriosclerosis and its Risk Factors; CVD - Cardiovascular Disease; CS - COPD and Smoking; DHD - Diabetes and Heart Disease; HLBS - Heart, Lung, Blood, and Sleep disorders; LD - Lung Diseases; FDO - Focus Disease Only (in JHS, FDO=blood pressure, heart/CVD, obesity, diabetes, kidney disease, or lung disease and risk factors); RD - Related Disorders; SAR - Sarcoidosis; SCD - Sickle Cell Disease.

| Study/Cohort Abbreviation | TOPMed Accession | Consent Group | Freeze 5 VCF | | | | | Freeze 7 VCF | Freeze 3 VCF | BAM |
|---|---|---|---|---|---|---|---|---|---|---|
| | | | PCA, Kinship | General variant analyses | Population genetics | BRAVO variant server | Imputation reference panel | Imputation accuracy | Selection/Adaptation, Demography | De novo assembly |
| Amish | phs000956 | HMB-IRB-MDS | X | X | X | X | X | | | X |
| ARIC | phs001211 | DS-CVD-IRB | X | X | X | X | X | | X | |
| | | HMB-IRB | X | X | X | X | X | | X | X |
| AustralianFamilialAF | phs001435 | HMB-NPU-MDS | | | | X | X | | | |
| BAGS | phs001143 | GRU-IRB | X | X | X | X | X | | X | X |
| BioMe | phs001644 | HMB-NPU | | | | | | X | | |
| CCAF | phs001189 | GRU-IRB | X | X | X | X | X | | X | X |
| CFS | phs000954 | DS-HLBS-IRB-NPU | X | X | X | X | | | X | X |
| CHS | phs001368 | HMB-NPU-MDS | X | X | X | X | X | | | |
| COPDGene | phs000951 | HMB | X | X | X | X | X | | X | X |
| | | DS-CS-RD | X | X | | | | | | |

| | | | | | | | | | | | | | | |
|---|---|---|---|---|---|---|---|---|---|---|---|---|---|---|
| CRA | phs000988 | DS-ASTHMA-IRB-MDS-RD | X | X | | X | X | | | | | | | X |
| DECAF | phs001546 | GRU | | | | X | X | | | | | | | |
| DHS | phs001412 | DS-DHD-IRB-COL-NPU | X | X | | X | X | | | | | | | |
| | | HMB-IRB-COL-NPU | X | X | | X | X | | | | | | | |
| EOCOPD | phs000946 | DS-CS-RD | X | X | | | | | | | | | | X |
| FHS | phs000974 | HMB-IRB-MDS | X | X | X | X | X | | | | | X | | X |
| | | HMB-IRB-NPU-MDS | X | X | X | X | X | | | | | X | | X |
| GALAII | phs000920 | DS-LD-RD | X | X | X | X | | | | | | X | | X |
| GeneSTAR | phs001218 | DS-CVD-IRB-NPU-MDS | X | X | X | X | X | | | | | | | |
| GENOA | phs001345 | DS-ASC-RF-NPU | X | X | X | X | X | | | | | | | |
| GenSalt | phs001217 | GRU-IRB | X | X | X | X | X | | | | | | | |
| GOLDN | phs001359 | DS-CVD-IRB | X | X | X | X | X | | | | | | | |
| HVH | phs000993 | HMB-IRB-MDS | X | X | X | X | X | | | | | X | | X |
| | | DS-CVD-IRB-MDS | X | X | X | X | X | | | | | X | | X |
| HyperGEN | phs001293 | DS-CVD-IRB-RD | X | X | X | X | X | | | | | | | |
| HyperGEN | phs001293 | GRU-IRB | X | X | X | X | X | | | | | | | |
| JHS | phs000964 | HMB-IRB | X | X | X | X | X | | | | | X | | X |
| | | HMB-IRB-NPU | X | X | X | X | X | | | | | X | | X |
| | | DS-FDO-IRB | X | X | X | X | X | | | | | X | | X |
| | | DS-FDO-IRB-NPU | X | X | X | X | X | | | | | X | | X |
| Mayo_VTE | phs001402 | GRU | X | X | X | X | X | | | | | | | |
| MESA | phs001416 | HMB | X | X | X | X | X | | | | | | | |
| | | HMB-NPU | X | X | X | X | X | | | | | | | |
| MGH_AF | phs001062 | HMB-IRB | X | X | X | X | X | | | | | X | | X |
| | | DS-AF-IRB-RD | X | X | X | X | X | | | | | X | | X |
| miRhythm | phs001434 | GRU | | | | X | X | | | | | | | |
| MLOF | phs001515 | HMB-PUB | | | | X | X | | | | | | | |
| OMG_SCD | phs001608 | DS-SCD-IRB-PUB-COL-MDS-RD | | | | X | X | | | | | | | |
| Partners | phs001024 | HMB | X | X | X | X | X | | | | | X | | X |
| PharmHU | phs001466 | HMB | | | | X | X | | | | | | | |
| REDS-III_Brazil | phs001468 | GRU-IRB-PUB-NPU | | | | X | X | | | | | | | |

| | | | | | | | | | | |
|---|---|---|---|---|---|---|---|---|---|---|
| SAFS | phs001215 | DS-DHD-IRB-PUB-MDS-RD | X | X | X | X | X | | | |
| SAGE | phs000921 | DS-LD | X | X | X | X | | | X | X |
| Sarcoidosis | phs001207 | DS-SAR-IRB | X | X | X | X | X | | | |
| SARP | phs001446 | GRU | | | | X | X | | | |
| SAS | phs000972 | GRU-IRB-PUB-COL-NPU-GSO | X | X | X | | | | | |
| THRV | phs001387 | DS-CVD-IRB-COL-NPU-RD | X | | | X | X | | | |
| VAFAR | phs000997 | HMB-IRB | X | X | X | X | X | | | X |
| VU_AF | phs001032 | GRU-IRB | X | X | X | X | X | | X | X |
| walk_PHaSST | phs001514 | DS-SCD-IRB-PUB-COL-NPU-MDS-RD | | | | X | X | | | |
| | | HMB-IRB-PUB-COL-NPU-MDS-GSO | | | | X | X | | | |
| WGHS | phs001040 | HMB-IRB | X | X | X | X | X | | | X |
| WHI | phs001237 | HMB-IRB | X | X | X | X | X | | | |
| | | HMB-IRB-NPU | X | X | X | X | X | | | |

**Supplementary Table 15. Description of samples in Framingham Heart Study.**

| Sample | Sequencing Center | Sample Size | Depth |
|---|---|---|---|
| TOPMed WGS | Broad Institute | 4,158 | >30X |
| CHARGE WGS | Baylor HGSC | 855 | 6-7X |
| CHARGE WES | Baylor HGSC | 1,702 | >30X |

**Supplementary Table 16. Population clusters in the iHS analysis.** Population clusters identified by k-means clustering, with total number of individuals, total number of unrelated individuals, and total number of unrelated and consented individuals per population, along with population label assignments.

| Population | Label Assignment | N | N Unrelated | N Unrelated and Consented |
|---|---|---|---|---|
| 1 | European A | 6,474 | 4,488 | 3,288 |
| 2 | Puerto Rican | 487 | 477 | 475 |
| 3 | Amish | 1,111 | 231 | 0 |
| 4 | Costa Rican | 1,062 | 567 | 0 |
| 5 | European B | 301 | 215 | 159 |
| 6 | African Admixed | 6,330 | 4,186 | 3,323 |
| 7 | Samoan | 384 | 363 | 0 |
| 8 | Mexican | 504 | 490 | 489 |
| 9 | European C | 1,581 | 742 | 643 |
| | TOTAL | 18,234 | 11,759 | 8,377 |

**Supplementary Table 17. Total number of sites analyzed per population in the iHS analysis.**

| Population | Label Assignment | Sites Analyzed |
|---|---|---|
| 1 | European A | 4,690,744 |
| 2 | Puerto Rican | 5,218,879 |
| 5 | European B | 4,766,815 |
| 6 | African Admixed | 6,543,573 |
| 8 | Mexican | 4,672,492 |
| 9 | European C | 4,741,500 |

**Supplementary Table 18. Funding sources for each study and genomic center in TOPMed**. Whole genome sequencing support for TOPMed studies was provided by the National Heart, Lung, and Blood Institute (NHLBI) through the Centralized Omics REsource (CORE) program. NYGC = New York Genome Center; BROAD = Broad Institute of MIT and Harvard; UW NWGC = University of Washington Northwest Genomics Center; ILLUMINA = Illumina Genomic Services; MACROGEN = Macrogen Corp.; BAYLOR = Baylor Human Genome Sequencing Center.

| Study Accession | TOPMed Parent Study Name | Sequencing Center | Sequencing Support |
|---|---|---|---|
| phs000956 | NHLBI TOPMed: Genetics of Cardiometabolic Health in the Amish | BROAD | 3R01HL121007-01S1 |
| phs001211 | NHLBI TOPMed: Trans-Omics for Precision Medicine Whole Genome Sequencing Project: ARIC | BAYLOR, BROAD | 3R01HL092577-06S1, HHSN268201500015C, 3U54HG003273-12S2 |
| phs001143 | NHLBI TOPMed: The Genetics and Epidemiology of Asthma in Barbados | ILLUMINA | 3R01HL104608-04S1 |
| phs001189 | NHLBI TOPMed: Cleveland Clinic Atrial Fibrillation Study | BROAD | 3R01HL092577-06S1 |
| phs000954 | NHLBI TOPMed: The Cleveland Family Study (WGS) | UW NWGC | 3R01HL098433-05S1 |
| phs001368 | NHLBI TOPMed: Cardiovascular Health Study | BAYLOR | HHSN268201500015C |
| phs000951 | NHLBI TOPMed: Genetic Epidemiology of COPD (COPDGene) in the TOPMed Program | BROAD, UW NWGC | HHSN268201500014C |
| phs000988 | NHLBI TOPMed: The Genetic Epidemiology of Asthma in Costa Rica | UW NWGC | 3R37HL066289-13S1 |
| phs001412 | NHLBI TOPMed: Diabetes Heart Study African American Coronary Artery Calcification (AA CAC) | BROAD | HHSN268201500014C |
| phs000946 | NHLBI TOPMed: Boston Early-Onset COPD Study in the TOPMed Program | UW NWGC | 3R01HL089856-08S1 |
| phs000974 | NHLBI TOPMed: Whole Genome Sequencing and Related Phenotypes in the Framingham Heart Study | BROAD | 3R01HL092577-06S1 |
| phs000920 | NHLBI TOPMed: Genes-environments and Admixture in Latino Asthmatics (GALA II) Study | NYGC | 3R01HL117004-02S3 |
| phs001218 | NHLBI TOPMed: GeneSTAR (Genetic Study of Atherosclerosis Risk) | MACROGEN, BROAD | HHSN268201500014C |
| phs001345 | NHLBI TOPMed: Genetic Epidemiology Network of Arteriopathy (GENOA) | BROAD, UW NWGC | HHSN268201500014C, 3R01HL055673-18S1 |
| phs001217 | NHLBI TOPMed: Genetic Epidemiology Network of Salt Sensitivity (GenSalt) | BAYLOR | HHSN268201500015C |
| phs001359 | NHLBI TOPMed: Genetics of Lipid Lowering Drugs and Diet Network (GOLDN) | UW NWGC | 3R01HL104135-04S1 |

| | | | |
|---|---|---|---|
| phs000993 | NHLBI TOPMed: Heart and Vascular Health Study (HVH) | BROAD, BAYLOR | 3R01HL092577-06S1, 3U54HG003273-12S2 |
| phs001293 | NHLBI TOPMed: HyperGEN - Genetics of Left Ventricular (LV) Hypertrophy | UW NWGC | 3R01HL055673-18S1 |
| phs000964 | NHLBI TOPMed: The Jackson Heart Study | UW NWGC | HHSN268201100037C |
| phs001402 | NHLBI TOPMed: Whole Genome Sequencing of Venous Thromboembolism (WGS of VTE) | BAYLOR | HHSN268201500015C, 3U54HG003273-12S2 |
| phs001416 | NHLBI TOPMed: MESA and MESA Family AA-CAC | BROAD | 3U54HG003067-13S1, HHSN268201500014C |
| phs001062 | NHLBI TOPMed: MGH Atrial Fibrillation Study | BROAD | 3R01HL092577-06S1 |
| phs001024 | NHLBI TOPMed: Partners HealthCare Biobank | BROAD | 3R01HL092577-06S1 |
| phs001215 | NHLBI TOPMed: San Antonio Family Heart Study (WGS) | ILLUMINA | 3R01HL113323-03S1 |
| phs000921 | NHLBI TOPMed: Study of African Americans, Asthma, Genes and Environment (SAGE) Study | NYGC | 3R01HL117004-02S3 |
| phs001207 | NHLBI TOPMed: African American Sarcoidosis Genetics Resource | BAYLOR | 3R01HL113326-04S1 |
| phs000972 | NHLBI TOPMed: Genome-wide Association Study of Adiposity in Samoans | UW NWGC, NYGC | HHSN268201100037C, HHSN268201500016C |
| phs001387 | NHLBI TOPMed: Rare Variants for Hypertension in Taiwan Chinese (THRV) | BAYLOR | 3R01HL111249-04S1, HHSN26820150015C |
| phs000997 | NHLBI TOPMed: The Vanderbilt AF Ablation Registry | BROAD | 3R01HL092577-06S1 |
| phs001032 | NHLBI TOPMed: The Vanderbilt Atrial Fibrillation Registry | BROAD | 3R01HL092577-06S1 |
| phs001040 | NHLBI TOPMed: Novel Risk Factors for the Development of Atrial Fibrillation in Women | BROAD | 3R01HL092577-06S1 |
| phs001237 | NHLBI TOPMed: Women's Health Initiative (WHI) | BROAD | HHSN268201500014C |

**Supplementary Table 19. Number of variants in each sequencing set in Framingham Heart Study.**

| Sequencing Set | Total | SNVs | Indels | Multi-allelic |
|---|---|---|---|---|
| TOPMed WGS Depth>0 (4,158) | 58,740,718 | 51,390,7555 | 4,519,101 | 2,830,862 |
| TOPMed WGS Depth>0 (430) | 23,836,812 | 21,120,031 | 1,697,530 | 1,019,251 |
| CHARGE WGS (855) | 25,832,397 | 25,832,397 | N/A | N/A |
| CHARGE WGS (430) | 20,546,566 | 20,546,566 | N/A | N/A |
| CHARGE WES (1,701) | 475,758 | 441,263 | 11,533 | 22,962 |
| CHARGE WES (430) | 225,455 | 210,025 | 4,629 | 10,801 |

**Supplementary Table 20. Average bi-allelic SNV Rates.**

| Study | All SNVs | Exonic SNVs |
|---|---|---|
| TOPMed | 0.138 (2,383,558.6) | 0.095 (15,829.1) |
| CHARGE WGS | 0.151 (2,658,391.9) | 0.115 (17,374.4) |

**Supplementary Table 21. Counts and percentages of variants by minor allele frequency.**

| MAF | TOPMed (%) | CHARGE WGS (%) | CHARGE WES (%) |
|---|---|---|---|
| 0.001-0.005 | 8,900.787 (47) | 6,724,501 (38) | 105,781 (60) |
| 0.005-0.01 | 1,527,058 (8) | 1,605,014 (9) | 15,502 (9) |
| 0.01-0.05 | 2,482,218 (13) | 2,920,460 (17) | 21,457 (12) |
| 0.05-0.1 | 1,178,543 (6) | 1,469,328 (8) | 8,414 (5) |
| 0.1-0.2 | 1,542,854 (8) | 1,695,692 (10) | 9,503 (5) |
| 0.2-0.3 | 1,139,687 (6) | 1,187,933 (7) | 6,277 (3) |
| 0.3-0.4 | 1,017,644 (5) | 1,048,952 (6) | 5,301 (3) |
| 0.4-0.5 | 944,966 (5) | 970,532 (5) | 4,709 (3) |

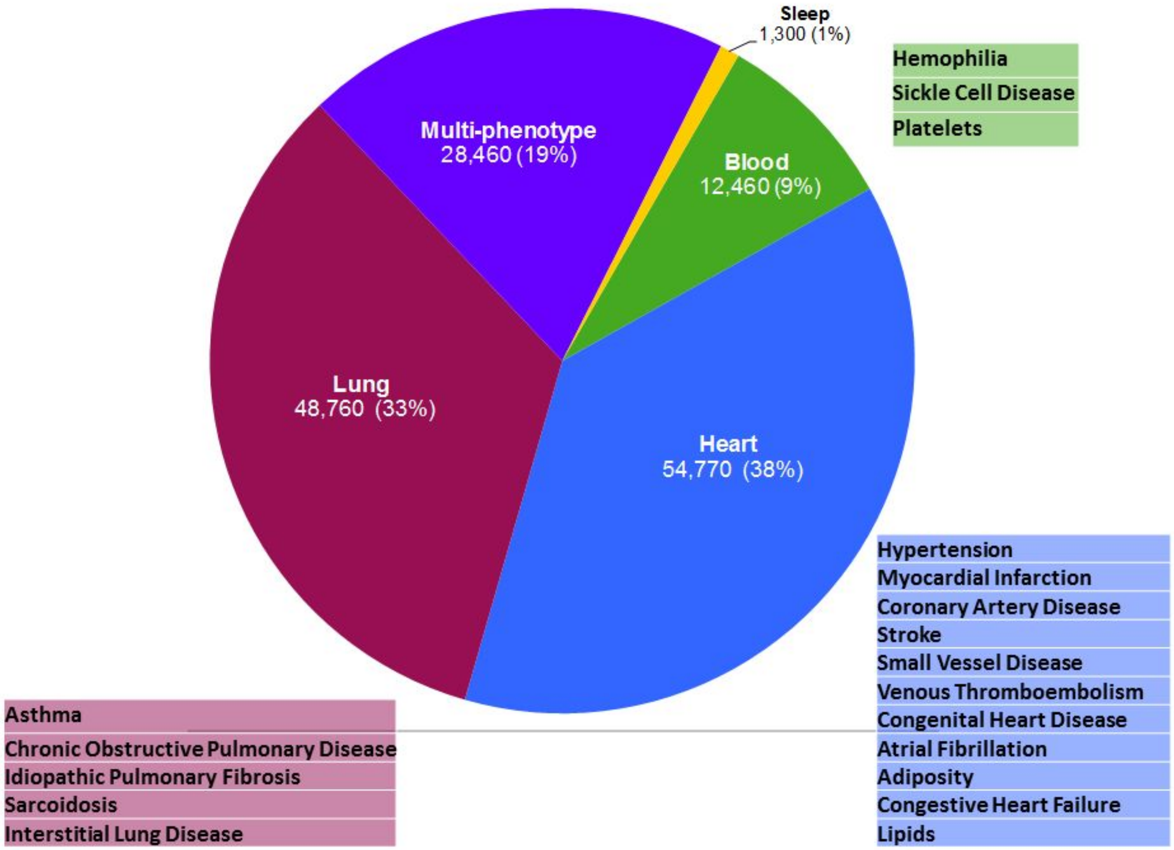


**Supplementary Figure 1. Areas of phenotype focus among TOPMed Parent studies.** Approximate sample sizes and percentages are given for ~145,000 participants in the first four Phases of the program who have been or are being whole-genome sequenced. "Multi-phenotype" refers to cohort studies with a wide range of phenotypes. In addition, lung studies also have many heart-related phenotypes and vice versa. The participant numbers given do not refer specifically to case counts, but rather to all participants in the study.

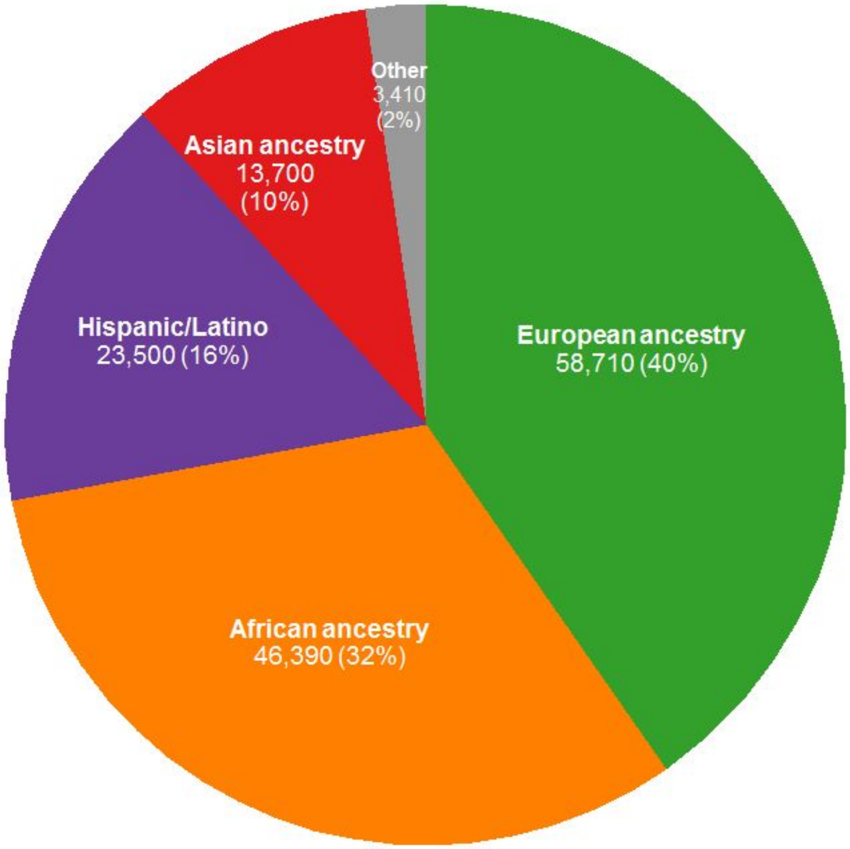


**Supplementary Figure 2. Major ancestral and ethnic groups among TOPMed participants.** Approximate sample sizes and percentages are given for ~145,000 participants in the first four Phases of the program who have been or are being whole-genome sequenced. These groups consist of European (European, European-American); African ancestry (African, African-American, African-Caribbean); Hispanic/Latino (including Mexican, Mexican American, Central American, South American, Cuban, Dominican, Puerto Rican); Asian ancestry (Chinese, Taiwanese, Asian American, Pakistani); and ‘Other’.

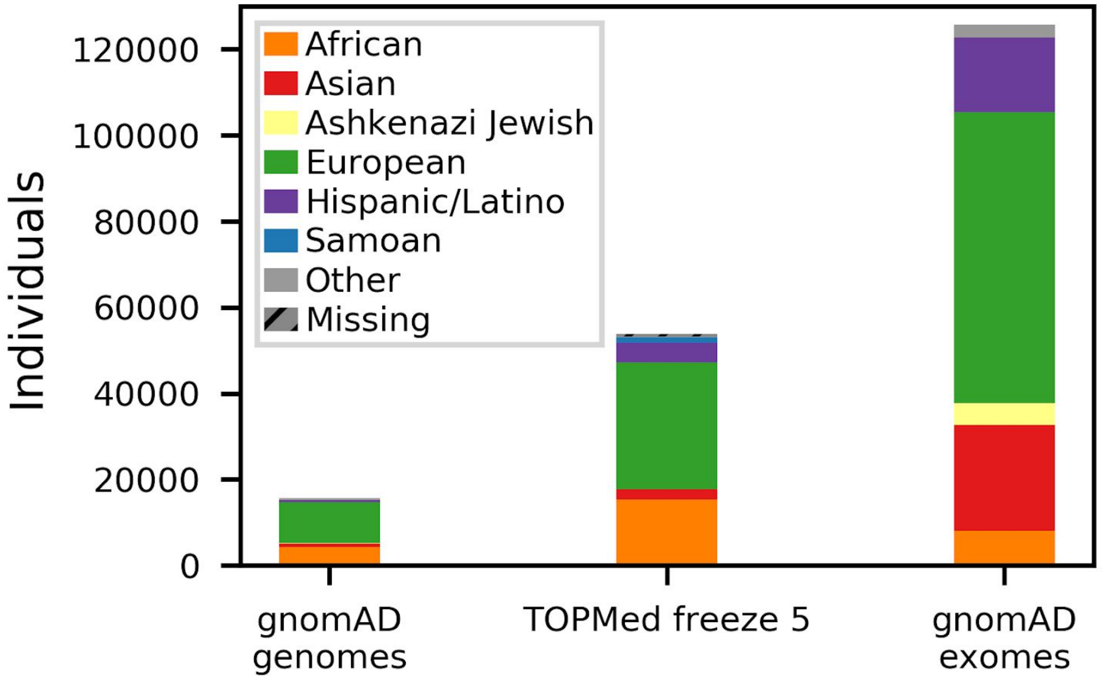


**Supplementary Figure 3. Ancestral diversity among 53,831 participants in TOPMed Freeze 5 compared to other largest public data sets.** TOPMed freeze 5 categories are based on self-identification.

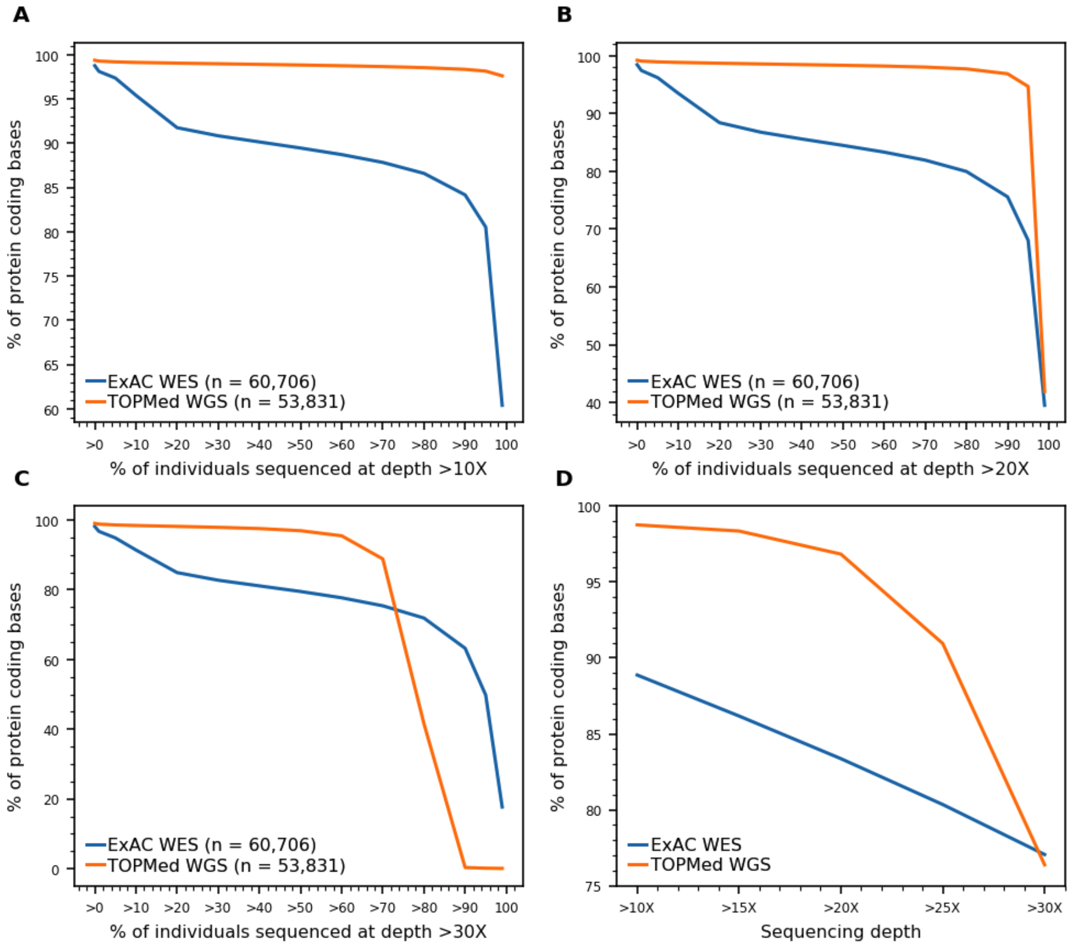


**Supplementary Figure 4. Sequencing depth at protein coding bases in TOPMed and ExAC.** We computed percent of protein coding bases in CCDS genes sequenced at different depth in TOPMed and ExAC datasets. A. Percent of protein coding bases which where sequenced at depth >10X. B. Percent of protein coding bases which where sequenced at depth >20X. C. Percent of protein coding bases which where sequenced at depth >30X. D. Percent of protein coding bases across all individuals sequenced at different depth.

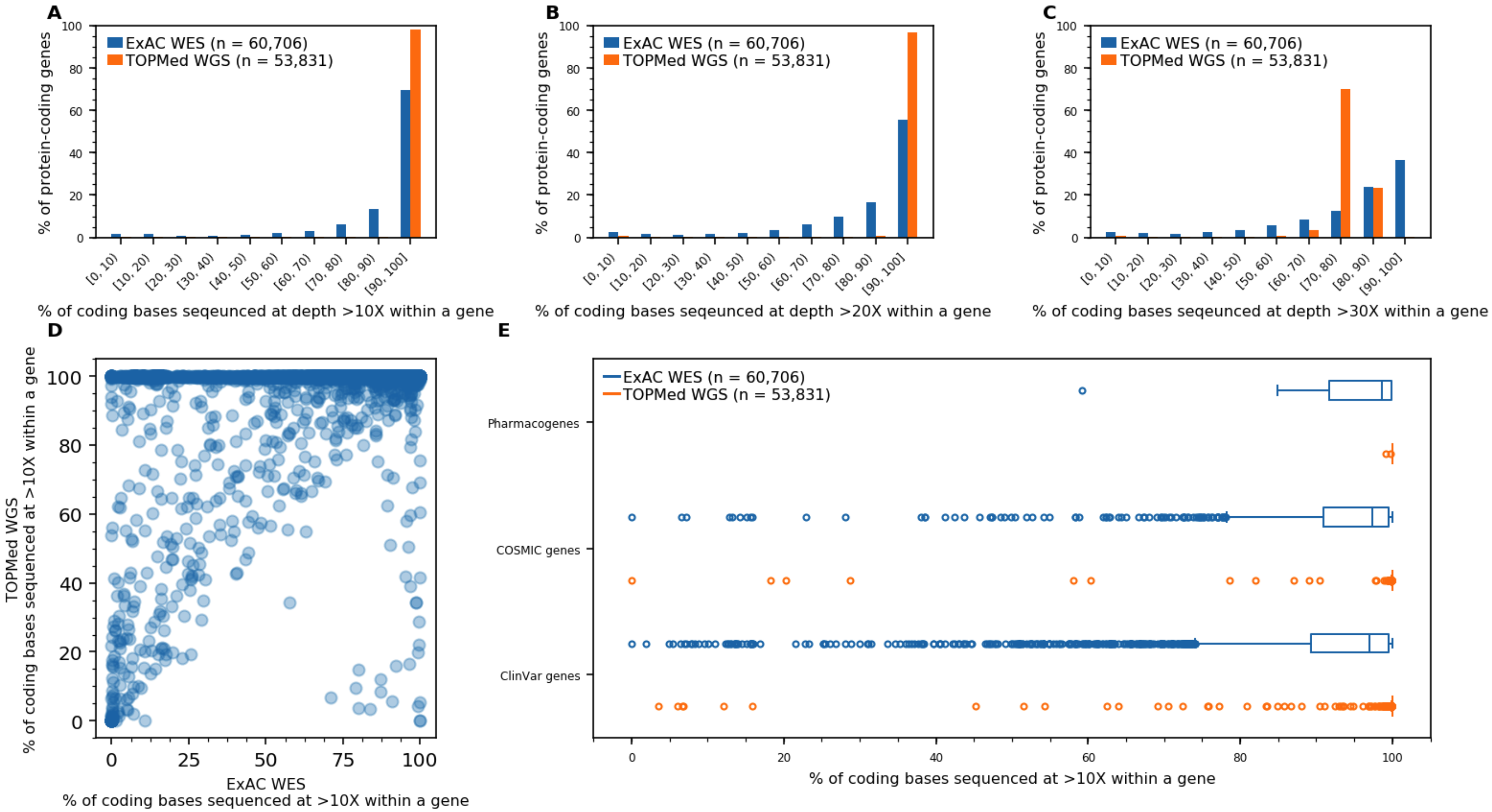


**Supplementary Figure 5. Sequencing depth at protein coding genes in TOPMed and ExAC.** We compared sequencing depth at protein coding genes, present in Consensus Coding Sequence (CCDS) Project, in ExAC and TOPMed. A-C. Percent of protein-coding CCDS genes in ExAC and TOPMed with different percent of coding bases sequenced at >10X, >20X, and >30X, respectively. D. For each protein-coding CCDS gene (point) shows percent of coding bases sequenced at >10X in ExAC (x axis) and TOPMed (y axis). E. Percent of coding bases sequenced at >10X in ExAC and TOPMed in genes from ClinVar database (at least one pathogenic variant reported), genes from COSMIC database, and pharmacogenes.

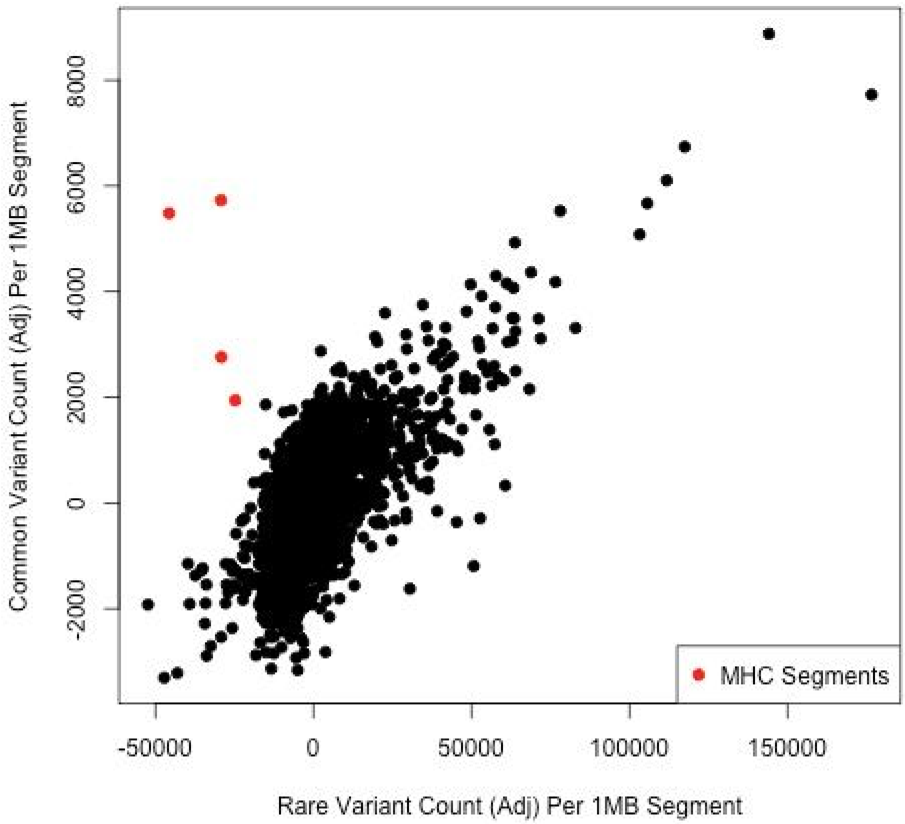


**Supplementary Figure 6. Scatter plot of common and rare variant counts per contiguous segment**. After using regression to adjust variant counts by the proportion of bases per 1Mb segment that was flagged for mappability concerns, the number of common (AF >= 0.5%) and rare variants (AF < 0.5%) in contiguous segments are highly and significantly correlated ($R^2$ = 0.462, p-value ≤ 2e-16). Outlier segments with higher than expected levels of common variation overlap MCH regions of the genome (red), which is consistent with the effects of balancing selection known to shape these loci.

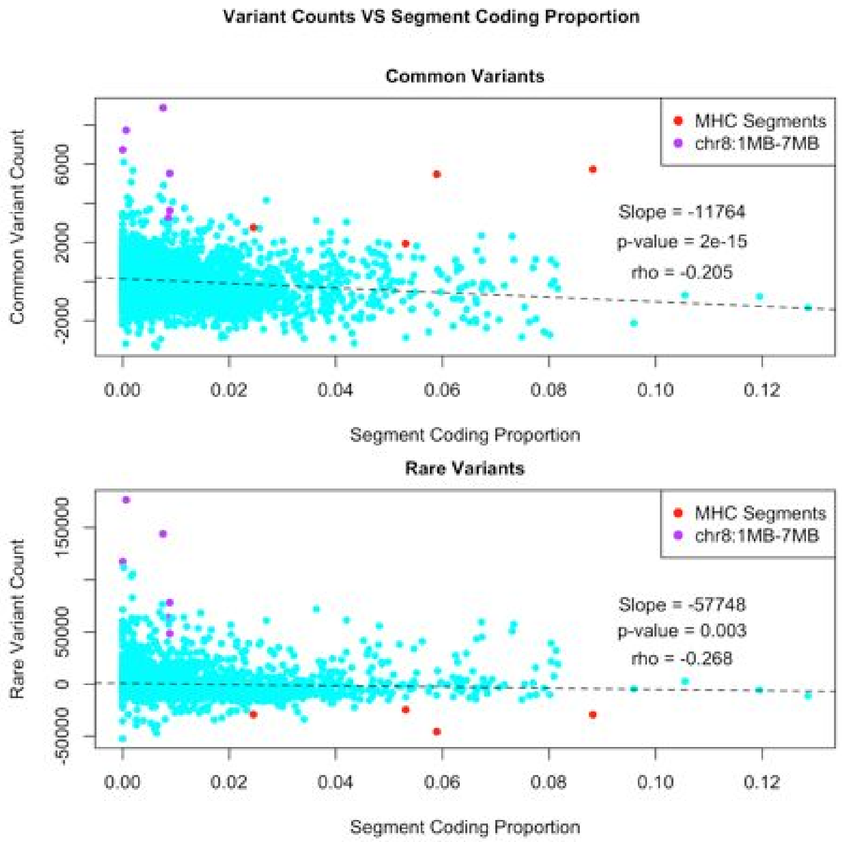


**Supplementary Figure 7. Scatter plot of the relationship between variant count and contiguous segment coding proportion.** There is a significant negative correlation between segment coding proportion and variant count, which holds when subsetting variants according to allele frequency (i.e. common and rare variants). Outliers, representing regions of potential interest can be seen, including megabases 1 to 7 on chromosome 8 (purple) and segments overlapping Major Histocompatibility Complex (MHC) genes (red). Counts are adjusted for segment coding proportion and mappability (i.e. accessibility mask, see Supplementary Section 1.2).

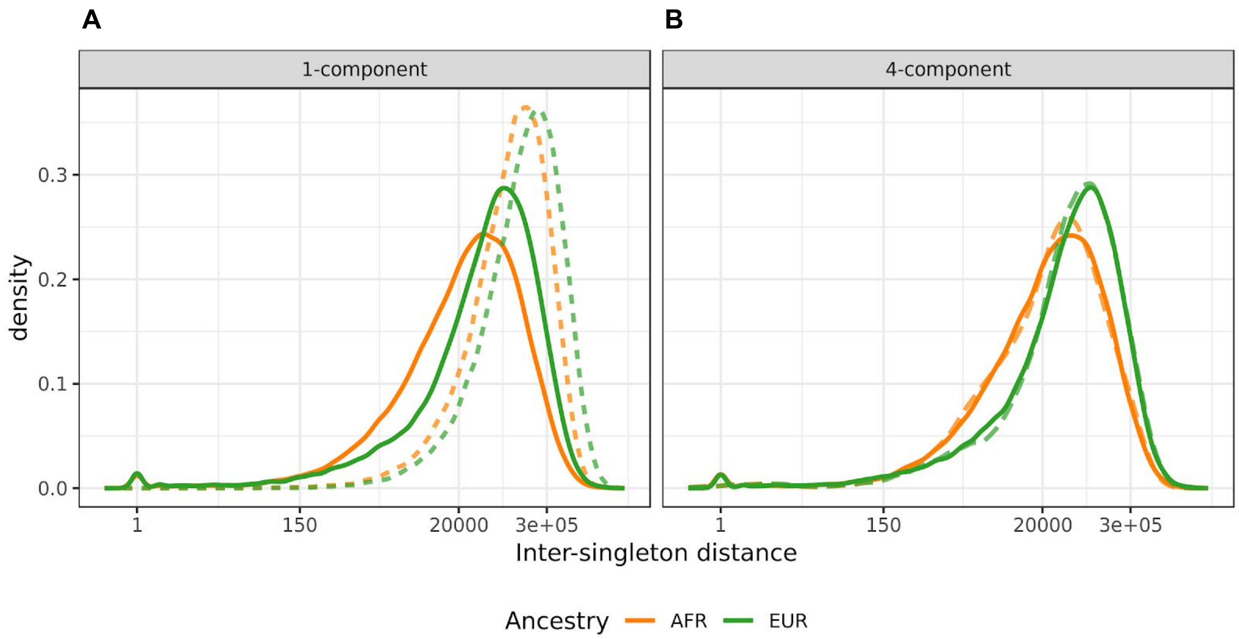


**Supplementary Figure 8. Comparison of observed (solid) and expected (dashed) inter-singleton distance distributions. A.** Comparison of observed inter-singleton distance distributions with the expected inter-singleton distance distributions under a simple exponential model, where we assume singletons occur through a single uniform Poisson process. The rate parameters for the expected distributions are based on the median number of singletons per individual, as observed in the European and African ancestry subsamples ($\theta_{AFR} = 6.3\times 10^{-6}$; $\theta_{EUR} = 4.5\times 10^{-6}$). **B.** Comparison of observed inter-singleton distance distributions with the inter-singleton distance distributions expected using the fitted 4-component mixture models.

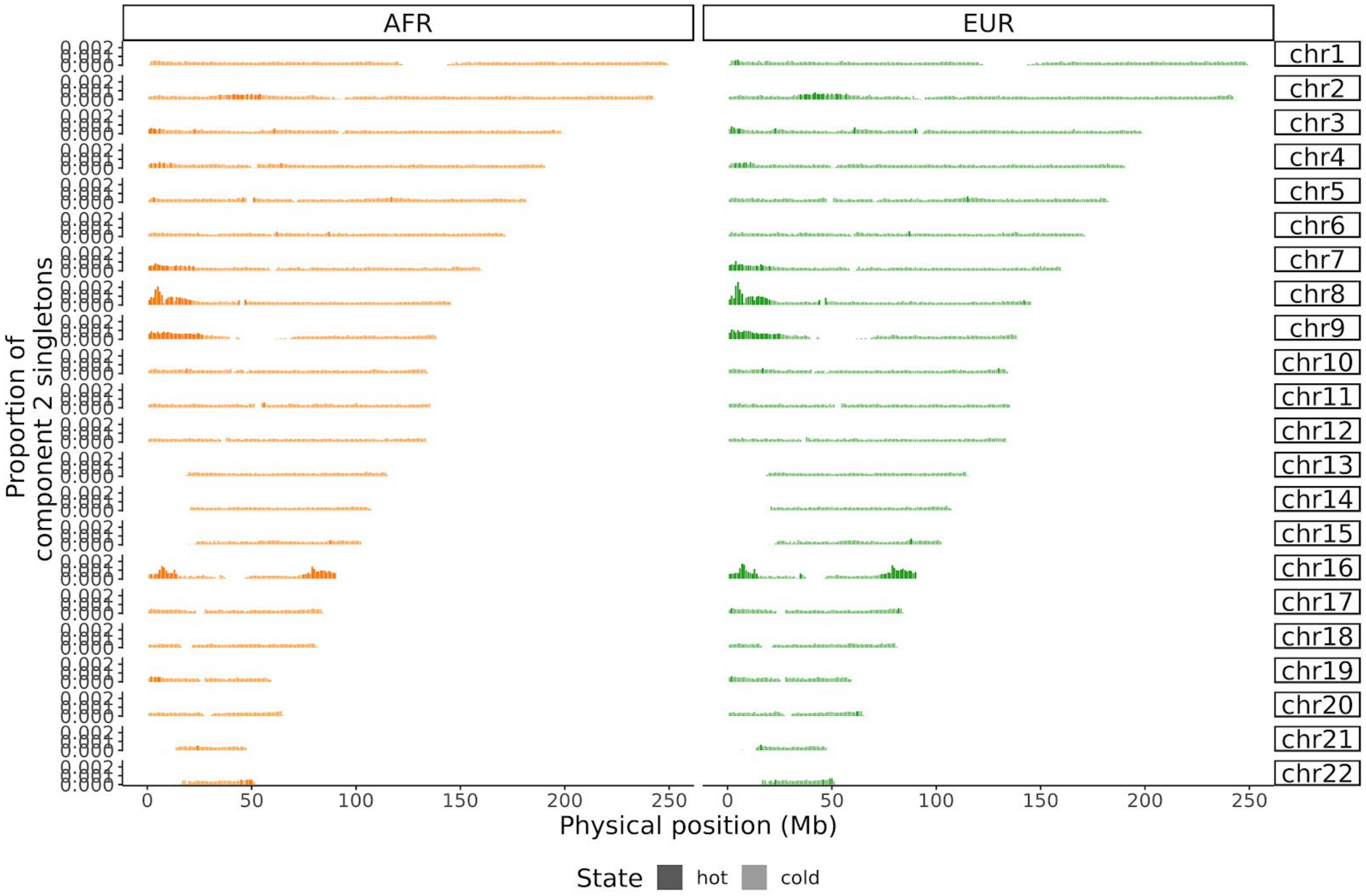


**Supplementary Figure 9. Genomic hotspots for cluster class 2 singletons.** Density of cluster class 2 singletons in 1Mbp windows throughout the autosomal chromosomes. Windows with class 2 singleton counts above the 95th percentile (calculated genome-wide) are classified as hotspots and are represented with a darker shade (as in Figure 2C).

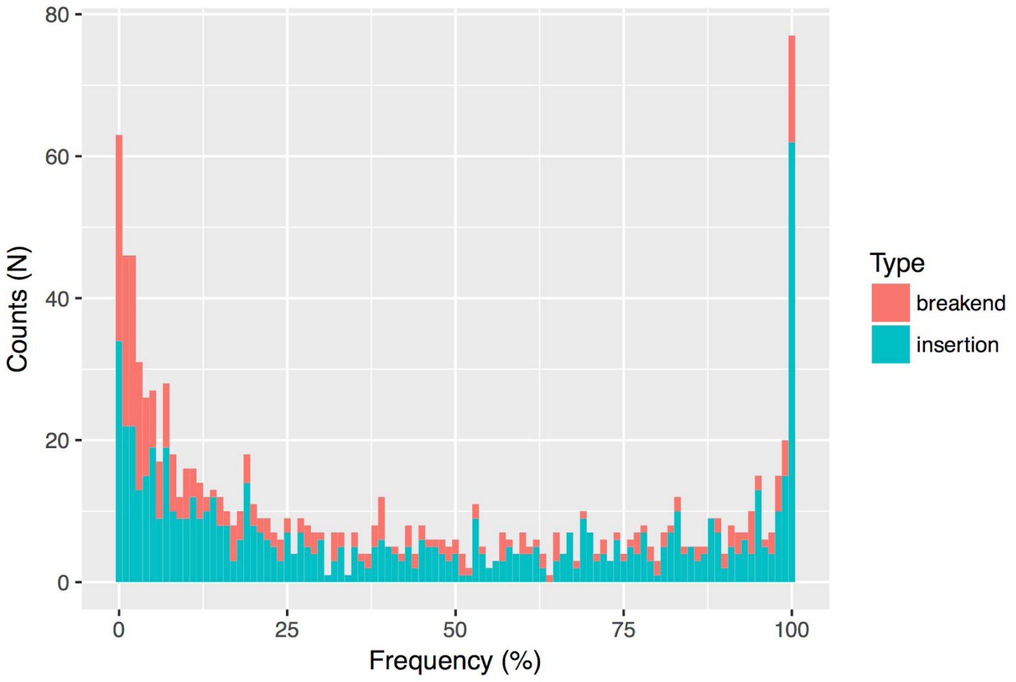


**Supplementary Figure 10.** Insertion and breakend frequency distribution on a subset of the samples (N = 17,443).

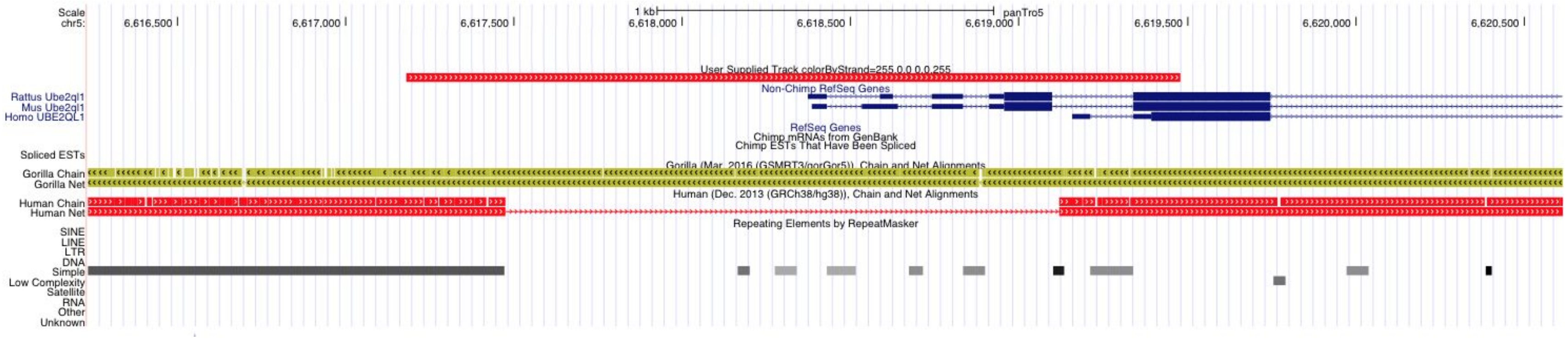


**Supplementary Figure 11.** UCSC Genome Browser screenshot of a non-reference human contig aligned to the chimp genome (panTro5), overlapping both the transcription and translation start sites of *UBE2QL1* gene from mouse and rat that been aligned to the same reference.

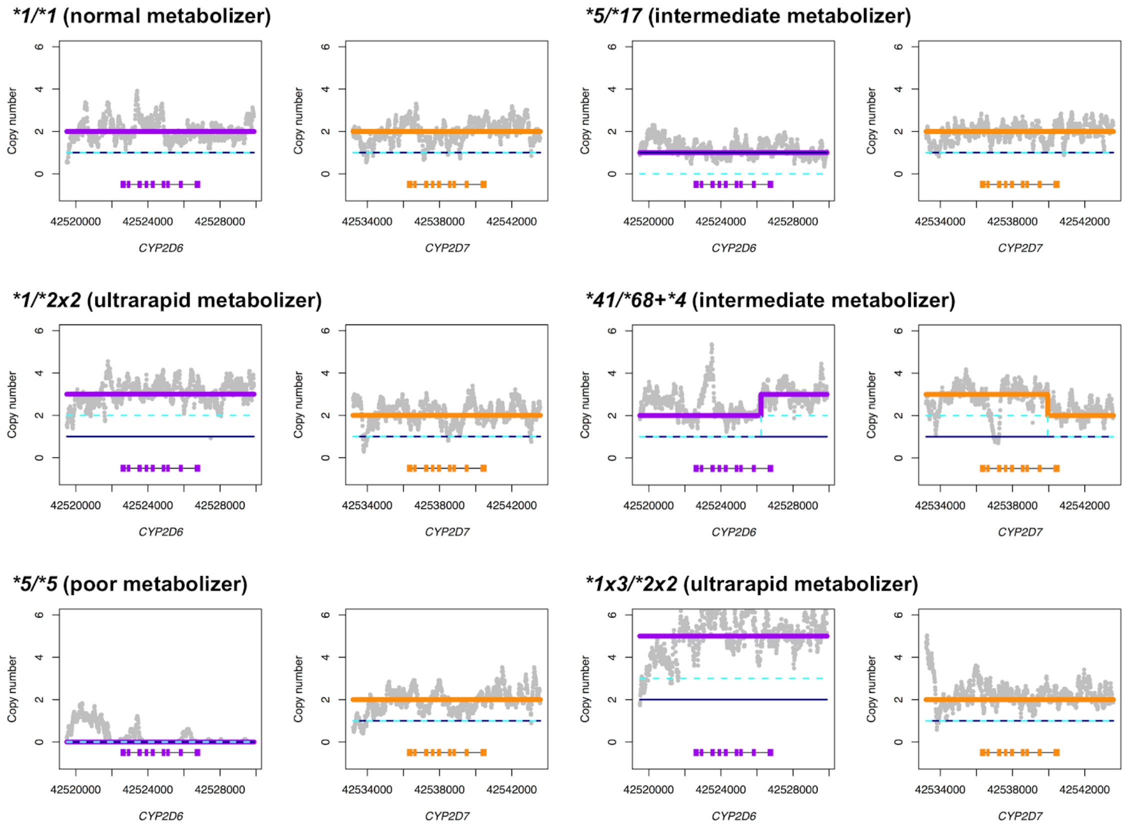


**Supplementary Figure 12. Stargazer's copy number plots for *CYP2D6* and *CYP2D7* of six representative African American individuals from the Jackson Heart Study.** Grey dots show the copy number calculated from the read depth. The navy solid and cyan dashed lines represent expected copy number of two haplotypes that are being tested against the sample's observed copy number. The purple and orange lines represent expected copy number profiles for *CYP2D6* and *CYP2D7*, respectively, formed by combining the two haplotypes in question. Each panel contains the scaled *CYP2D6* and *CYP2D7* gene models, in which the exons and introns are depicted with boxes and lines, respectively.

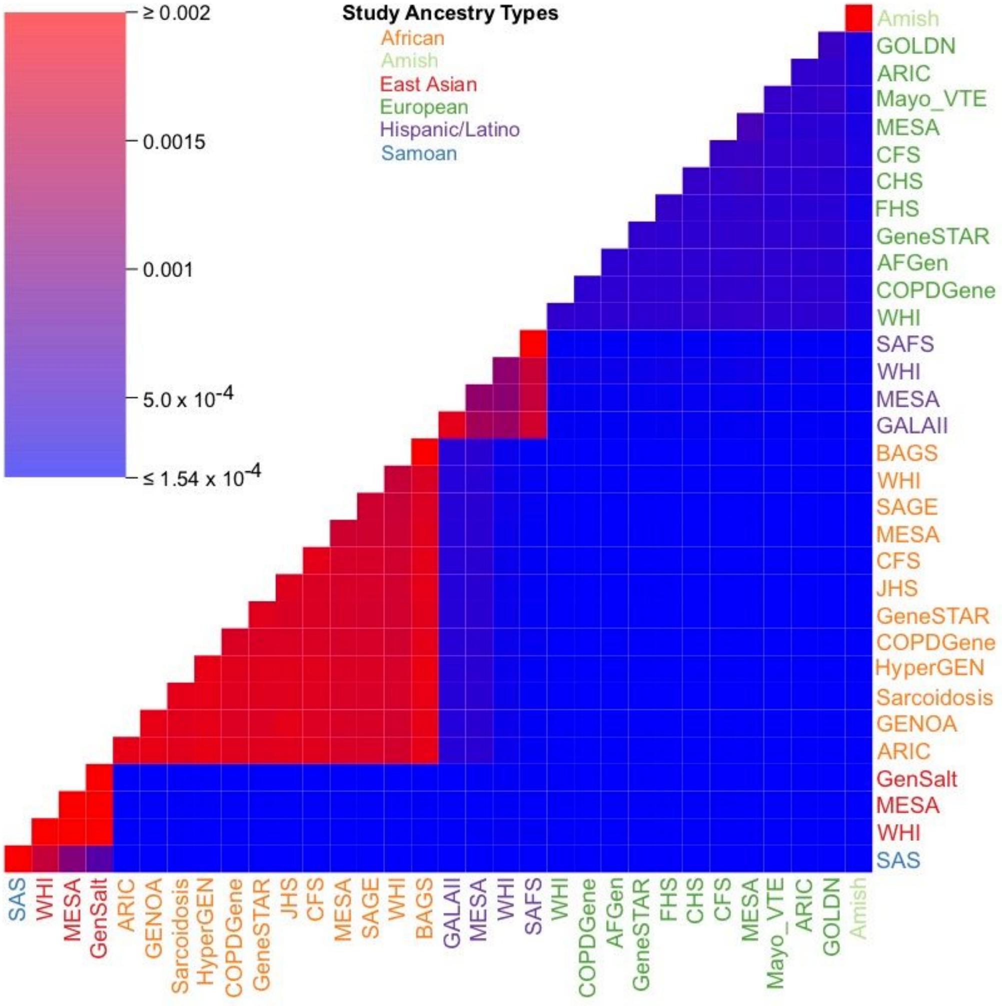


**Supplementary Figure 13.** Rare variant sharing between and within TOPMed studies. Each study label is colored based on self-described ancestry. The heatmap scale depicts the 50th percentile of between and within rare variant sharing values.

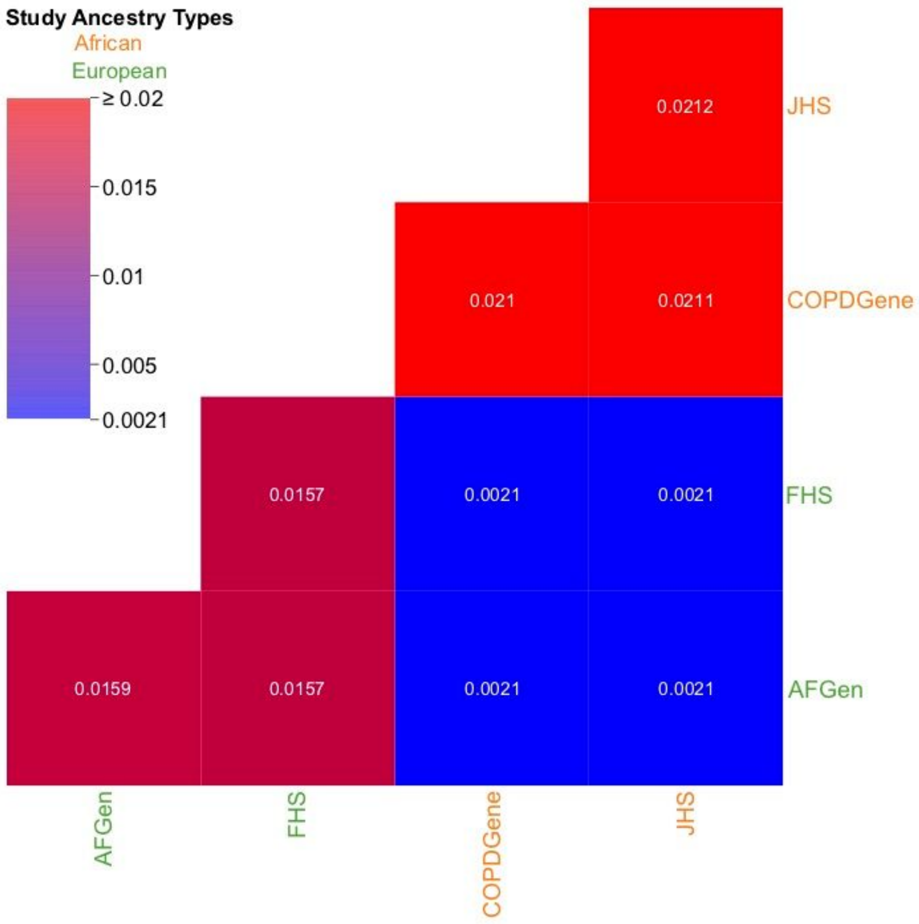


**Supplementary Figure 14. Equal Study Sizes Rare Variant Sharing Control.** Heatmap representation of sharing within and between 4 TOPMed studies each sampled to 500 individuals.

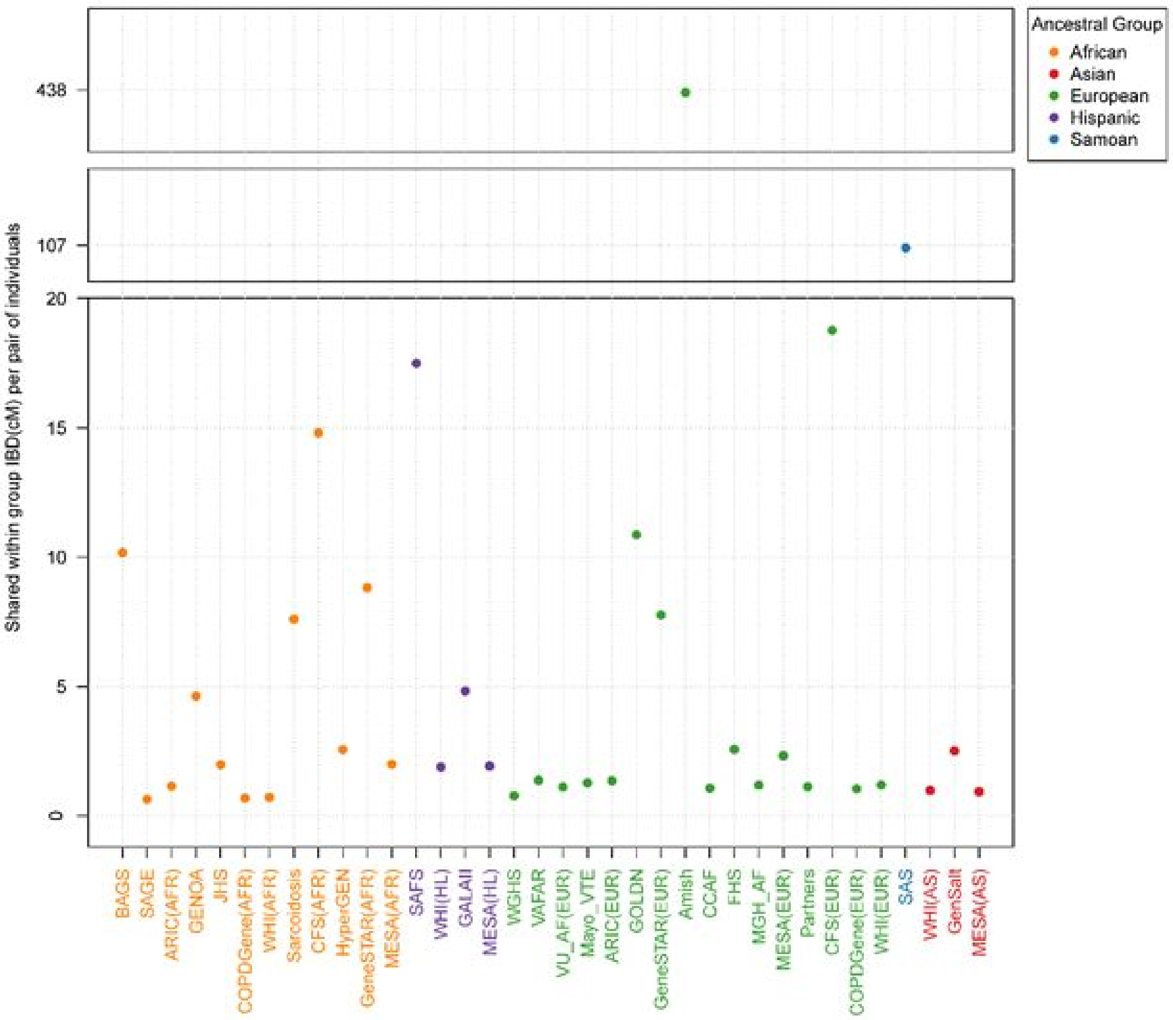


**Supplementary Figure 15. Within-group average IBD sharing.** We calculated the total autosome-wide length of detected IBD segments per pair of individuals and averaged across pairs within each group. For studies with multiple self-identified ancestries, parentheses after the study name identify the ancestry (AFR, African; HL, Hispanic/Latino; EUR, European; AS, Asian). Points and labels are colored by ancestry.

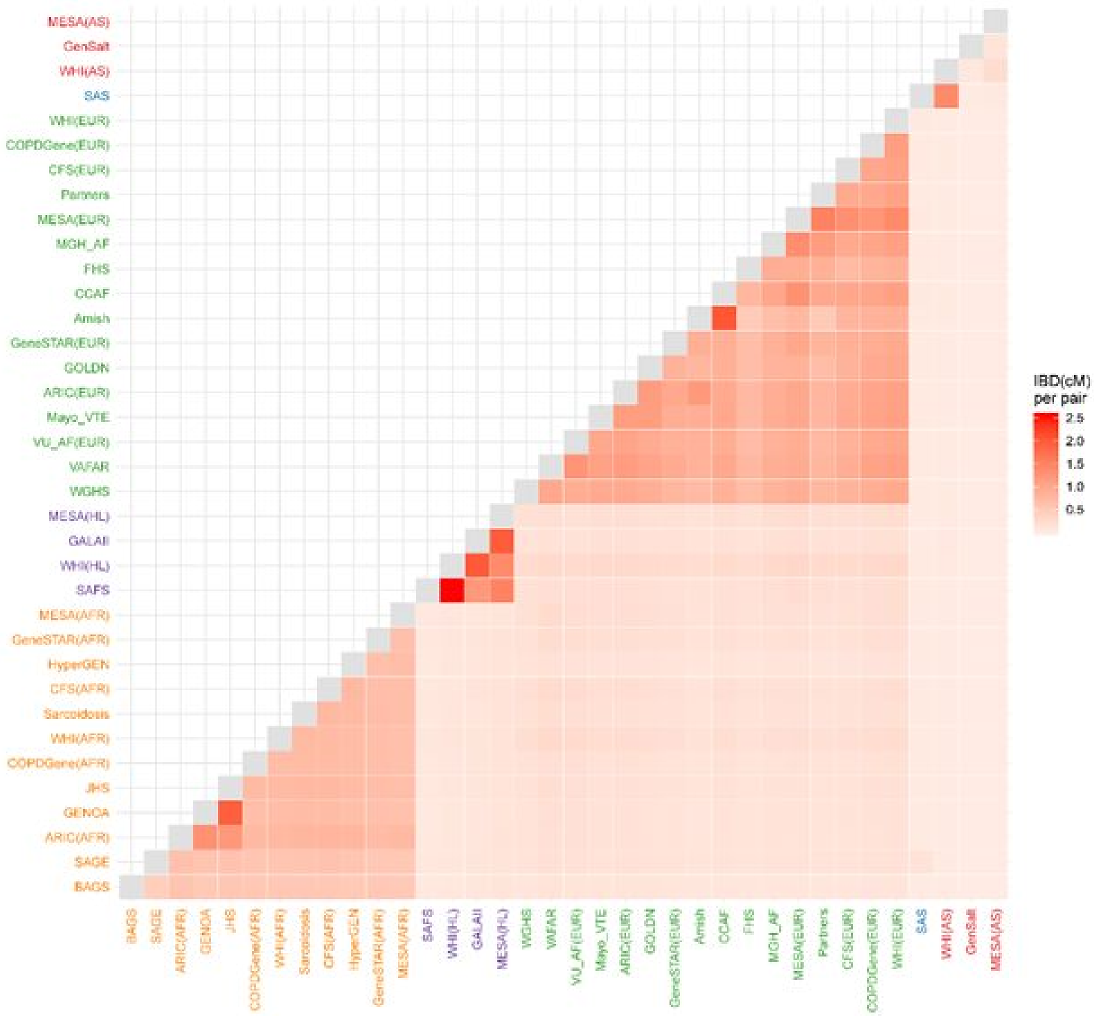


**Supplementary Figure 16. Between group average IBD sharing.** We calculated the total autosome-wide length of detected IBD segments per pair of individuals and averaged across pairs. For studies with multiple self-identified ancestries, parentheses after the study name identify the ancestry (AFR, African; HL, Hispanic/Latino; EUR, European; AS, Asian). Labels are colored by ancestry (Asian, red; Samoan, blue; European, green; Hispanic/Latino, purple; African, orange).

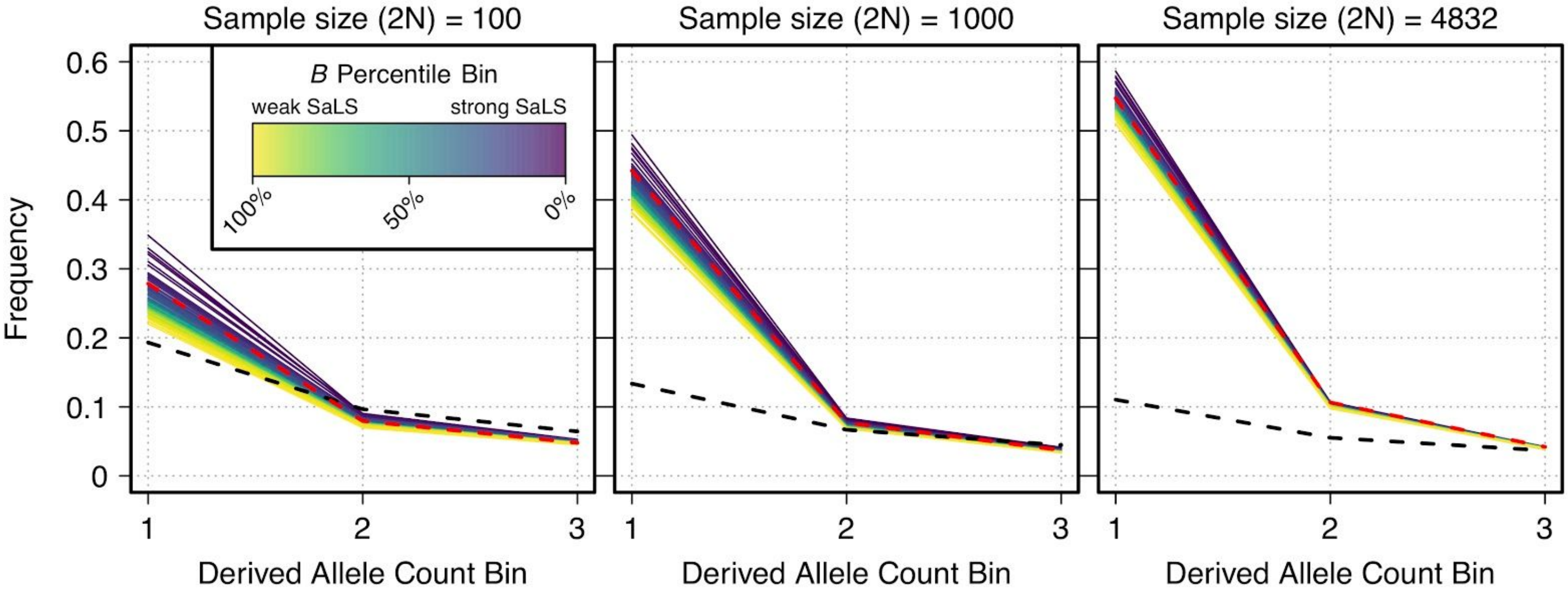


**Supplementary Figure 17. Site-frequency spectrum (SFS) for different sample sizes and *B* for the first three derived allele counts**. SFS data is shown for each of 100 percentile bins of *B* (McVicker's *B* statistic; higher percentiles of *B* indicate weaker effects of selection at linked sites [SaLS]). Each separate plot shows a different sample size from which the SFS was made. Dashed red lines show the SFS from fourfold degenerate sites. Dashed black lines show the SFS from a standard neutral model for the given sample size.

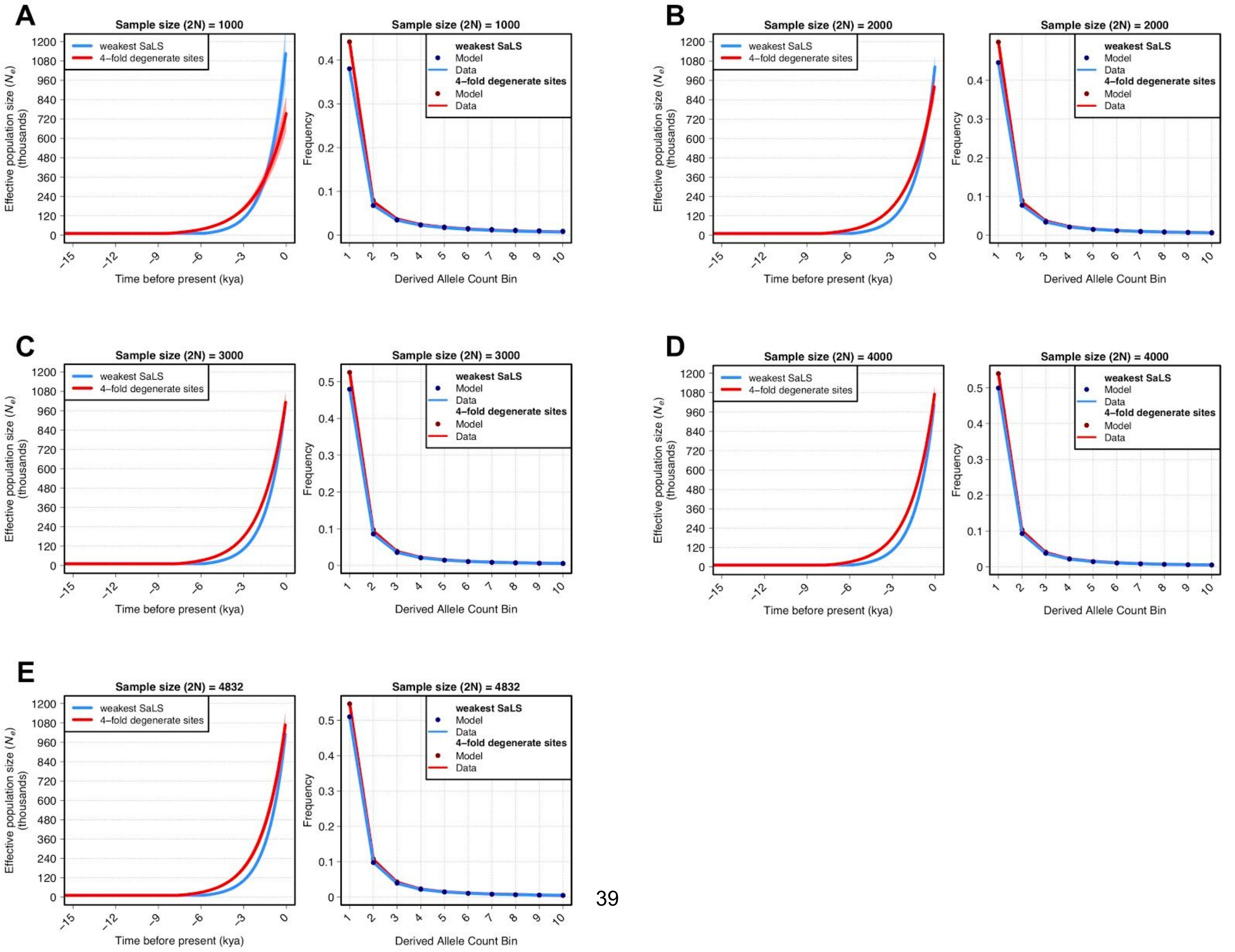
A
B
C
D
E
Sample size (2N) = 1000
Sample size (2N) = 2000
Sample size (2N) = 3000
Sample size (2N) = 4000
Sample size (2N) = 4832
weakest SaLS
4–fold degenerate sites
Model
Data
Effective population size ($N_e$) (thousands)
Time before present (kya)
Frequency
Derived Allele Count Bin

**Supplementary Figure 18. Results from performing demographic inference fitting a model of exponential growth to fourfold degenerate sites and sites under the weakest effects of selection at linked sites (SaLS).** Weakest SaLS represent sites from the highest 1% *B* bin (McVicker's *B* statistic). The left panels of each figure show the inferred exponential growth using various sample sizes. Shaded envelopes represent 95% confidence intervals (see Supplementary Table 9 for parameter values). The right panels of each figure show the observed site-frequency spectrum represented as solid lines. The resulting fits to the site-frequency spectrum from the fitted demographic models are shown as points.

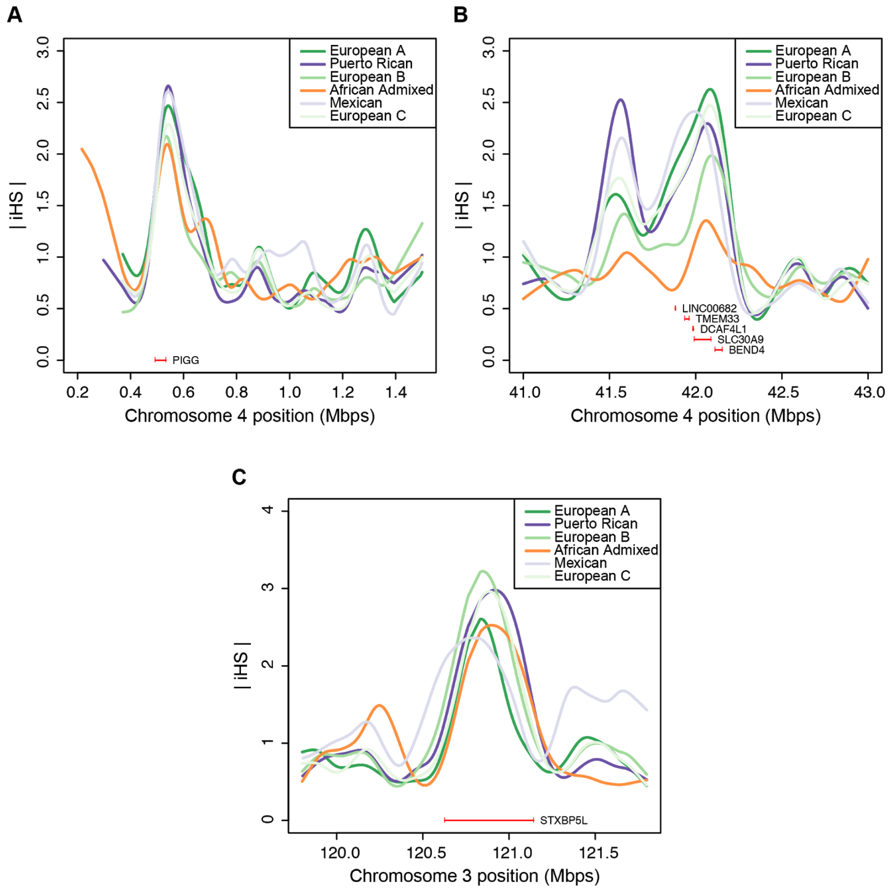


**Supplementary Figure 19. Smoothed plot of |iHS| scores in each population in a region identified in each population as having strong evidence for positive selection. A.** Region around protein coding gene *PIGG*. **B.** Region around protein coding gene *SLC30A9*. **C.** Region around protein coding gene *STXBP5L.*

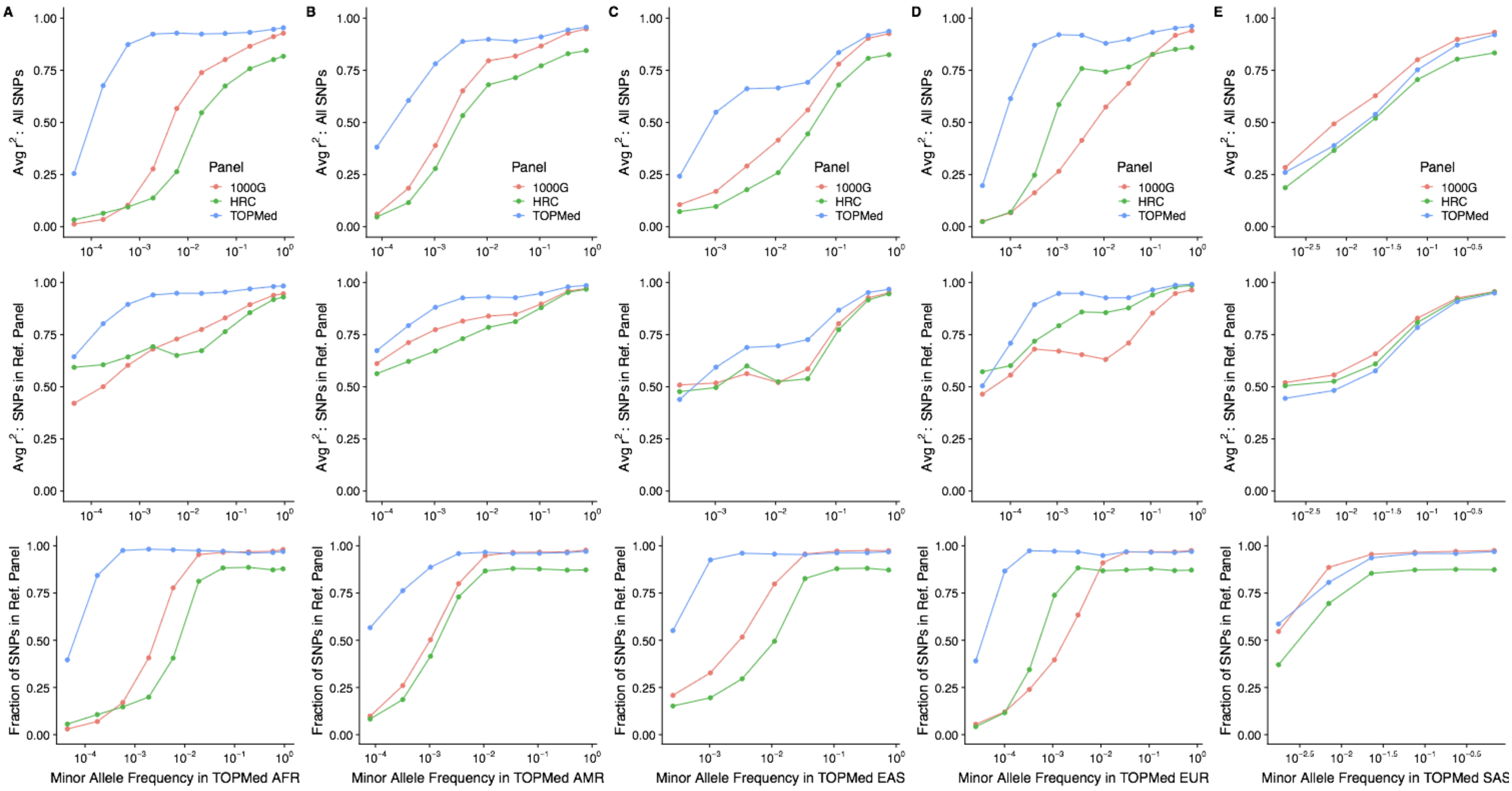


**Supplementary Figure 20. Evaluation of imputation accuracy.** Evaluation of genotype imputation accuracy from various reference panels - 1000 Genomes Phase 3 Panel (1000G), Haplotype Reference Consortium Panel (HRC), and TOPMed Panel (TOPMed). Each column represents a different continental population matched with five 1000 Genomes continental populations, namely (A) AFR : Africans, (B) AMR : Admixed Americans, (C) EAS : East Asians, (D) EUR : Europeans, (E) SAS : South Asians. 100 samples from the BioMe study that were not included in the imputation panel are selected from each continental population, and population-specific allele frequencies are calculated excluding the selected target samples. Top panels show the average squared correlation (r2) between the sequence-based genotypes and imputed dosages across all variants, assigning r2 = 0 to variants absent from each Reference Panel. The middle panels compute average r2 with only the variants present from each Reference Panel. The proportion of variants present in the reference panels is shown in the bottom panel.

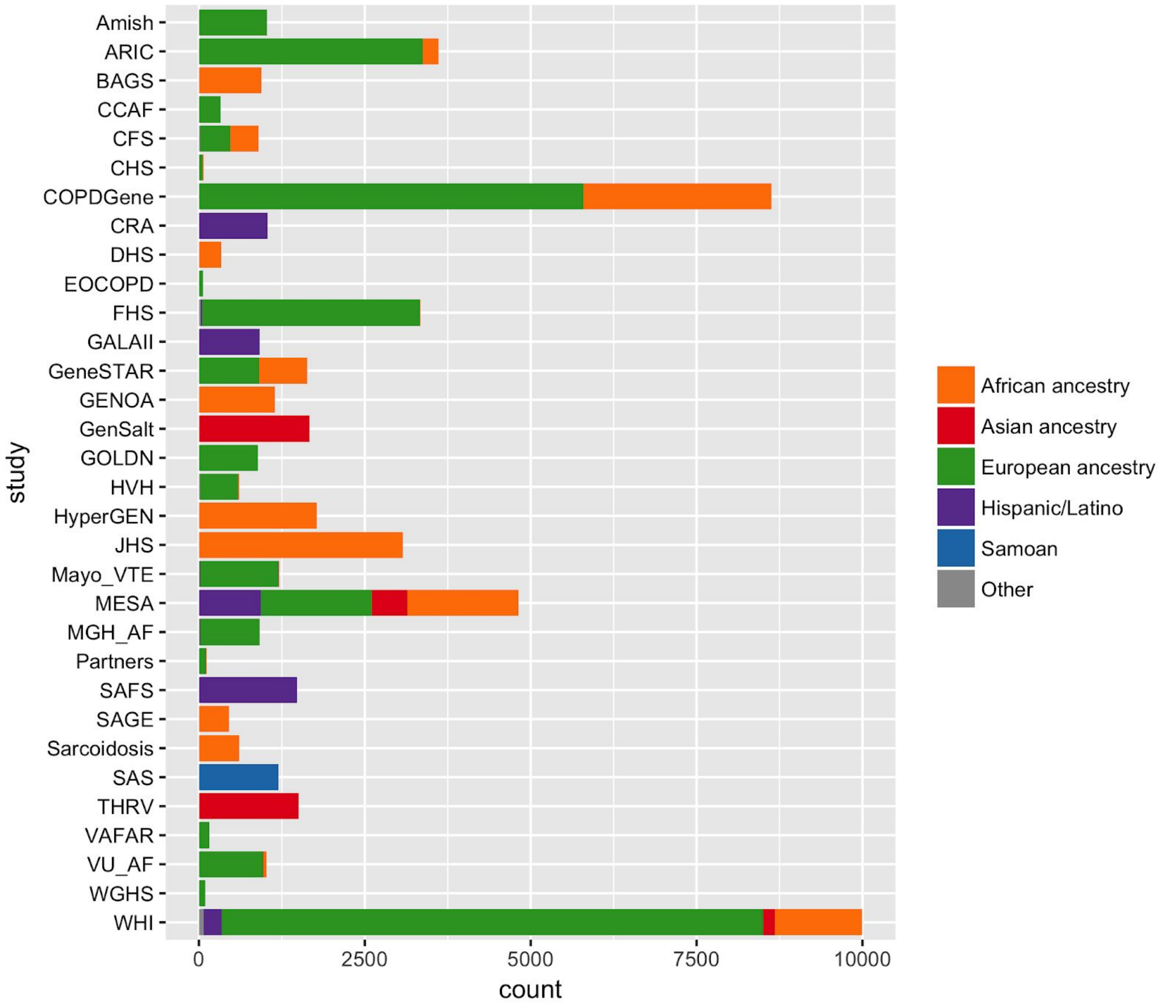


**Supplementary Figure 21. Ancestral/ethnic composition of studies included in the TOPMed Freeze 5 genotype call set.** These counts are based on participant responses to questions regarding race and ethnicity and/or study recruitment criteria. See Supplementary Table 2 for study abbreviations.

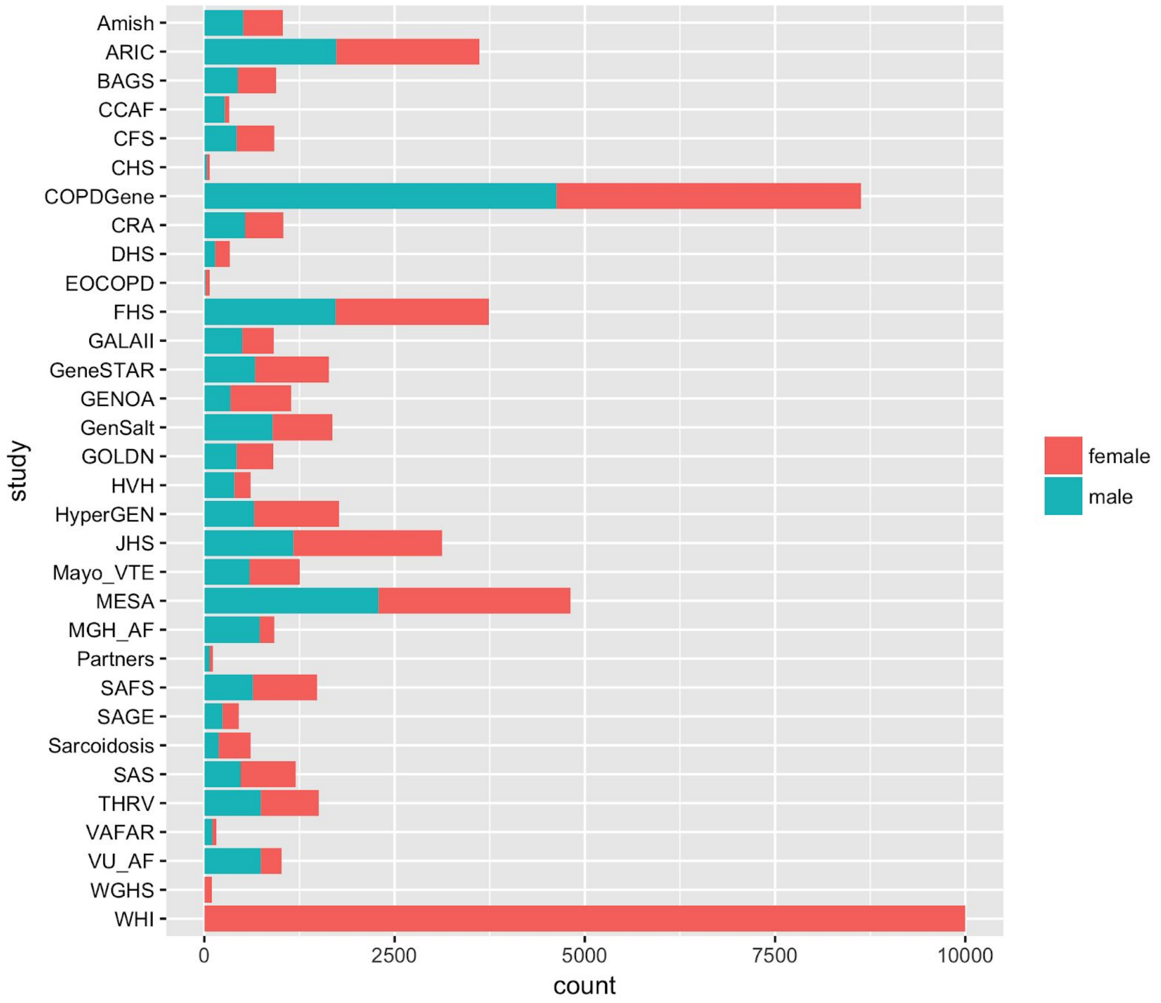


**Supplementary Figure 22. Sex composition of studies included in the TOPMed Freeze 5 genotype call set**. See Supplementary Table 2 for study abbreviations.

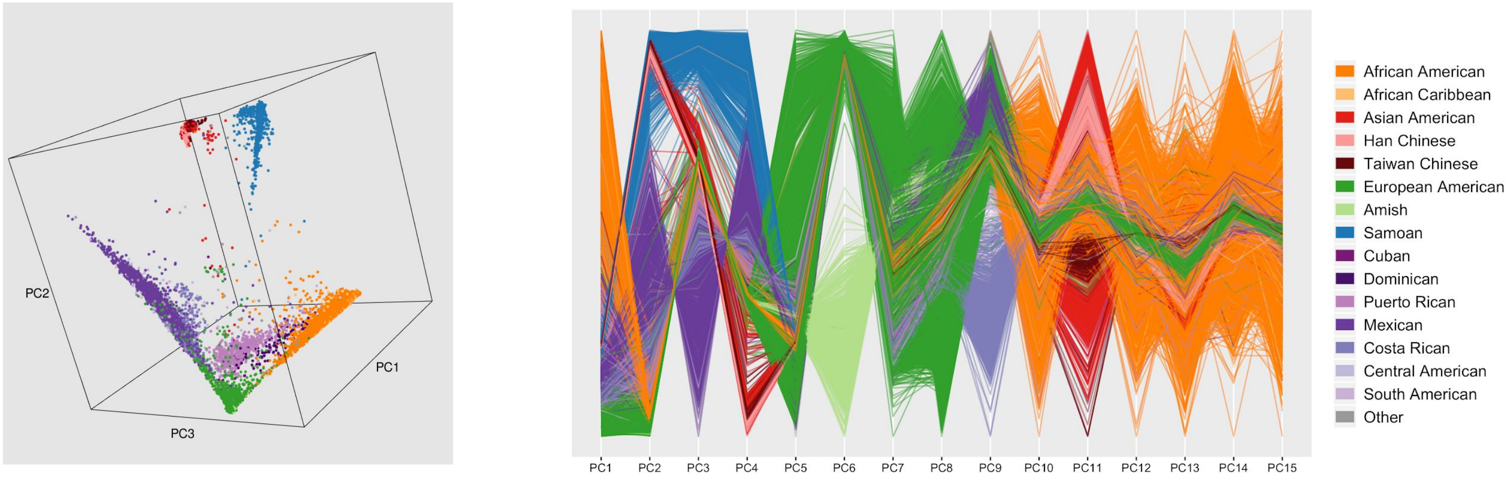


**Supplementary Figure 23. Principal components of the genotypic data from Freeze 5 pooled over studies.** The left panel is a three-dimensional plot of principal components 1, 2 and 3. The right panel is a parallel coordinates plot color-coded by categories defined according to race, ancestry and/or ethnic information provided by the study participants and/or by study investigators according to study inclusion criteria. Subjects with missing values for ancestry are excluded.

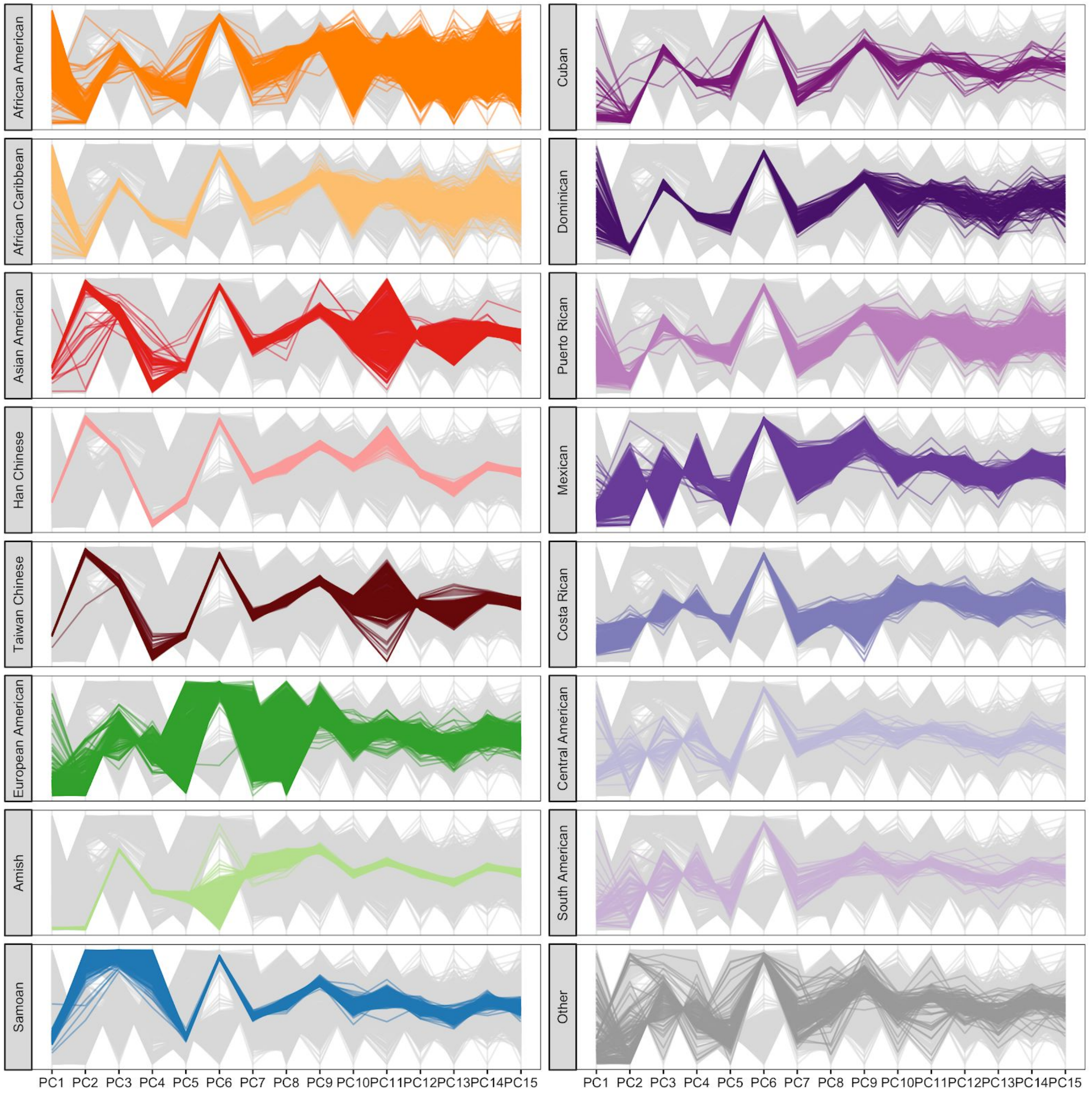
African American
African Caribbean
Asian American
Han Chinese
Taiwan Chinese
European American
Amish
Samoan
Cuban
Dominican
Puerto Rican
Mexican
Costa Rican
Central American
South American
Other
PC1 PC2 PC3 PC4 PC5 PC6 PC7 PC8 PC9 PC10 PC11 PC12 PC13 PC14 PC15

**Supplementary Figure 24. Parallel coordinates plots for the first 15 principal components of Freeze 5 genotype data pooled across studies.** All panels contain the same set of lines, but each panel highlights a single category reflecting race, ancestry, and/or ethnic information provided by the participants or specified by study inclusion criteria.

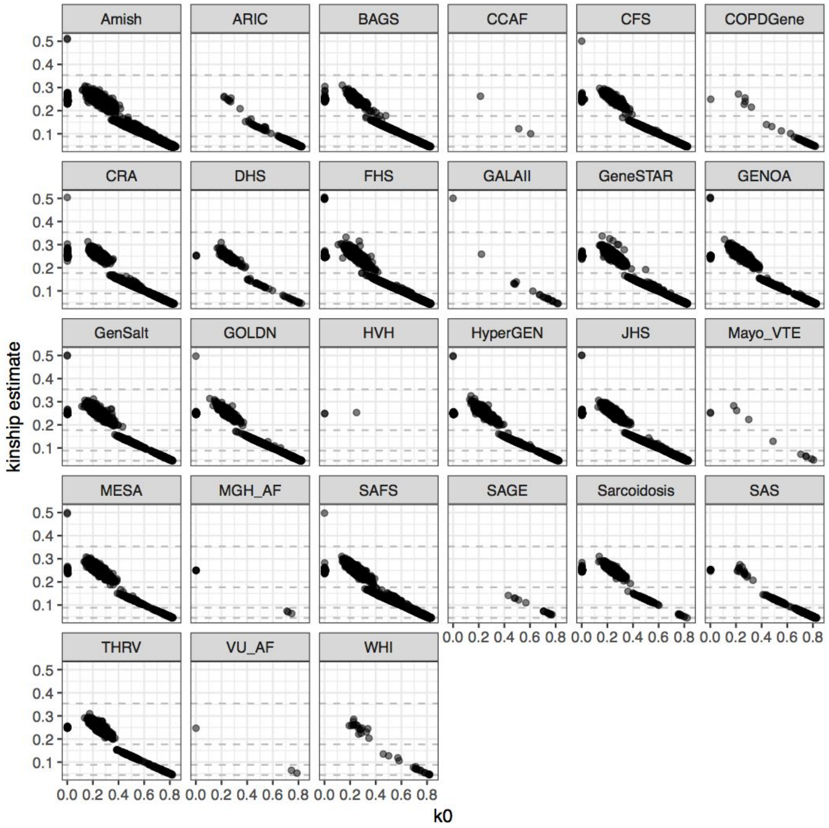


**Supplementary Figure 25. Relatedness of subjects within each study.** The y-axis shows the kinship coefficient (KC) estimated by PC-Relate. The kinship coefficient for a pair of participants is KC = k2/2 + k1/4, where k2 is the probability that two pairs of alleles are identical by descent (IBD) and k1 is the probability that one pair of alleles is IBD. The x-axis shows k0, the probability that zero alleles are identical by descent. Each point represents a pair of samples. Gray dashed horizontal lines show boundaries for KC values for inferring varying degrees of relatedness. Moving from the top down, monozygotic twins are in the top left corner, the first and second dashed lines form a region for expected first-degree relatives (parent-offspring and full siblings), the second and third form a region for expected second-degree relatives, the third and fourth for expected third-degree relatives, and below the fourth we expect unrelated or related at fourth or higher degree. See Supplementary Table 2 for study abbreviations.

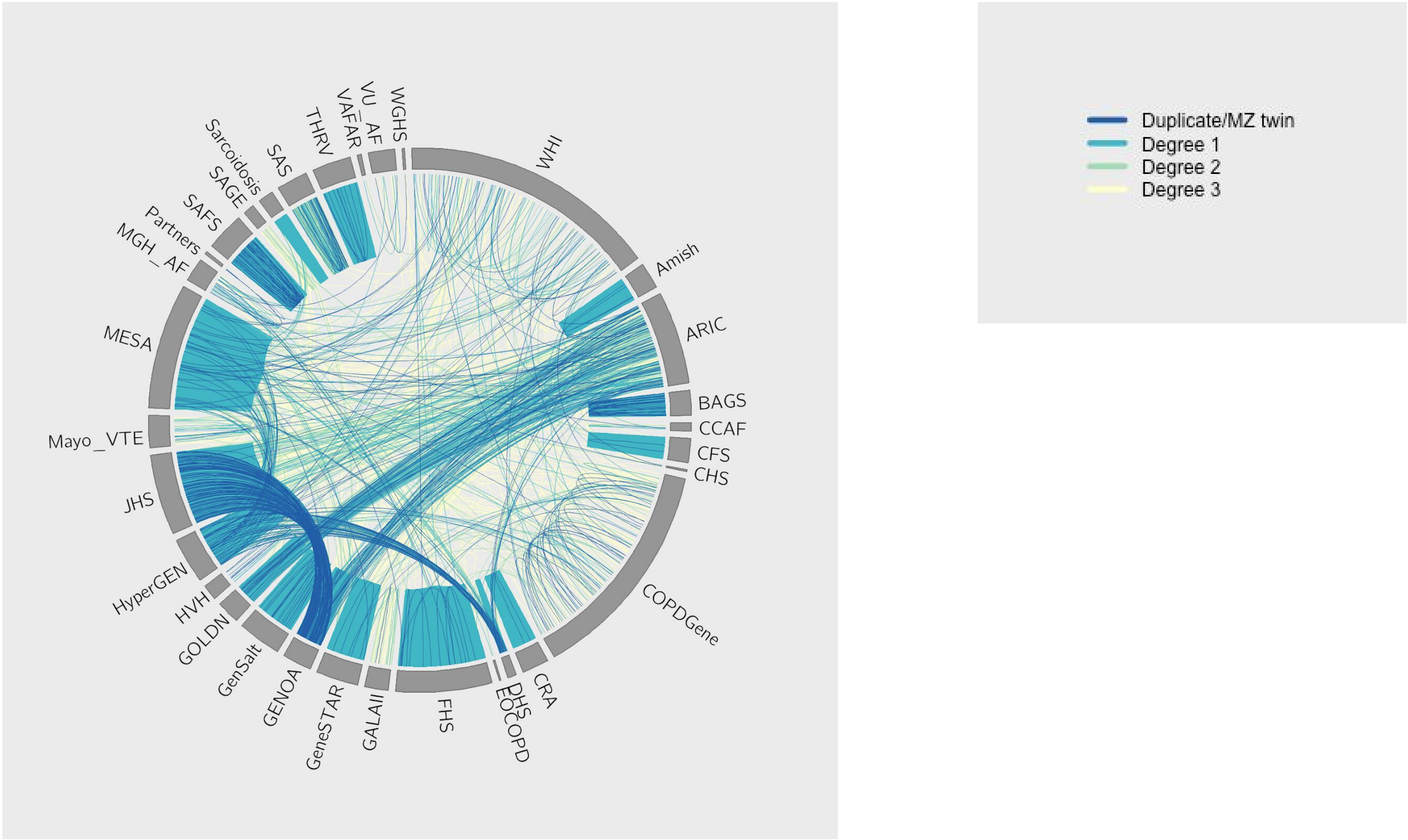


**Supplementary Figure 26. Relatedness of subjects within and across studies.** Line color indicates degree of relationship: blue=duplicates or monozygotic twins, blue-green=first-degree relatives (parent-offspring and full siblings), green=second-degree relatives, yellow=third-degree relatives. See Supplementary Table 2 for study abbreviations.

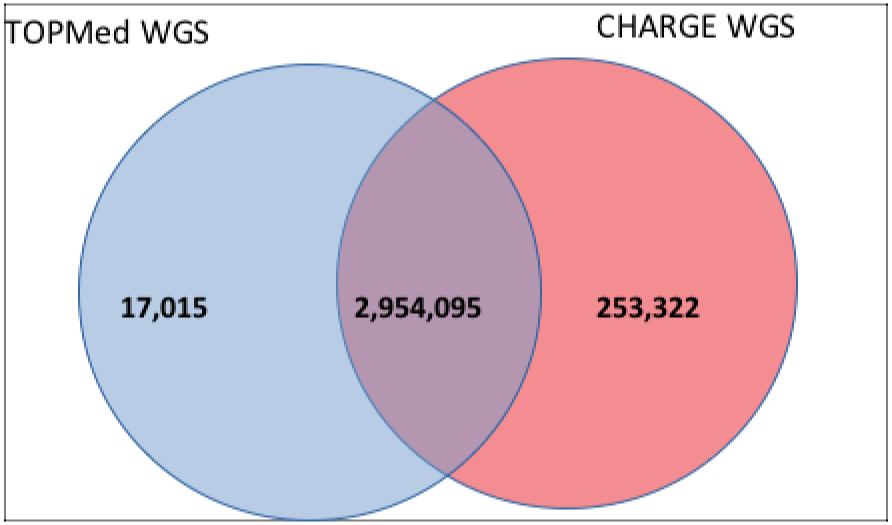


**Supplementary Figure 27. Common variants overlap between TOPMed WGS and CHARGE WGS.**

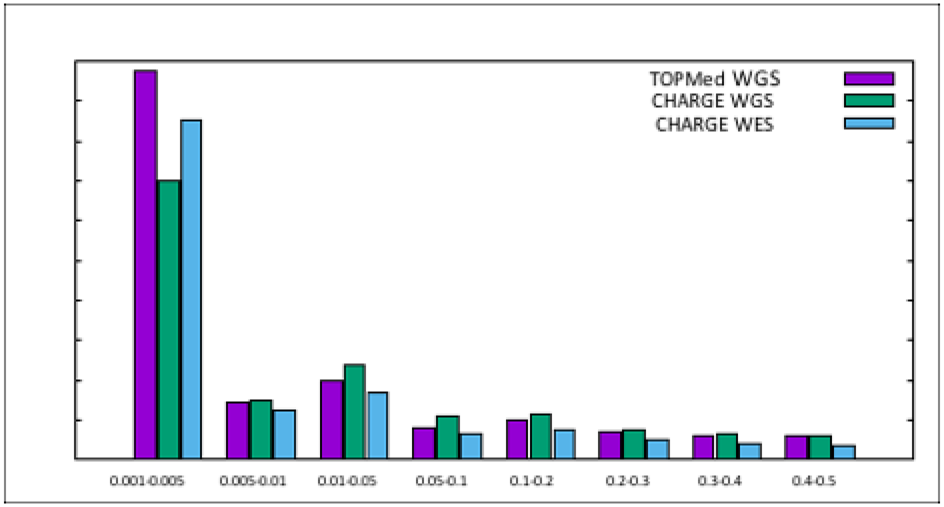


**Supplementary Figure 28. Exonic bi-allelic SNV counts by minor allele frequency.**

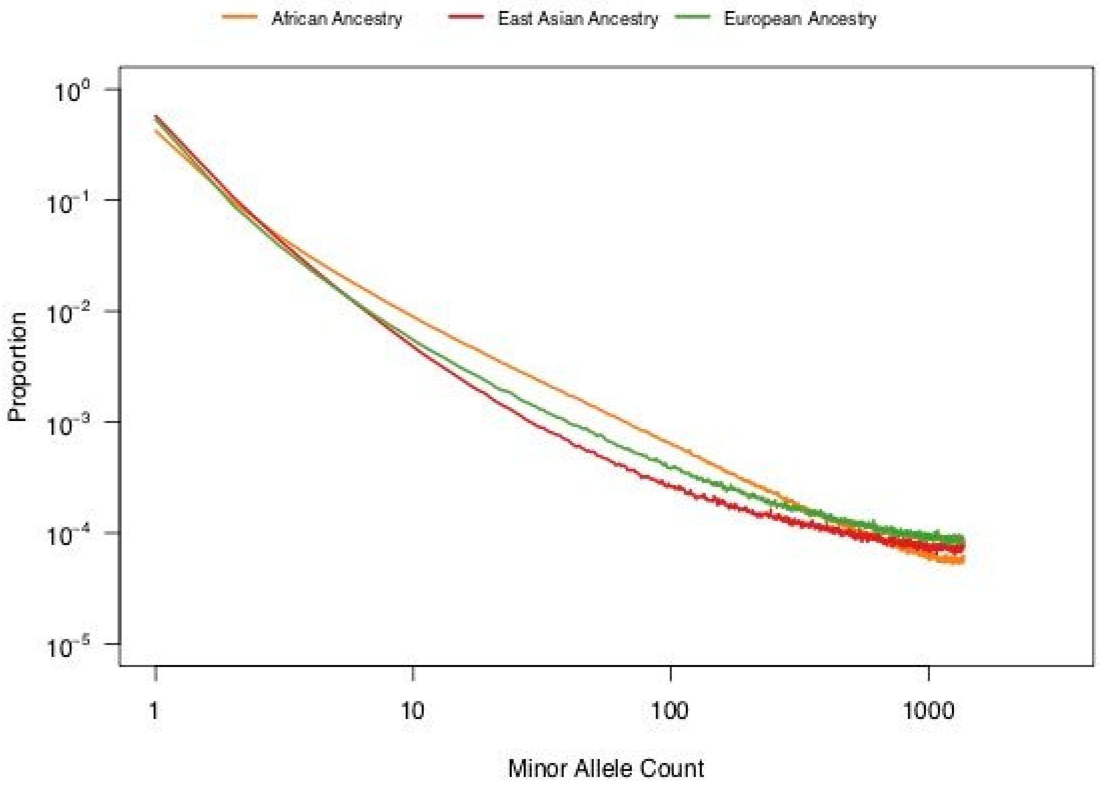


**Supplementary Figure 29. Site Frequency Spectrum (SFS) across three major populations.** This log-log histogram of the SFS is based on 1,370 unrelated individuals per population. In all three populations we see a shift towards extremely rare variation consistent with exponential growth in the last 5000-10000 years. We see a reduction in common variants for East Asian individuals consistent with a protracted bottleneck.

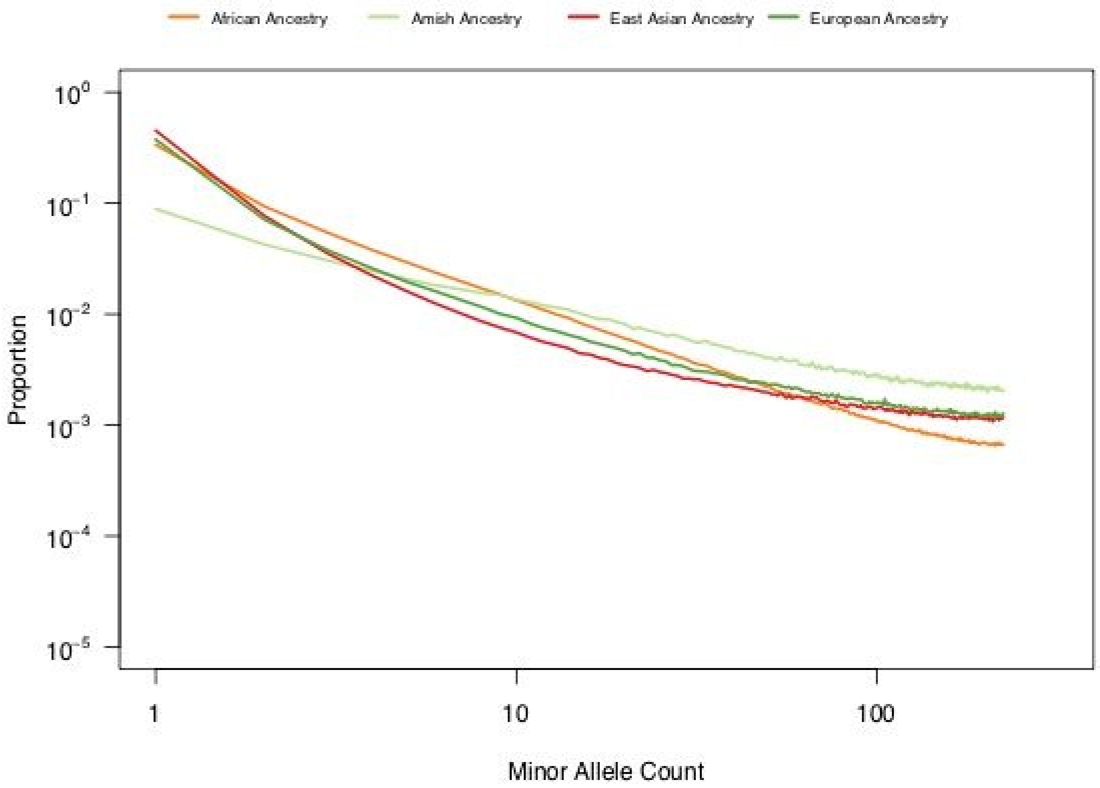


**Supplementary Figure 30.** This log-log histogram of the SFS is based on 225 unrelated individuals per population. While all four populations show a shift towards extremely rare variation consistent with exponential growth in the last 5000-10000 years, this pattern is notably less pronounced in the Amish, which is consistent with the Amish experiencing a very recent bottleneck.

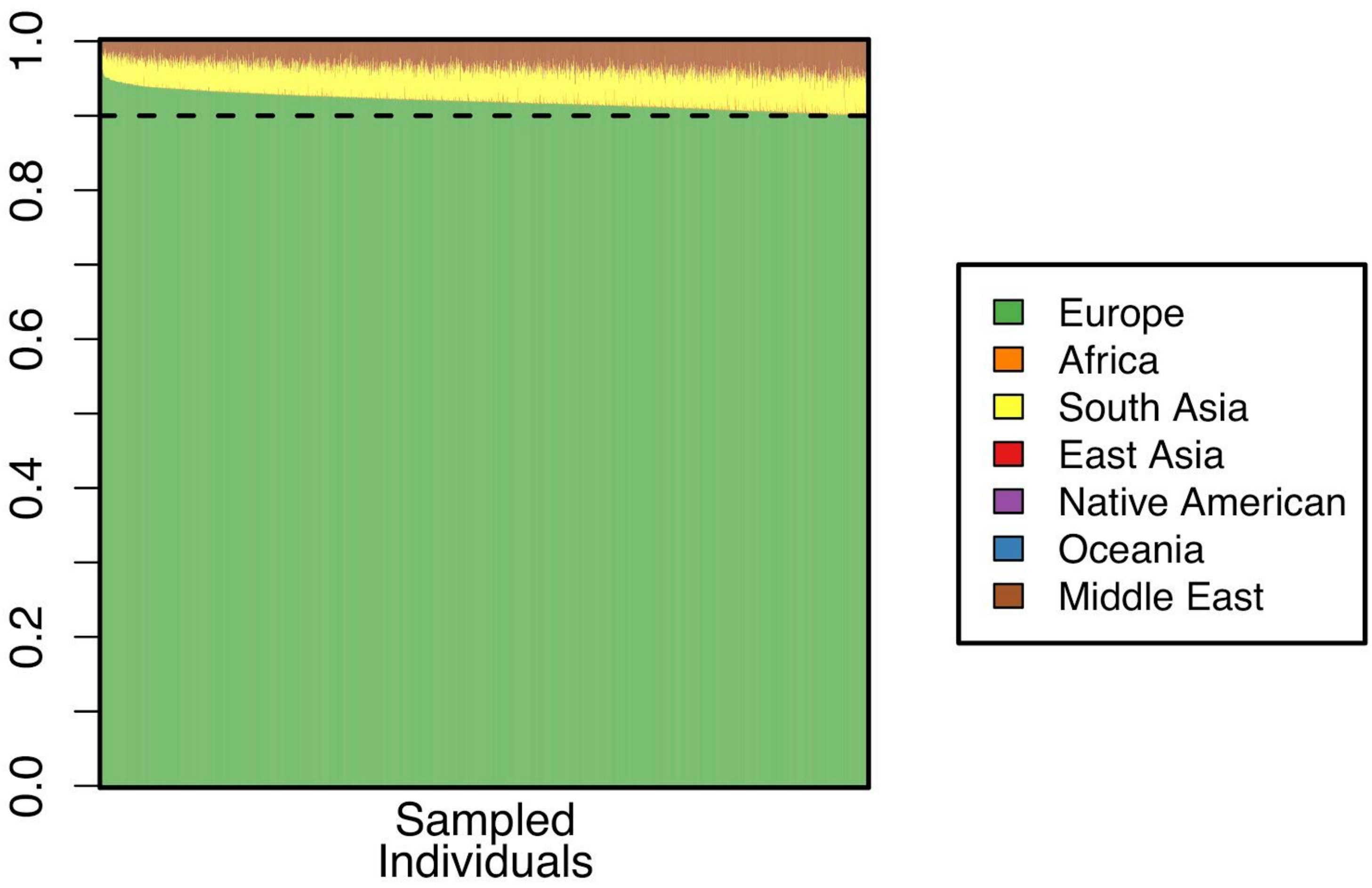


**Supplementary Figure 31. Global ancestry of 2,416 European individuals (inferred by RFMix) used for demographic inference.** All 2,416 individuals have 90% or greater European ancestry (represented by dashed line).

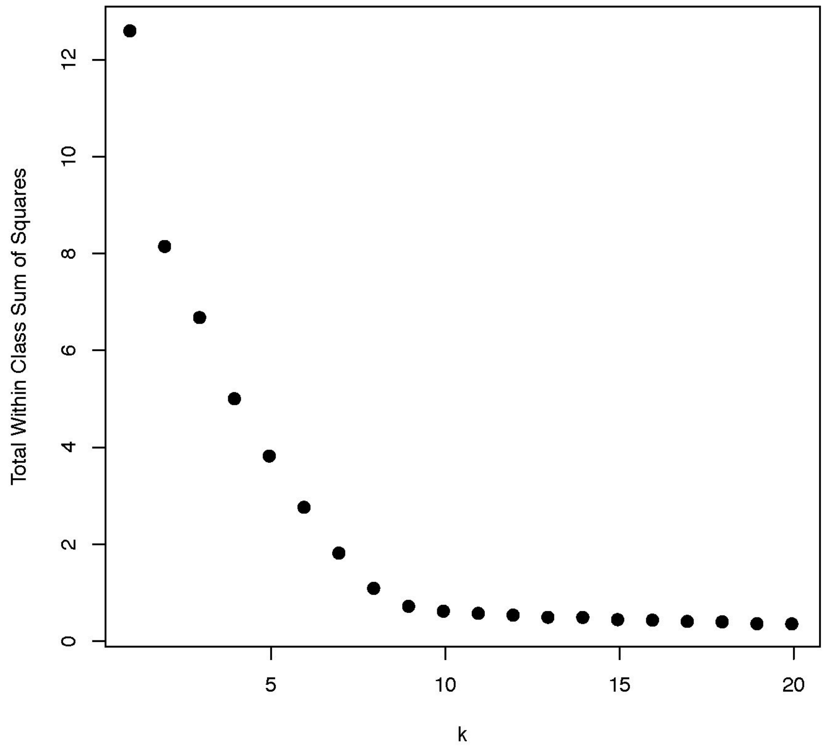


**Supplemental Figure 32**. Within class sum of squares versus number of clusters (k) for k-means clustering of PCA coordinates.
